# Fluoroscopic-Guided Magnetic Soft Millirobot for Atraumatic Endovascular Drug Delivery

**DOI:** 10.1101/2025.02.17.637470

**Authors:** Tuan-Anh Le, Husnu Halid Alabay, Prabh G. Singh, Melissa Gioia Austin, Sanjay Misra, Hakan Ceylan

## Abstract

Atraumatic and precise drug delivery to arteries and veins remains an unmet need in interventional medicine, with significant implications for managing vascular diseases and long-term patient outcomes. Conventional endovascular methods, such as drug-coated balloons and drug-eluting stents, often damage vessel endothelium and compromise wall integrity, leading to reduced therapeutic efficacy and severe complications including restenosis and thrombosis. To address these limitations, we introduce EndoBot, an untethered soft millirobot designed for atraumatic vessel navigation and localized drug delivery under physiological blood flow. EndoBot achieves this through magnetically actuated corkscrew propulsion and mechanically adaptive surface crawling, exerting low radial pressure (< 1 kPa) to preserve endothelial integrity. We validated its performance in phantom vessels, ex vivo human umbilical veins under normothermic perfusion, and in vivo rat inferior vena cava under fluoroscopic guidance using a human-scale magnetic manipulation platform. Despite dynamic vessel contractions and compressions along non-uniform cross-sections, mechanically adaptive locomotion strategy ensures safe navigation and protects the inner endothelial lining. EndoBot is deployable and retrievable via clinical vascular sheaths and remains stable even under supra-physiological blood flow conditions (> 155 cm/s). Extensive blood compatibility tests demonstrated minimal hemolysis (<0.01%), low platelet activation on the robot surface, and no increased coagulation tendency. For drug delivery, a hydrophobic transfer coating enables EndoBot to gently deposit a stable, flow-resistant drug layer onto the vessel wall without fragmentation. This novel platform enables safer and more effective targeted endovascular drug delivery, paving the way for transformative approaches to address early-stage vascular pathologies and deliver more effective preventive interventions.

## INTRODUCTION

Vascular diseases remain one of the leading causes of morbidity and mortality worldwide, accounting for over 18 million deaths annually and placing an immense burden on global healthcare systems(*1–3*). Each year, millions of patients undergo minimally invasive endovascular procedures to restore vascular perfusion and manage inflammatory complications such as intimal growth, stenosis and thrombosis(*4*). Traditionally, these procedures rely on catheter-directed approaches, drug-eluting stents (DESs) and drug-coated balloons (DCBs) to mechanically deliver therapeutics into the vessel wall(*5–7*).

Despite their widespread clinical use, the efficacy of these endovascular technologies for localized drug delivery is fundamentally constrained by safety limitations. Catheter-directed drug delivery often demands high dosing to offset systemic dilution, leading to off-target toxicity and potential organ damage, including the kidneys and liver(*8*). While DESs and DCBs reduce systemic drug exposure, they require high radial pressures on the order of ∼800-1000 kPa during angioplasty, far exceeding the physiological tolerance (∼12.4 kPa) of endothelial cells (ECs) that line the vessel lumen(*9–13*).

ECs play a vital role in modulating immune responses, maintaining hemostasis, regulating vascular tone, and preventing excessive smooth muscle cell proliferation(*8, 14–17*). Consequently, damage to ECs and delayed reendothelialization are strongly associated with neointimal hyperplasia, restenosis, thrombosis, and long-term vascular dysfunction(*18, 19*). To mitigate these adverse events, current practices often require prolonged dual antiplatelet therapy (6–12 months) following DES implantation or balloon deployment. However, this extended therapy not only increases risk of bleeding complications but also delays natural healing process at the damage site, compounding the challenges of patient recovery(*18, 20*).

Given the pivotal role of ECs in maintaining vascular health, the endothelium stands out as a critical therapeutic target(*21*). Yet, despite decades of innovation, no device-based platform has been developed that can deliver drugs directly onto a discrete segment of the vascular endothelium while preserving its structural and functional integrity. Although nanocarrier-based approaches have been explored for targeted deposition, they often face challenges with spatial localization and systemic toxicity(*22*). Consequently, there remains a pressing need for a technology capable of gentle, mechanically adaptive interaction with the vessel lumen to enable safe and precise drug delivery.

Small-scale robots can offer a new avenue to address these safety challenges inherent to conventional vascular interventions. Magnetically steerable continuum robots have recently demonstrated deeper and more precise vascular access. However, they still rely on tethered catheters and guidewires, whose friction against highly curved, three-dimensional (3D) vessel segments can induce vasoconstriction, EC denudation and vessel wall injury(*23, 24*). This friction elevates the risk of vessel perforation, embolization, and subsequent ischemia. By contrast, magnetically steered milli- and microrobots that provide medical image-guided untethered mobility, mechanical compliance and compatibility with blood flow environments could overcome these limitations. Indeed, various rigid and soft-bodied small-scale robots have been proposed for proof-of-concept endovascular applications—including targeted drug delivery, clot removal, and embolization(*25–33*). Nevertheless, these demonstrations have largely remained confined to in vitro or ex vivo cadaveric models, offering only limited insights into the operability within complex physiological environments of living systems. Also, many face challenges in systematically addressing design strategies and elaborate on well-defined safety considerations when interacting with live vasculature. This gap limits their translational potential for a real-world application. To our knowledge, no untethered miniature robot has yet been demonstrated for medical image-guided navigation and endovascular drug delivery in a live animal using a human-scale magnetic navigation system. Although sophisticated cargo-release mechanisms have been explored, a fully atraumatic, localized approach to delivering therapeutics to the vessel wall remains unproven, underscoring a significant unmet clinical need in the management of vascular diseases(*29, 34–36*).

To address this grand challenge, we introduce EndoBot, an endoluminal soft millirobot designed for atraumatic, localized drug delivery in small- to mid-sized vessels with diameters of 1-4 mm. This diameter range encompasses a diverse set of clinically significant small- to mid-sized vessels, including many coronary branches, peripheral arteries, and superficial veins commonly targeted for minimally invasive interventions **(Table S1)**. EndoBot navigates via magnetic actuation under fluoroscopic guidance within a human-scale workspace **(Fig 1A)**. It features two main components: a soft, elastic robot backbone and a hydrophobic, lubricant drug-transfer coating. The backbone is formed as a thin, magnetic, and hollow helix, drawing inspiration from biological locomotors, such as bacterial flagella(*37*), self-burying plant seeds(*38*), and miniature robot designs proposed for swimming(*29*) and surface crawling(*32*) (**Fig 1B**).

**Figure 1.**
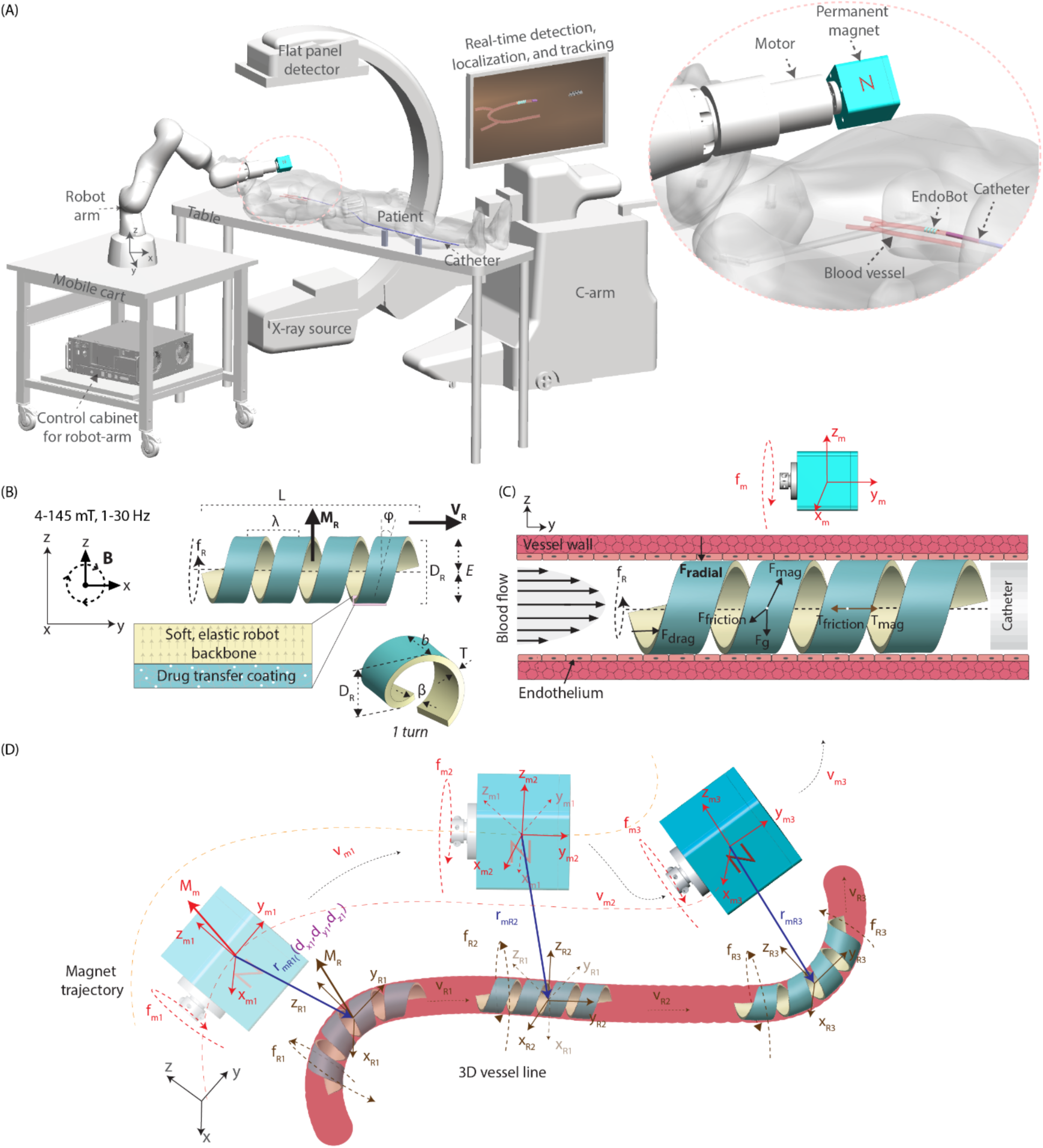
Clinical vision for the EndoBot-mediated endovascular drug delivery system. (A) X-ray fluoroscopy–guided magnetic microrobot manipulation system, integrated with real-time detection, tracking, localization, and steering control, designed for human-scale interventions. (B) Structural design features and main components of EndoBot, including its soft, elastic soft robot backbone, and hydrophobic, flow resistant drug transfer coating. (C) The open helical structure of EndoBot, which offers high radial deformability, enables mechanically adaptive crawling on the vessel surface and corkscrew locomotion under physiological blood flow without injuring inner lining endothelial cell layer and the vessel wall. (D) Six degrees-of-freedom magnetic actuation, propulsion, and steering are achieved through external control of magnetic torque and force, ensuring flow stability, precise navigation, and effective drug delivery.

To enable atraumatic and effective navigation and drug delivery in complex vascular networks, a key design feature of the backbone is its adaptive crawling mechanism, which maintains gentle contact with the vessel lumen to ensure unobstructed blood flow and robust locomotion. The body undergoes elastic and conformal deformation along curved and non-uniform cross-sections, enabling it to adapt to pulse waves and vasospasms in vivo while applying minimal radial forces to preserve endothelial integrity **(Fig 1C-D).** Meanwhile, as EndoBot moves, its hydrophobic coating is transferred to the vessel surface, forming a stable, flow-resistant drug-depot layer without causing downstream fragmentation. By combining adaptive crawling with a novel, wiping-based transfer method, EndoBot achieves safe, efficient, and sustained endoluminal drug delivery capability.

In this study, we validated the EndoBot drug delivery platform in phantom vessels, normothermically perfused human umbilical veins ex vivo, and in vivo rat inferior vena cava. These experiments demonstrated the feasibility under realistic physiological and anatomical conditions within human-scale operational constraints and established safety limits. Extensive performance tests and comprehensive design of the platform provide a strong basis for its translational potential. Future efforts will focus on refining EndoBot and evaluating its capabilities in both small and large animal models, particularly for preventive interventions targeting vascular endothelium inflammation and vessel walls over extended time points. Ultimately, EndoBot could transform vascular disease management by shifting the paradigm from treatment to prevention, offering safer, more effective, and minimally invasive solutions that may surpass current gold standards.

## RESULTS

### EndoBot fabrication and design refinement

The EndoBot backbone was fabricated from a polydimethylsiloxane (PDMS)-based composite (Sylgard™ 184, Dow Inc.) embedded with NdFeB powder (D50, 5 μm, Magquench GmbH). This magnetic elastomer composite is highly reproducible and offers tailorable elasticity and magnetic responsiveness, ensuring robust performance under various operational conditions. Additionally, the industrial-scale availability of these materials aligns with FDA requirements for consistent, scalable manufacturing and regulatory compliance, bolstering translational feasibility.

To systematically refine the structural and compositional features of EndoBot for nominal robot diameters (*D_R_*) ranging from 1.0 to 4.0 mm we established a high-fidelity soft lithographic fabrication method **(Supplementary Text 1, Figs S1-3)**. *D_R_* is chosen according to the vessel segment being targeted, while other key helical parameters, including length (*L*), helical pitch length (λ), helical blade thickness (*b*), groove size (*β*) and thickness (*T*) are scaled relative to *D_R_* using constant coefficients **(Fig 1B, Table S2)**. Certain parameters, such as helical angle (φ) is not scaled to *D_R_* and remained constant. These relationships were established through rational design requirements, empirical measurements, and earlier studies of optimized corkscrew microswimmers designed for low Reynolds number environments.

While the scaling coefficients originate from the as-fabricated dimensions, some parameters (*D_R_*, *L*, and λ), dynamically adjust relative to one another under vascular confinement and magnetic propulsion. For example, maintaining an *L*/*D_R_* ratio greater than 3.0 enables more efficient magnetic torque application along the major helical axis, reducing the risk of tumbling under off-axis loads(*29, 39, 40*). In this study, EndoBot was constructed with an initial *L*/*D_R_*ratio of 3.3, which dynamically ranged from 3.3 to 7.0 during navigation due to partial elongation in the axial dimension and compression in the radial dimension. This dynamic adaptability works in favor of maintaining effective torque application and robust corkscrew propulsion even in narrower vessel segments.

The *L*/λ ratio, representing the number of helical turns, reflects the ability of the helix to act as a mechanical dampener under radial compression. The as-fabricated EndoBot has an initial *L*/λ ratio of 4.8, which dynamically adjusts between 4.8 and 6.7 during navigation across smaller vessels than *D_R_*. This adaptability allows the structure to distribute stress along its length, absorbing and storing elastic energy without forming localized high-strain points that could lead to buckling or protrusions into the blood vessel. By creating new, conformal turns under compression, the helix maintains structural integrity, enabling smooth navigation while preserving vascular patency and ensuring vessel wall safety.

To impose a high mechanical cost on erratic deformations and maintain controlled adaptations during motion, *T* was identified as a strategic variable for tuning deformation behavior of the helix **(Supplementary Text 2)**. The effective elastic modulus (*E*) across the cross-section scales nearly cubically with *T* (*E* ∝ *T*^2.54^), so even moderate increases in *T* substantially raise intrinsic radial force (**F_intrinsic_**) arising from stored elastic energy under compressive strain (**F_intrinsic_** ∝ *T*^2.54^). At the same time, increasing *T* reduces both the conformal deformation capacity (*C*) under magnetic actuation (*T*∝ 1/*C*) and vascular patency (*S*) (*T* ∝ 1/*S*) **(Fig S4-8)**. However, increasing *T* also raises radial force **(Figs S4,6)**. Consequently, *T* was judiciously optimized to balance robust structural integrity and mechanical compliance for adaptive crawling, while minimizing radial pressure on the vessel wall and preserving vessel patency.

In addition to refining *T*, we iterated the NdFeB mass ratio within the PDMS matrix and adjusted curing conditions of the elastomer to fine-tune *E* and enhance magnetic responsiveness without compromising vascular patency **(Figs S4-8, Tables S3-6)**.

For drug transfer efficiency, the contact area between EndoBot and the vessel wall was maximized by employing a high *b/β* ratio while minimizing φ **(Fig S9A-C)**. A smaller φ not only increases wall contact but also reduces the cross-sectional area occupied by EndoBot, thereby improving overall vessel patency **(Fig S9D)**. Guided by these design priorities and comprehensive refinements, we have established a scalable blueprint for validation in diverse vascular and flow conditions **(Table S2)**.

### Establishing operational limits for endovascular navigation performance

When planning EndoBot deployment in a specific vessel, it is essential to evaluate the minimum (*D_min_*) and maximum (*D_max_*) luminal diameters of the target segment. *D*_R_ must be chosen to ensure sufficient lumen contact for consistent surface crawling, while still accommodating diameter variations **(Fig 2A)**. However, when EndoBot with *D_R_* = 3.6 *mm* (hereafter referred to as 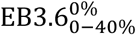, following the nomenclature in **Fig 2B**)) was subjected to cyclic deformation up to 40% compressive strain (ε), it recovered approximately 94% of its original diameter after the first loading/unloading cycle (**Fig 2C, Supplementary Movie 1)**. On subsequent cycles, the diameter stabilized at this slightly reduced value, indicating a mild hysteresis effect during the first compression. The recovery rates following the initial load suggest that the hysteresis effect is more pronounced at ε < 10% and it diminishes with increased strain **(Fig S10)**. To alleviate this effect, we introduced a bias compressive strain (*ε_Bias_*) as a pre-applied minimum strain throughout navigation. Practically, ε_Bias_ is achieved by selecting D_R_ to be larger than D_max_, so that sufficient compressive strain is consistently maintained. Indeed, 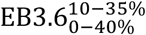 reproducibly recovered its biased diameter after each loading-unloading cycle under 10-40% compression **(Fig 2D)**. Based on these observations, we define Rule 1: EndoBot must always maintain at least 10% bias compressive strain, which is equivalent to *D_R_* ≥ 1.1× *D_max_*.

**Figure 2.**
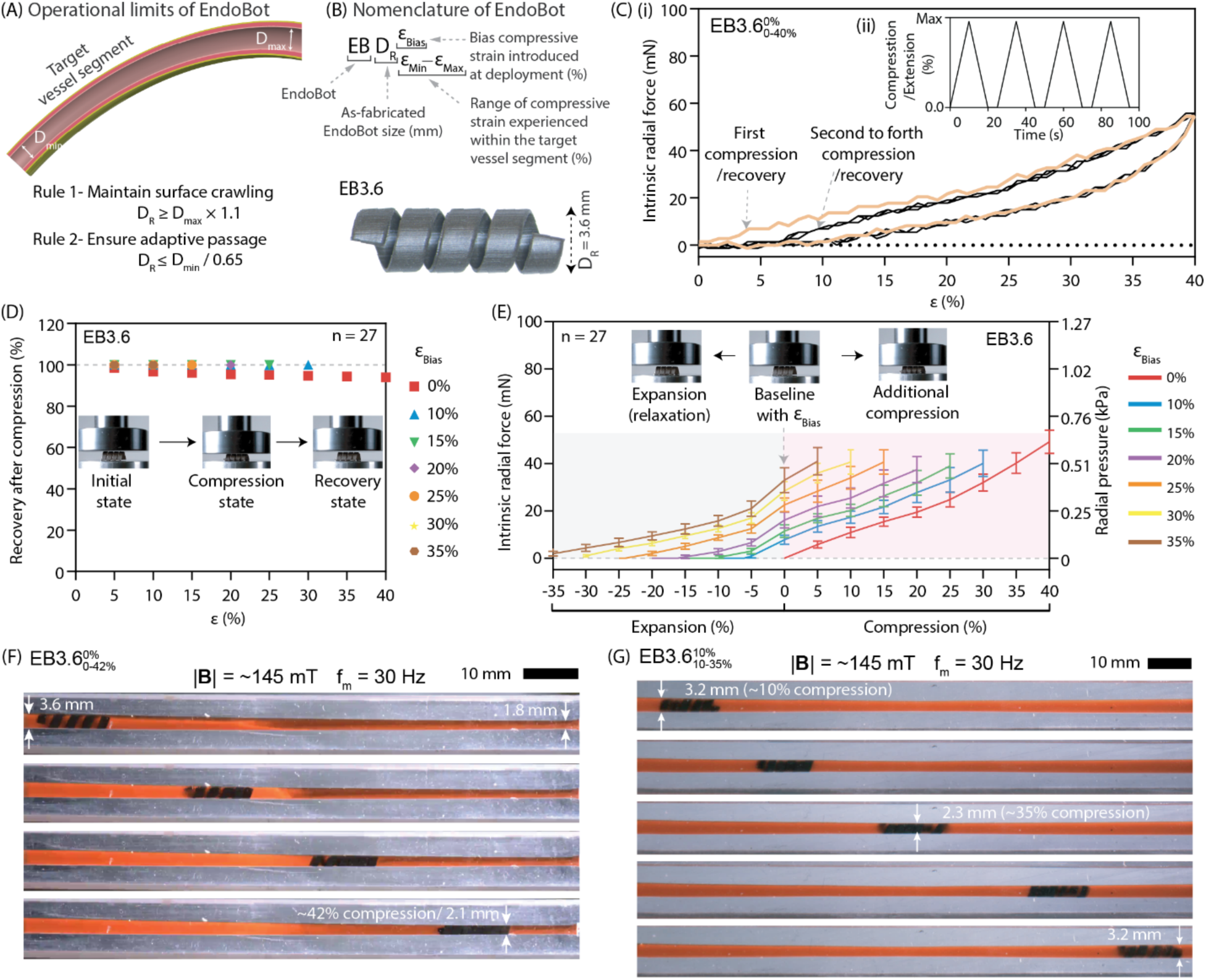
Operational safety limits of EndoBot. (A) EndoBot selection principles. Two key rules govern the appropriate selection of EndoBot for a given target vessel: Rule 1: EndoBot must maintain a minimum bias compressive strain of 10%, ensuring its effective diameter (D_R_) is at least 1.1× the maximum vessel diameter (D_max_). Rule 2: EndoBot must operate under a maximum strain of 35%, meaning D_R_ should not exceed 65% of the minimum vessel diameter (D_min_). (B) Nomenclature and design features of EndoBot. “EB” represents EndoBot and “D_R_” (in mm) is its as-fabricated diameter. The superscript “ε_Bias_” indicates the bias compressive strain applied at deployment, while the subscript “ε_Min_–ε_Max_” specifies the range of compressive strain experienced during operation or mechanical testing. (C) Intrinsic radial force and cyclic deformation. (i) Measurement of the intrinsic radial force generated by EB3.6 when subjected to cyclic deformations up to 40%. (ii) Time-course profile of the cyclic deformation pattern. (D) Diameter recovery post-compression. Recovery of the EB3.6 diameter following 40% compressive strain by introducing a bias compression. (E) Intrinsic radial force and radial pressure of EB3.6 during compression and recovery. (F) Conformal deformation capacity measured in a tapered vessel. 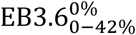 under rotating magnetic field in a blood-filled conical phantom vessel with diameters tapering from 3.6 mm to 1.8 mm. (G) Adaptive locomotion in a non-uniform vessel. Testing of 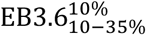 in a phantom vessel with diameters ranging from 3.2 mm to 2.3 mm under a rotating magnetic field, confirming safe and adaptive crawling locomotion within its operational safety limits.

Moreover, if the vessel has wider cross-sections along its length, a higher bias strain can be pre-applied to store additional elastic energy, enabling EndoBot to expand and maintain contact even in larger-diameter regions **(Fig 2E)**. For instance, with a 30% bias compression, EB3.6 can safely recover almost all its compressed diameter (> 28%), providing a robust operational margin that prevents loss of contact in broader segments while still allowing safe navigation through narrower portions. Notably, even at 40% compression, the radial pressure generated by EndoBot is below 0.75 kPa, which is over an order of magnitude smaller than the endothelial rupture threshold (∼12.4 kPa), suggesting atraumatic interactions with the vessel wall across the spectrum of the robot deformation(*12, 13*).

Next, to evaluate the conformal deformation capacity of EndoBot, we deployed 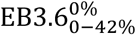 in a blood-filled conical phantom vessel of which diameter tapers from 3.6 mm to 1.8 mm **(Fig 2F, Supplementary text 3, Table S7, Fig S11)**. Although 50% diameter variation exceeds what might be observed in early or moderate disease states, it provides a single, continuous model to systematically test the upper deformation limits of EndoBot. Under these conditions, 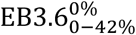 navigated successfully at up to 42% compression **(Supplementary Movie 1)**. Beyond that threshold, however, the constant magnetic torque could not overcome the combined resistive effects: (i) stored elastic energy increasing with *ε* (U_elastic_ ∝ *ε*^2^); (ii) amplified normal force (|**F_radial_|**∝ *ε*) (**Supplementary text 2)**, which raises friction at the EndoBot–vessel interface (|**F_friction_|** ∝ |**F_radial_|**) **(Fig S12, Supplementary text 4)**. To prevent this immobility and ensure safe, repeatable locomotion, we incorporated a 7% safety margin and established Rule 2: EndoBot must operate with a maximum *ε* of 35%, or equivalently *D_R_* ≤ *D_min_* /0.65. For example, if the vessel segment has D_max_ = 3.2 mm and D_min_ = 2.3 mm (**Table S7)**, then selecting *D_R_* = 3.6 mm meets both Rule 1 and Rule 2, not risking prohibitive deformation while retaining sufficient surface contact. Indeed, testing EB3.6 in a blood-filled phantom vessel in this diameter range further verified safe and adaptive crawling locomotion (**Fig 2G**, **Supplementary Movie 1**).

### Stability and propulsion of EndoBot under physiological and supraphysiological blood flow

A safe EndoBot design must remain stationary without drifting and must avoid obstructing blood flow at physiological velocities typically encountered in small-to medium-sized arteries and veins, with lumen diameters ranging from 1.0 to 3.0 mm **(Table S1)**. This stability is primarily achieved by balancing static friction at the robot-lumen interface with the drag forces (**Supplementary text 5, Fig S13**) exerted by blood flow at the robot’s surface. For stability, static frictional forces must exceed the drag forces to prevent the Endobot from drifting. To evaluate these dynamics, we measured the maximum blood flow rates and velocities that caused 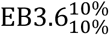 and 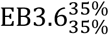 to drift in phantom vessels. This was achieved by developing a system that mimics the anatomical and physiological conditions of blood vessels (**Fig 3A**, **Supplementary text 6** and **Fig S14**).

**Figure 3.**
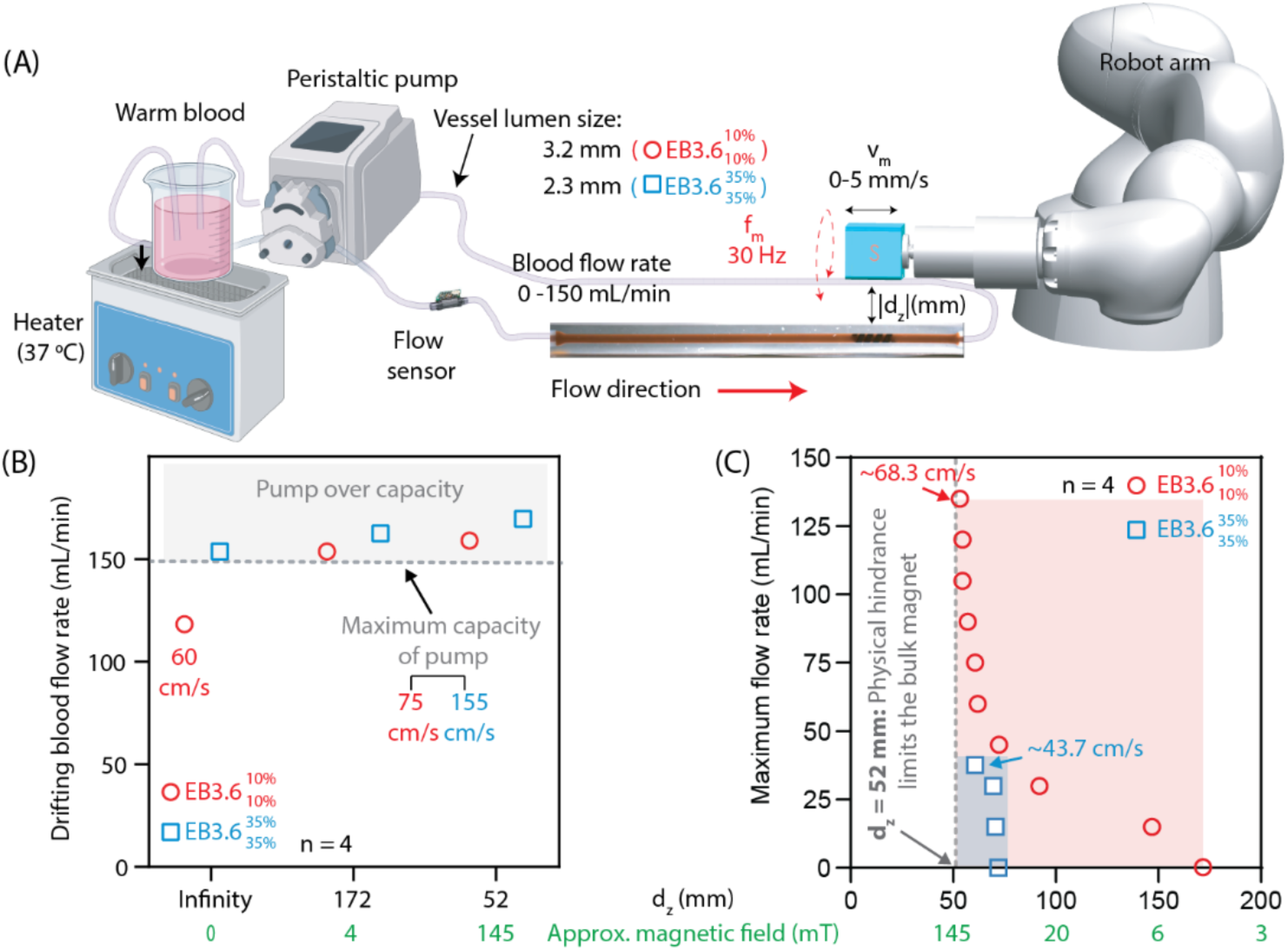
Stability of EndoBot in physiological blood flow. (A) Description of experimental conditions and the design of the testbed. (B) Maximum flow rate and flow velocity that 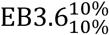 and 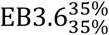 can withstand before drifting with the flow. The indeterminate area represents flow rates beyond the capacity of the peristaltic pump, designated as the highest flow rate and velocity causing drift. |**B**| = 0-145 mT, f_m_ = 0 Hz, |**v**_m_| = 0 mm/s. (C) Maximum flow rates at which 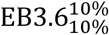 and 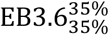 could be safely navigated with and against the flow under the minimum external magnetic field required for motion. |**B**| = 0-145 mT, f_m_ = 30 Hz, |**v**_m_| = 0-5 mm/s.

Both 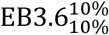 and 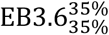 demonstrated remarkable stability, tolerating flow rates up to 120 mL/min (60 cm/s) and 150 mL/min (155 cm/s), respectively, without the assistance of external magnetic fields **(Fig 3B)**. When an external magnetic field as low as 4 mT was applied, the stability exceeded the maximum flow rate capacity of our pump (150 mL/min) for both designs due to the appearance of **F_extrinsic_** friction caused by the magnetic pulling force **F**_mag_ (**Supplementary texts 2 and 7, Fig S15-S19**). These flow rates correspond to physiological conditions observed in arteries and veins such as the middle cerebral artery, coronary artery, brachial artery, radial vein, and cephalic vein. Human blood flow under anesthesia in these vessels typically ranges between 5 and 150 mL/min, depending on vessel size and hemodynamic state **(Table S1)**. To assess stability under more challenging conditions, we introduced random pinches and folds into the tube a few centimeters upstream of the robot to create irregular and turbulent flow patterns. Despite these disruptions, both EndoBot maintained their stability, demonstrating their safety for deploying in vascular environments with diverse flow dynamics.

To further explore the stability dynamics of EndoBot under kinetic friction, we applied a constant rotational frequency of 30 Hz while maintaining a propulsion optimization to achieve highest locomotion performance for EndoBot (**Supplementary text 7)**. We then determined the maximum flow rates at which 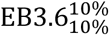 and 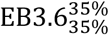 could be safely navigated with and against the flow under the minimum external magnetic field required for motion. In this assessment, while 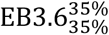 demonstrated slightly superior stability under static friction conditions due to increased radial compression, significant differences emerged under kinetic friction, with 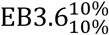 exhibiting substantially better performance **(Fig 3C)**.

Under an external magnetic field of approximately 145 mT, where both designs achieved their best actuation performance, 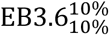 maintained navigational stability up to a flow rate of 135 mL/min (68.3 cm/s), whereas 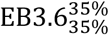 was limited to a flow rate of 38 mL/min (43.7 cm/s). This disparity is primarily attributed to the interplay between frictional and drag forces. For 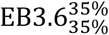, increased radial compression enhanced frictional forces |***F***_*friction*_| ∝ ε_Bias_ (**Supplementary Texts 2,4)** improving static stability. However, this came at the expense of increased drag forces 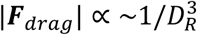 (**Supplementary text 5**) due to the narrowing vessel diameter, which became the dominant limiting factor for upstream navigation. The heightened drag forces necessitated significantly higher magnetic field strength to maintain control, thereby reducing the efficiency and stability of 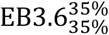 compared to 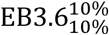.

Based on the superior stability demonstrated under both static and kinetic friction conditions, we selected ε_Bias_ of 10% as the primary design configuration for determining EndoBot size relative to vessel diameter. To further evaluate the magnetic actuation performance of this configuration under systematically controlled multi-physical challenges, including flow, gravity, vessel curvature, and narrowing, we tested 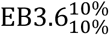 motion in various phantom vessels (**Table S7)** for round trips, both upstream and downstream **(Fig 4, Supplementary Text 7, Supplementary Movie 2)**.

**Figure 4.**
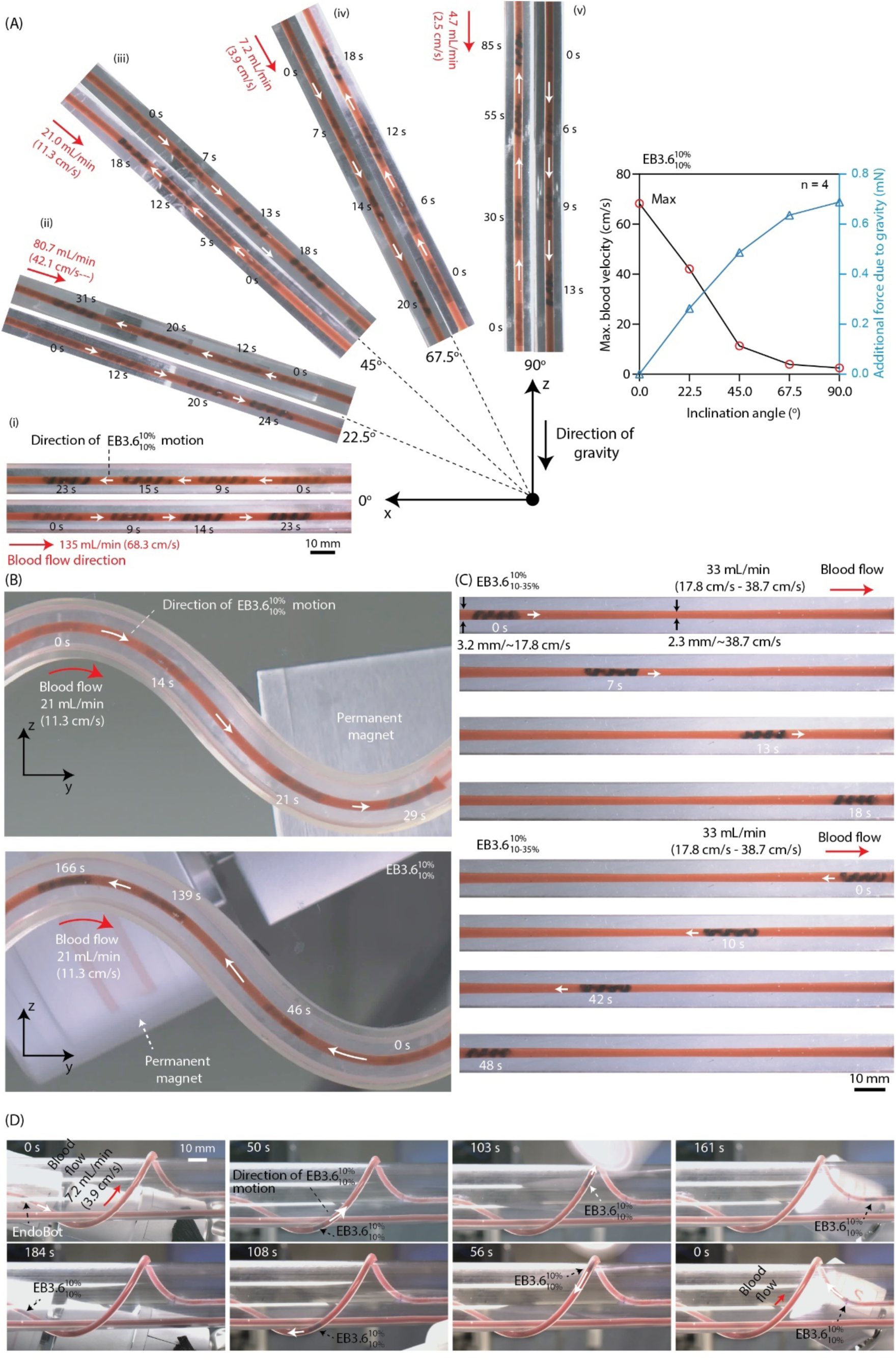
Magnetic propulsion performance of EB3.6 under multi-physical challenges evaluated in high blood flow. (A) The influence of gravity on 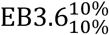 propulsion was assessed in a uniform phantom vessel by varying the inclination of the vessel from 0° to 90°. For each angle, the maximum flow speed at which 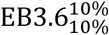 could move both with and against the flow was recorded. (B) The ability of 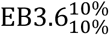 to navigate curved trajectories was evaluated along a sinusoidal phantom vessel. (C) Propulsion performance of 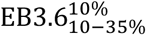 through a non-uniform channel. (D) Navigation control of 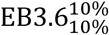 was demonstrated in a human arm-size, arbitrarily shaped three-dimensional phantom vessel under operating conditions of magnetic field strength. |**B**| = 15-145 mT, f_m_ = 30 Hz, |**v**_m_| = 1-5 mm/s.

First, we investigated the influence of gravity on propulsion in a uniform vessel performance by adjusting the inclination angle of straight-shaped phantom vessels from 0° to 90° **(Fig 4A)**. Best upstream motion performance was observed when the vessel was oriented perpendicular to gravity. In this vessel configuration, 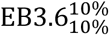 moved against as high as 135 mL/min (68.3 cm/s) bloodstream. As the inclination angle increased, however, gravitational effects degraded upstream control performance. Nevertheless, 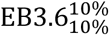 successfully climbed against gravity and a 4.7 mL/min (2.5 cm/s) bloodstream while maintaining precise magnetic control.

Next, we tested propulsion performance along a sinusoidal curve **(Fig 4B)**. 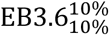 successfully moved against a blood flow rate of 21 mL/min (11.3 cm/s), performing comparably to its motion in a straight vessel inclined at 45° to gravity **(Fig 4Aiii)**. Importantly, the bending of the robot through the curve did not negatively affect its locomotion behavior. We further evaluated propulsion performance through a narrowing channel **(Fig 4C)**. 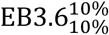 navigated blood flow at 33 mL/min, where the velocity reached 17.8 cm/s at a 3.2 mm diameter segment and 38.7 cm/s at a narrower 2.3 mm segment at the middle. Finally, we evaluated the navigation control of 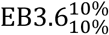 in a human arm-size, arbitrary 3D phantom vessel **(Fig 4D)**. 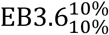 is capable of moving against a max fluid flow of 7.2 mL/min (3.9 cm/s). The experiment results shown that under a 3D condition, performance of EndoBot almost similar to the conditions observed at an angle of approximately 67.5 degrees in the first test. Across all motion tests, upstream navigation emerged as the limiting step. Notably, under all successful upstream motion conditions, 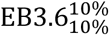 could be seamlessly repositioned downstream without losing magnetic control, underscoring its robust capabilities for round-trip navigation.

Building on the under-flow stability, magnetic navigation capabilities, and adaptive elasticity of EndoBot, we developed a wireless deployment and retrieval strategy applicable for different EndoBot sizes **(Fig S20 and Supplementary Movie 3)**. To demonstrate this approach, we used vascular access sheaths suitable for each EndoBot configuration. 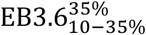 was deployed using an 2.3 mm inner diameter custom-fabricated sheath, while 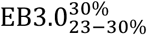 and 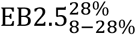 were deployed using 7F (2.1 mm inner diameter) and 6F (1.8 mm inner diameter) PINNACLE® introducer sheaths (Terumo), respectively. These sheaths were inserted into a phantom vessel with a blood flow rate of 55 mL/min for the custom-fabricated sheath and 40 mL/min for the PINNACLE® introducer sheaths.

Once the catheters were positioned within the vessel, we applied a rotating magnetic field (f_m_ = 30 Hz) with a flexible axial displacement (|d_z_|) ranging from 52 to 112 mm (corresponding to a field strength of 145 mT to 15 mT). As the EndoBots began rotating under magnetic actuation, the external magnet was moved at a speed (|**v**_m_|) between 0 and 5 mm/s to guide the EndoBot out of the sheath and into the vessel phantom. Deployment was completed under 15 seconds once the sheath reached the target vessel segment. For retrieval, we experimented and observed various scenarios with varying degrees of success. In some cases, EndoBot could move back upstream and be retrieved to the same catheter. However, we experienced reproducibility issues with this approach. The most reliable approach was using a secondary catheter at the receiver end of the vessel segment downstream. This catheter required the use of at least the same size catheter, or ∼1F larger than the initial deployment sheath. This demonstrates the adaptability of the EndoBot deployment and retrieval strategy across varying size and compression configurations, providing flexibility for diverse vascular scenarios.

Collectively, these experiments demonstrate the versatility and robust performance of EndoBot under a wide range of physiological and anatomical conditions. This capability promises effective operation in human and animal blood vessels of comparable dimensions and hemodynamic parameters **(Table S1)**.

### Biocompatibility of EndoBot

As a blood-contacting device, EndoBot and its corkscrew locomotion must result in minimal adverse interactions with blood and endothelial cells (ECs) lining the vessel intima. Although the thrombogenicity of such devices is difficult to predict, we conducted a comprehensive series of tests to evaluate coagulative properties of EndoBots, hemolytic potential, platelet activation, and cytotoxicity. These tests followed commonly accepted in vitro protocols aligned with ISO-10993-Part 4 (Biological evaluation of medical devices: Selection of tests for interactions with blood) guidelines and customized procedures tailored to our application context(*41, 42*).

The envisioned operational timescale of EndoBot within a blood vessel is short-term (<15 min). During this period, its surface material composition and locomotion could potentially trigger blood coagulation. To assess this risk, we continuously monitored the real-time translational velocity of 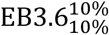 moving in a phantom vessel filled with whole bovine blood under constant magnetic actuation (∼12-15 mT, 10 Hz) for 30 min (**Supplementary Text 8.1, Fig 5A**). The blood was pre-heparinized at 0.5 U/mL, approximately matching the prophylactic dose recommended for adults undergoing endovascular procedures for deep vein thrombosis(*43*). A decrease in the robot velocity over ∼40 forward-backward traversals (∼560 cm total distance) would suggest coagulation-induced drag. However, mean velocities remained stable (5.9 ± 0.9 mm/s forward and 5.9 ± 0.8 mm/s backward), demonstrating no signs of coagulation-induced drag. This key observation provides translatable evidence for future animal and human procedures.

**Figure 5.**
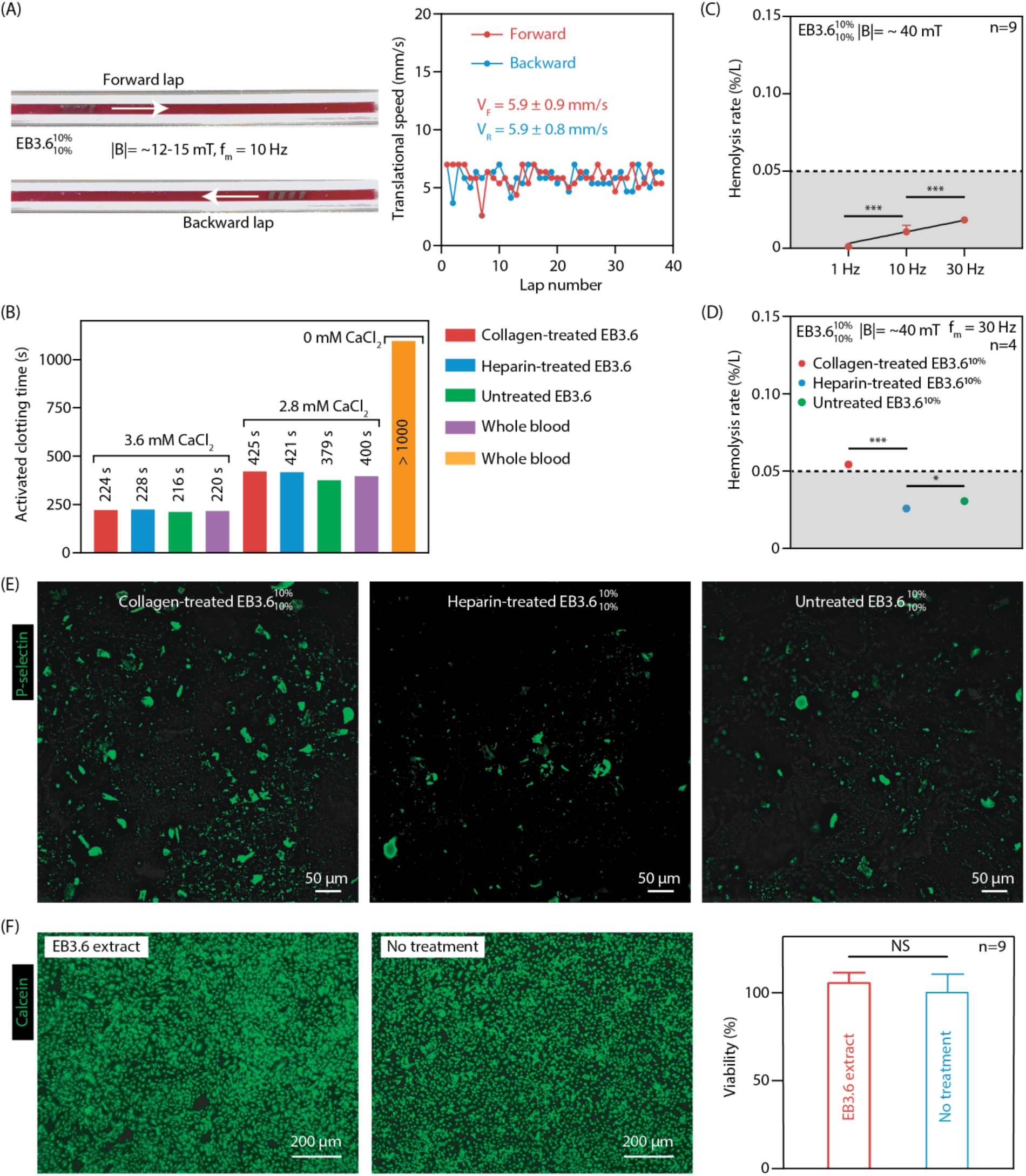
Biocompatibility evaluation of EndoBot. (A)-(E) Hemocompatibility tests. (A) 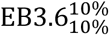 was continuously propelled forward and backward in a phantom vessel filled with fresh cow whole blood for approximately 40 round trips over 30 minutes, covering a total distance of ∼560 cm. The consistent mean velocity throughout the experiment indicates that blood coagulation did not occur (which would have increased viscosity and drag). (B) Activated clotting time (ACT) measurements in single-donor human blood following exposure to various surface-treated EndoBots. (C) Hemolysis as a function of magnetic field frequency. Assessment of hemolysis in single-donor human blood induced by the locomotion of 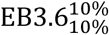 across different magnetic field rotation frequencies. (D) Hemolysis with surface treatment: Evaluation of hemolysis resulting from the locomotion of 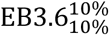 after applying surface treatments. Collagen I served as a positive control, and heparin as a negative control, in single-donor human blood. (E) Platelet activation and aggregation: P-selectin positive platelet binding and aggregation in response to EndoBot exposure. (F) Cytotoxicity assessment: Evaluation of leachable materials from EndoBot on human umbilical vein endothelial cells (HUVECs) in accordance with ISO-10993 standards. ***p < 0.0001, *p < 0.05, NS: No Significance.

To further assess the potential coagulation risks posed by the material composition of EndoBot, we measured the activated clotting time (ACT) of pre-heparinized single-donor human blood after overnight contact with the device. Heparin- and collagen-treated EndoBots served as negative and positive controls, respectively. Under these conditions, ACT values for the untreated EndoBot were comparable to both controls, indicating no heightened or reduced procoagulant effect **(Supplementary Text 8.2, Fig 5B)**. This finding remained consistent even when calcium ion concentrations were varied. Additionally, whole blood hemogram analyses revealed that all measured hematological parameters, including hemoglobin, red blood cells (RBCs), hematocrit, platelet count, white blood cell count, and differential leukocyte counts, remained within normal ranges **(Supplementary Text 8.2, Fig S21)**, showing no adverse changes in blood cell morphology or counts. The absence of detectable pro-coagulant behavior also aligns with the U.S. National Heart, Lung, and Blood Institute’s designation of silica filler-free PDMS as a reference material for blood compatibility(*44*).

Hemolysis, the rupture of RBCs and subsequent release of their intracellular components, can be induced by mechanical stress or surface interactions. Such released molecules may, in turn, trigger thrombogenesis by activating platelets(*45*). To evaluate potential hemolytic effects, we quantified free hemoglobin released from RBCs after exposing fresh, heparinized bovine blood to 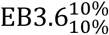 in a straight vessel phantom at varying locomotion frequencies (1, 10, and 30 Hz) for 15 min **(Supplementary Text 8.3, Fig S22)**. Higher magnetic actuation frequencies correlated with increased hemolysis, which we attribute to intensified mechanical shear forces at the blood-robot interface **(Fig 5C)**. Surface treatments also influenced hemolytic outcomes where heparin-treated EndoBots produced slightly lower levels of hemolysis compared to untreated devices, while collagen-treated EndoBots exhibited the highest hemolysis **(Fig 5D)**. Nonetheless, according to ASTM 756-17 (Standard Practice for the Assessment of Hemolytic Properties of Materials), materials are considered non-hemolytic if their hemolysis rate remains below 2% following the blood contact(*46*). Under these criteria, EndoBot and its locomotion parameters demonstrated acceptable hemolytic behavior with lower than 0.1% hemolysis under all conditions investigated.

Subsequent to the hemolysis assessment, we examined the blood-contacting inner surfaces of recovered EndoBots for platelet activation, adhesion, and aggregation, which are key indicators of thrombogenicity(*42*). We used platelet P-selectin expression as a reliable marker to characterize platelet activation and aggregation(*47*) **(Supplementary Text 8.4, Fig 5E)**. Consistent with the hemolysis rate patterns, collagen-treated EndoBots exhibited significantly elevated levels of P-selectin-expressing platelets attached to the robot surface, forming denser and larger aggregates. This outcome aligns with the known properties of collagen I, which presents epitopes for platelet-binding and activation(*48*). In contrast, heparin-treated EndoBots displayed substantially lower levels of P-selectin-positive platelet adhesion and aggregation. Although untreated EndoBots induced slightly more platelet activation and aggregation than the heparin-treated variant, these levels remained closer to the heparin-treated group, suggesting potential thrombogenic potential over long-term exposure.

To evaluate the potential cytotoxicity of EndoBot, we tested the impact of any leachable substances from its body adversely affect human umbilical vein endothelial cells (HUVECs). EndoBots were initially incubated in EC culture medium for one week at 37°C, after which the conditioned medium was applied to HUVECs without any dilution. Compared to no-treatment cultures, HUVECs exposed to EndoBot-conditioned medium showed no significant changes in cell viability or morphology (**Supplementary Text 8.5, Fig 5F**).

Collectively, these key observations demonstrate under the tested short-term operational timescale and clinically relevant pre-heparinized conditions, EndoBot operates without inducing problematic coagulation, excessive hemolysis, significant platelet activation, or cytotoxicity. Its stable robotic performance under both immobile and mobile conditions under flow, coupled with benign interactions with RBCs, platelets, and endothelial cells, provides a promising foundation for safe, short-term endovascular applications. Future studies examining longer exposure times, varying heparinization levels and dosage patterns, and more physiologically complex models will further validate its biocompatibility.

### Fluoroscopic-guided steering, localization and tracking approaches in vitro

Developing safe and effective endovascular interventions with EndoBot requires necessitates methods for real-time imaging, localization, and tracking. Since the vasculature is inherently non-transparent for visible light, visualizing small-scale robots depends on imaging techniques routinely employed in clinical practice. Interventional fluoroscopy stands out for its ability to provide deep tissue penetration, high spatial resolution, and near real-time image acquisition. These capabilities make it a powerful tool for guiding small endovascular instruments, such as catheters.

To leverage these advantages for EndoBot, we constructed a clinical C-arm-guided magnetic manipulation platform (eX-MMPt) tailored for endovascular imaging, localization, and tracking **(Supplementary Text 9, Fig 6A)**. The open workspace provided by the C-arm system accommodates the robot arm with flexible kinematics, enabling precise modulation of the magnitude and direction of the external magnetic field for actuation. As a result, EndoBot can be accurately steered through complex three-dimensional vascular pathways under fluoroscopic guidance.

**Figure 6.**
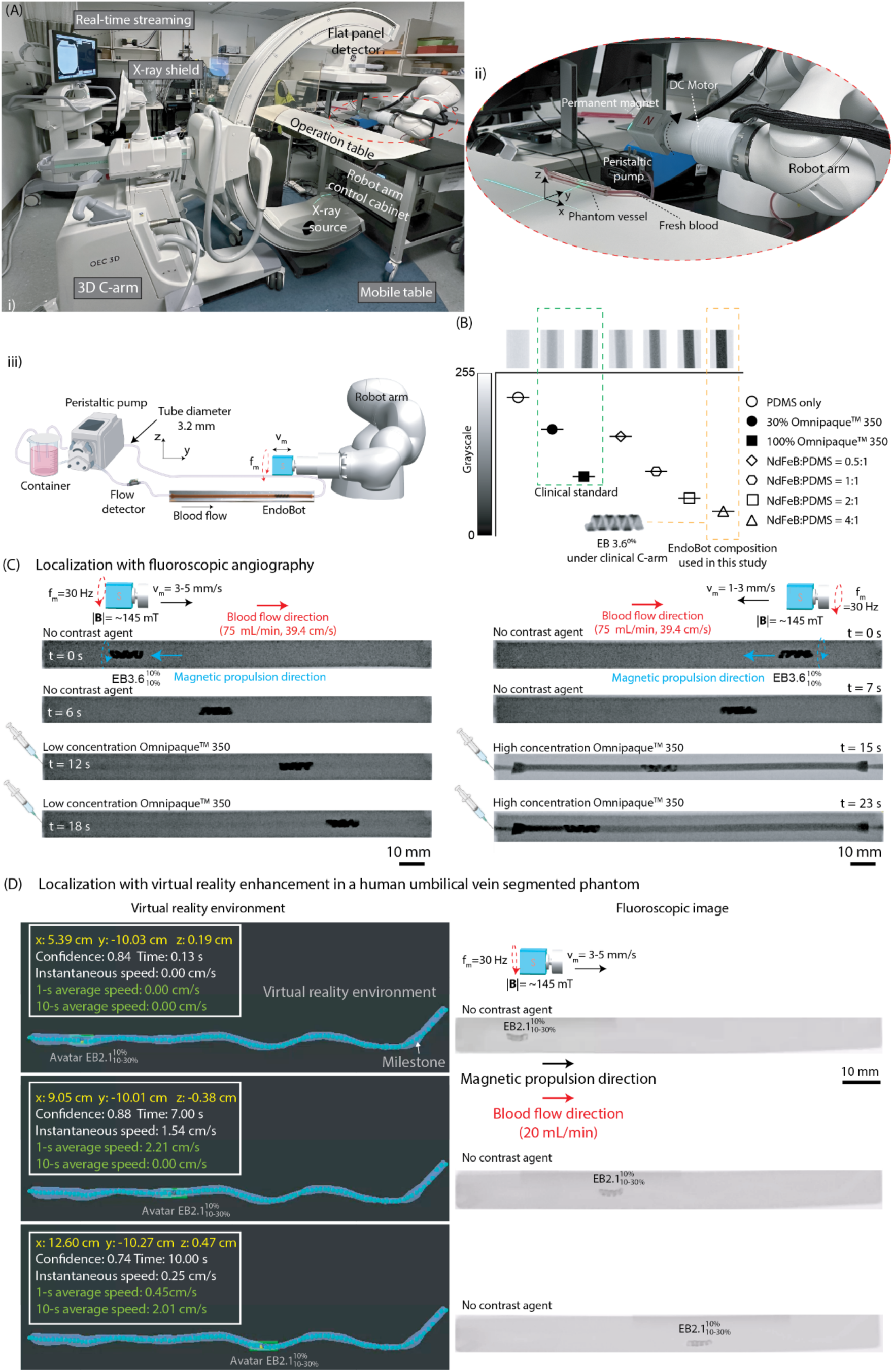
Clinically viable approaches for fluoroscopic-guided EndoBot navigation. (A) (i), (ii) X-ray fluoroscopy–guided magnetic manipulation platform (eX-MMPt) designed for human-scale micro- and millirobotic interventions. (iii) Schematic of the blood flow configuration in phantom vessels along with the magnetic actuation conditions used during robot testing. (B) Greyscale analysis of the relative fluoroscopic contrast for EndoBot precursor composites with varying NdFeB:PDMS ratios (0.5:1 to 4:1 by weight) in 1 mL syringes, benchmarked against syringes filled with Omnipaque™ 350. A concentration of 30 vol.% Omnipaque™ 350 establishes the lower threshold for clinical visibility. Notably, the EB3.6 composite (NdFeB:PDMS = 4:1) exhibits an X-ray contrast that surpasses the clinical standard, ensuring reliable detection during real-time C-arm–guided interventions. (C) Fluoroscopic angiography-based localization of 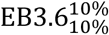 in a straight-line phantom vessel. (D) Virtual reality–enhanced localization of 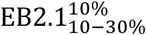 within a segmented human umbilical vein derived phantom in the absence of contrast agent injection.

To ensure the compatibility of EndoBot with clinical C-arm systems, we initially quantified its radiographic contrast relative to iohexol, an iodine-based medical contrast agent, and its endovascular formulation Omnipaque™ 350, as the clinical benchmark. Achieving sufficient contrast is critical for detecting the robot as well as for managing patient radiation exposure and its potential short- and long-term side effects(*49*). For the contrast assessment, we filled 1 mL standard syringes with varying compositions of EndoBot precursor material, i.e., different NdFeB:PDMS ratios, and compared their X-ray contrast to that of iohexol **(Fig 6B)**. For fluoroscopy-guided interventions, a medical device should exhibit contrast levels at least equivalent to 30% iohexol to ensure adequate visibility. Grayscale analysis of the samples revealed that all tested EndoBot precursor compositions exceed the 30% iohexol threshold, indicating that EndoBot would be readily visible in a clinical setting. This superior X-ray visibility arises from the higher atomic number of _60_Nd compared to _53_I, which increases X-ray attenuation and thus enhances detectability under C-arm. Notably, the refined EndoBot composition (NdFeB:PDMS = 4:1) surpassed even the contrast level of 100% iohexol, outperforming this clinical standard.

To demonstrate the magnetic propulsion of EndoBot under fluoroscopic guidance, we applied magnetic actuation within a phantom vessel at high blood flow rate of up to 75 mL/min **(Fig 6C, Supplementary Movie 4)**. We continuously visualized EndoBot and its motion at 15 fps using cinefluoroscopic (cine) imaging mode and 8 fps using digital subtraction (DS) mode **(Figs 6C, S23, and Supplementary Movie 4)**. In DS mode, EndoBot was not initially visible while remaining stationary due to the real-time frame subtraction process performed by the C-arm system. Once it started moving, it became clearly distinguishable against the blank background, leaving a white digital trace at its initial position. Although it is slower than the cine mode, DS imaging has unique advantageous for endovascular applications in the chest and head regions, where overlying bone structures can obscure vascular details.

Despite successful visualization of EndoBot and near real-time image acquisition during motion, both cine and DS modes of fluoroscopy presented significant limitations. First, they provided limited structural information within the robot workspace **(Fig 6C)**. Without clear lumen visibility, localization, tracking and navigation of the robot using external magnetic interactions could be severely compromised, endangering the procedural safety. To address this, we performed fluoroscopic angiography, similar to established clinical procedures for guiding catheters and guidewires(*50*). By injecting iohexol contrast agent into the bloodstream, we achieved lumen visualization that improved proportionally with the agent’s final concentration in the blood **(Supplementary Movie 4)**.

While fluoroscopic angiography is effective for localizing and tracking EndoBots and many other untethered milli- and microrobots in vitro and ex vivo, adapting this method for in vivo use poses additional challenges. Unlike tethered devices, such as catheters, untethered devices like EndoBot require continuous localization to prevent uncontrolled drift under magnetic propulsion. Periodic contrast agent injections can aid localization; however, continuous administration is infeasible, due to the nephrotoxicity of contrast agents and potential hypersensitivity reactions in patients(*51*).

To overcome these challenges, we recently developed an innovative virtual reality (VR)-based approach that integrates a digital twin of the robot’s operational environment, a robot avatar and real time robot position data(*52*) (**Supplementary Text 9, Figs S24, S25**). This framework synchronizes the physical and virtual workspaces through precise scale and orientation calibration, facilitated by positional data exchange via the Robot Operating System (ROS) network. The robot avatar, a virtual representation of the physical robot, mirrors its real-time position and movements within the digital twin, enabling accurate and dynamic tracking in three-dimensional space.

Fluoroscopic imaging inherently provides 2D projection data, lacking z-axis information essential for 3D tracking. To overcome this major caveat, we implemented an algorithm that infers z-axis positions by correlating 2D positional data with the pre-defined geometry of a virtual model **(Figs S26-28)**. This approach involves the introduction of virtual “milestones”, which are spherical markers with known 3D coordinates strategically placed manually along the central axis of the vessel **(Figs 6D, S27)**. These milestones span the entire navigable region within the vessel, providing a comprehensive spatial framework for localization. Each milestone is sized at approximately one-quarter of the length of EndoBot (∼3 mm), enabling precise segmentation of the robot’s trajectory. The segmentation framework allows the algorithm to infer z-axis positions directly from the 2D pixel data by mapping the robot’s projected location to the closest milestone in the virtual environment. By combining this inferred z-axis data with the x-y positional information from fluoroscopic imaging, we constructed accurate 3D positional datasets **(Figs 6D, S27)**.

To validate the robustness of this approach, we created a digital twin of a human umbilical vein segmented from a fluoroscopic 3D scan **(Supplementary Text 9, Figs 6D, S27)**. The fluoroscopic stream did not show any detail of the vascular trajectory as a model 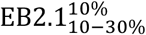 moves along the vessel. However, spatially calibrating the virtual and real environments allowed us to localize and track the motion of 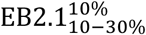 along the phantom vessel with an average delay of approximately 66 ms in the digital twin **(Fig 6D, Supplementary Text 9, Fig S29)**.

Real-time processing of visual data further allowed us to precisely determine and display average robot velocity. Even when the visibility of structural details diminished due to the passage of the robot arm or magnet through the imaging field, the EndoBot detection algorithm maintained reliable performance without interruptions or loss of localization precision. Similarly, our previous results demonstrated reliable tracking even when the visual signal is nearly indistinguishable from background noise(*52*). This highlights the significance of robot detection algorithms for ensuring the safe and effective navigation of untethered endovascular robots.

While this VR-enhanced method significantly reduces reliance on contrast agents for localizing and tracking EndoBot, future efforts should focus on extending the capability to 3D navigation in animal models. Although biplane fluoroscopy, which employs two X-ray sources could address the challenge of 3D localization limitation, it also significantly increases overall X-ray radiation exposure(*53*). Additionally, for magnetic robot control, biplane fluoroscopy imposes operational constraints on the external magnet and robotic arm, as X-ray beams are obstructed in at least one plane at any given time. Expanding the virtual twin concept offers a promising alternative, enabling precise 3D localization with a single X-ray source while minimizing radiation exposure and preserving the flexibility of robotic actuation.

### Atraumatic fluoroscopic navigation in perfused human umbilical veins ex vivo

To evaluate the mechanically adaptive, robust locomotion of EndoBot and its impact on the vessel wall integrity and ECs, we conducted navigation experiments using normothermically perfused human umbilical vein models ex vivo. Fresh human umbilical cords were obtained immediately after cesarean sections, and re-perfusion was established by circulating heparinized whole cow blood (∼37 °C) through the umbilical vein, following previously described protocols(*54*) **(Supplementary Text 10, Figs 7A, S30)**.

**Figure 7.**
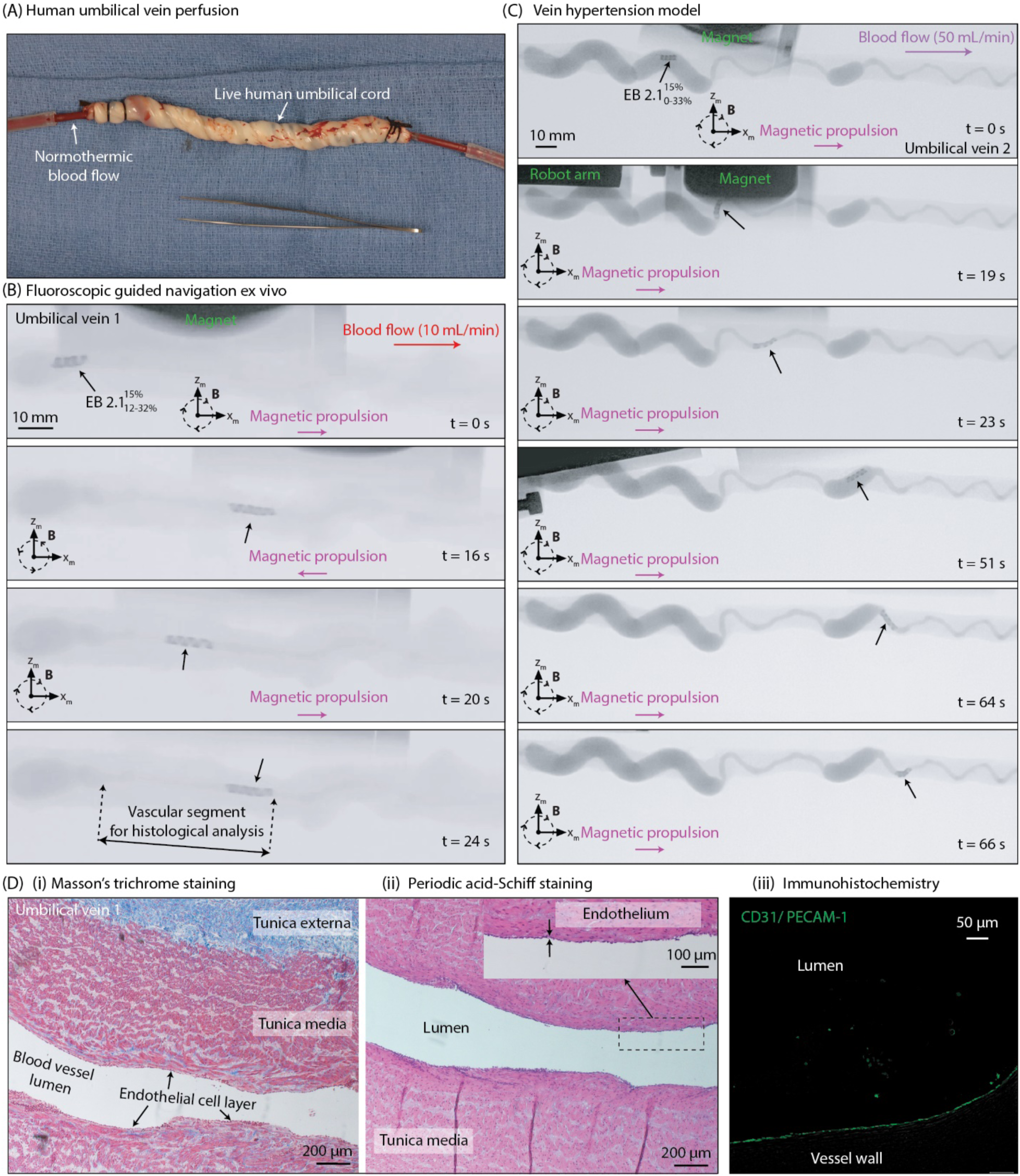
Fluoroscopic-guided EndoBot navigation in normothermically perfused ex-vivo human umbilical veins. (A) Re-perfusion of a human umbilical vein with heparinized whole blood within one hour post-cesarean section. (B) Fluoroscopic-guided navigation of 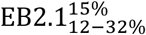 in the Umbilical Vein 1 using an iohexol contrast enhancer injected into the bloodstream. (C) Navigation of 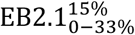 in Umbilical Vein 2, serving as a venous hypertension model. (D) After four forward and backward passes along a 60 mm segment of Umbilical Vein 1 (as marked in B), the segment was harvested for histological analysis. Longitudinal sections stained with (i) Mason’s trichrome, (ii) Periodic acid-Schiff, and (iii) CD31/ PECAM-1 reveal an intact endothelial cell layer and preserved inner vessel wall, demonstrating that robust EndoBot navigation took place without vascular injury under physiological flow conditions.

Similar to the phantom vessels, perfused veins were completely invisible under the C-arm without contrast agents **(Fig S30)**. Thus, iohexol was infused into the bloodstream to enable uninterrupted visualization of the vessel throughout the experiment. Based on the initial angiographic characterization of the lumen’s minimum and maximum diameters across the target segment, we selected EB2.1 to ensure safe navigation by adhering to two fundamental design rules for EndoBot: maintaining continuous surface contact (Rule 1) and avoiding excessive deformation beyond allowable limits **(Fig 2A, B)**.

**Figure 7B** demonstrates the forward and backward motion of EndoBot against and with a blood flow rate of 10 mL/min (**Supplementary Movie 5**). Along the target vessel segment, EndoBot 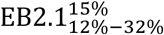 underwent dynamic elastic deformations ranging from ∼12% to 32% **(Fig S31** for moving in Umbilical Vein 1), conforming to the irregular and likely vasoconstricted lumen caused by surgical trauma. Frame-by-frame analysis of deformation patterns confirmed that EndoBot’s ability to elastically adapt to these irregularities enhanced the robustness and safety of navigation while maintaining blood flow patency **(Supplementary Movie 5, Figs 7B, S31)**. This mechanically compliant design prevented excessive force on the vessel walls, reducing the risk of localized damage to the endothelial lining.

In a worst-case scenario experiment, conducted in Umbilical Vein 2, EndoBot was subjected to supraphysiological blood flow conditions (50 mL/min, >200 cm/s flow velocity at regular segments) and extreme vessel dilation **(Fig 7C)**. Even under this condition, we maintained magnetic stability of EndoBot in the dilated segment without drifting, successfully moving it into the regular segments without losing helical structure and causing disruptions to the blood flow.

After four round trips of EndoBot within a ∼60 mm segment of the vein (<5 min) under optimized actuation parameters (|**B**| = 100 mT, f_m_ = 30 Hz), no evidence of EC denudation was observed. Histological analysis (**Supplementary Text 8)** of the vascular segments navigated by EndoBot confirmed that the overall tissue morphology of Umbilical Veins 1 and 3 remained intact (**Fig 7D**, **Fig S32**). Hematoxylin and eosin (H&E), Masson’s trichrome, and periodic acid-Schiff (PAS) staining demonstrated the preservation of vessel wall structure and the single-cell endothelial lining. Additionally, CD31/PECAM-1 immunohistochemistry staining validated that the outer lining cells were indeed endothelial cells, further confirming atraumatic navigation.

Each umbilical vein presented a distinct three-dimensional vascular structure, yet EndoBot demonstrated consistent and reproducible locomotion and safety outcomes across all tested models. For example, in Umbilical Vein 2, 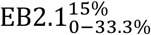 experienced deformation ranging from 0% to 33% (**Fig 7C, S33**), while in Umbilical Vein 3, 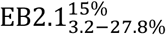 experienced deformation ranging from 3.2% to 27.8% (**Fig S34**). These results highlight the ability of EndoBot to safely adapt to varying vessel geometries, supporting its potential for use in anatomically diverse or pathological vasculature.

The ex vivo results emphasize the translational potential of EndoBot for safe and effective endovascular interventions. The mechanically adaptive locomotion ensures robust navigation in highly variable vessel geometries, such as those encountered in diseased vasculature with stenoses, dilations, or vasospasms. Furthermore, the preservation of vessel wall and EC integrity suggests that EndoBot could perform interventional tasks with reduced vascular trauma compared to traditional devices, such as catheters or stents, which often cause endothelial denudation or mechanical injury.

### Endoluminal drug delivery

The drug delivery mechanism of EndoBot relies on its ability to perform effective surface crawling along the vessel lumen, ensuring constant contact with the vessel surface. This design prevents blood flow from prematurely washing away the coating before it is fully deposited to the vessel wall. As EndoBot emerges from a vascular sheath or catheter and navigates through arteries or veins, its outer hydrophobic coating layer is gently transferred to the lumen via mechanical rubbing against the vessel surface **(Fig 8A)**. For a successful delivery, this process must form a stable, flow-resistant drug depot layer without causing downstream fragmentation.

**Figure 8.**
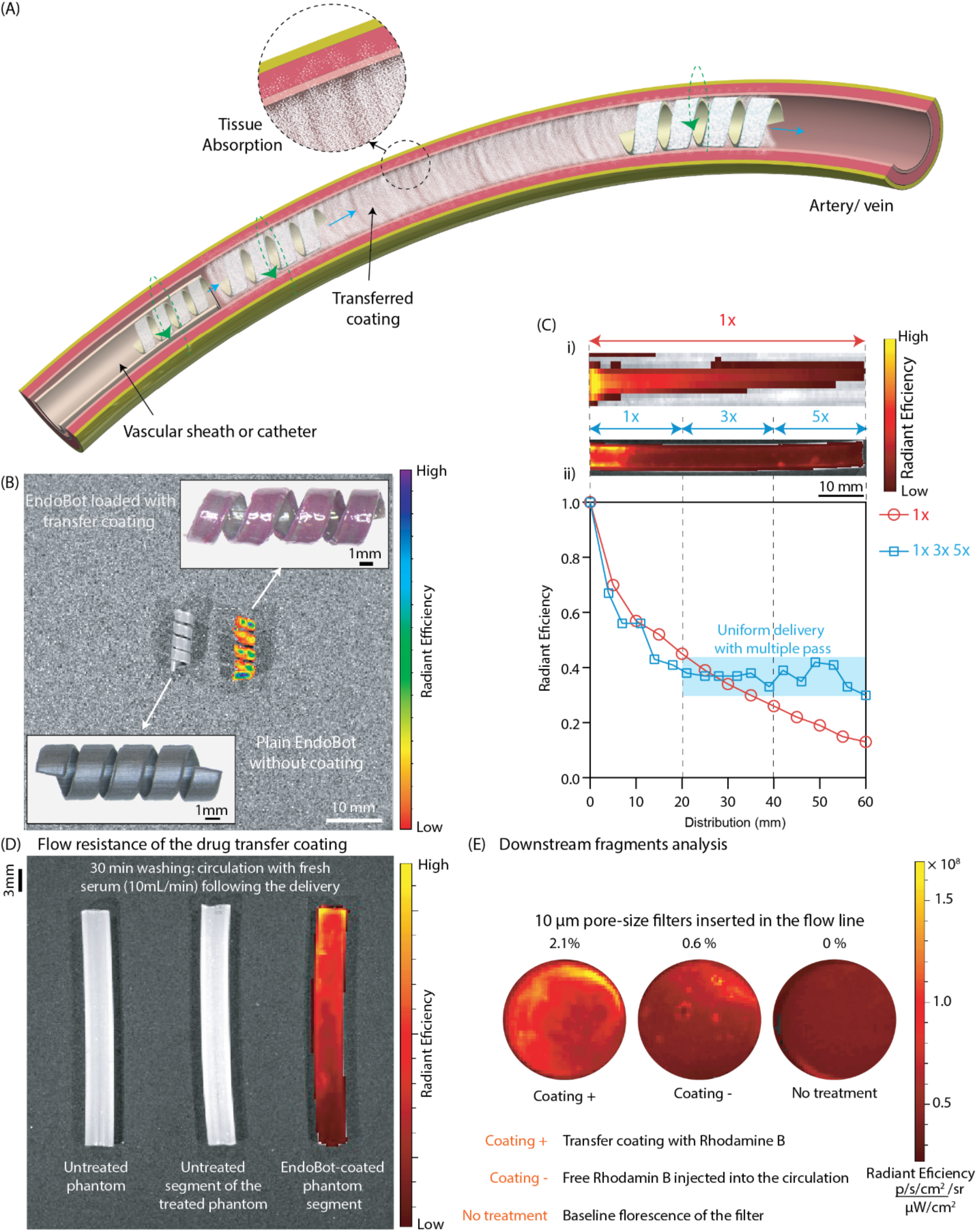
Endoluminal drug delivery using EndoBot. (A) Conceptual illustration: As EndoBot exits a vascular sheath or catheter and navigates through arteries or veins, its outer hydrophobic coating is gradually transferred to the vessel lumen via gentle mechanical rubbing against the vessel wall. (B) In vivo imaging system (IVIS) images of plain and loaded EndoBot with a refined transfer coating that incorporates Rhodamine B as a fluorescent reporter. This optimized formulation readily wets the surface of EndoBot, forming a stable, lubricious, and self-supporting outer layer. (C) (i) Endoluminal drug delivery demonstration (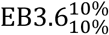, |***B***| = 50 mT, f_m_ = 10 Hz) along a 60-mm vascular phantom segment. (ii) Under continuous magnetic actuation, the spatial distribution of Rhodamine B is modulated by adjusting the number of passes, achieving precise spatial control and enhanced distribution uniformity along the vessel. (D) Coating stability against flow-induced erosion: The Rhodamine B-delivered vessel segment was recirculated with fresh serum at 10 mL/min for 30 minutes. The coating remained firmly attached to the vessel walls during the washing step. (E) Coating fragmentation analysis: After 30 minutes of closed-circuit circulation, only 2.1% of the circulating coating material was retained by the filter. Considering that free Rhodamine B exhibits approximately 0.6% non-specific adsorption to the filter, about 98.5% of the coating material released into circulation consisted of fragments smaller than 10 µm, confirming the safety of this drug delivery procedure with minimal risk of downstream perfusion obstruction.

To enable this capability, we formulated a transfer coating using acetyl tributyl citrate (ATBC), an FDA-approved pharmaceutical excipient(*55*). ATBC is a slightly hydrophobic, oil-like plasticizer that can protect pharmaceutical agents from rapid dissolution into the bloodstream(*55*). A similar coating strategy is employed in current DCB systems. For example, TransPax™ platform (Boston Scientific) utilizes a citrate ester excipient combined with low-dose paclitaxel (2 µg/mm^2^) to adhere the therapeutic agent to the vessel surface during balloon inflation(*56*).

ATBC exhibits a blood half-life of approximately 0.5 and 5 hours in rats and humans, respectively due to serum esterase activity(*55*). It is metabolically cleared easily by rat and human liver microsomes rapidly with half-life less than 30 min. These properties make ATBC an excellent material foundation for developing a biodegradable, blood-compatible transfer coating tailored for localized drug delivery and sustained release on the endothelium and vessel wall.

To enhance the mechanical properties of the transfer coating for improved surface applicability and increased resistance to aqueous dissolution and flow-induced erosion, we additionally incorporated poly(methyl methacrylate) (PMMA), another hydrophobic, biocompatible polymer, into the formulation. We assessed the coating stability using Rhodamine B, a fluorescent dye molecule, as a reporter. Increasing the PMMA ratio in the coating formulation resulted in a more viscous material **(Supplementary Text 11.1, Figs S35A,B, and Supplementary Movies 6** for optimal formulation containing 3.7 wt% PMMA in ATBC). We observed the same trend in its resistance to serum dissolution with increased PMMA ratio **(Supplementary Text 11.2, Fig S35C)**. An optimal formulation containing 3.7 wt% PMMA in ATBC was identified, balancing surface applicability, dissolution resistance, and flow-induced erosion stability **(Fig S35D)**. Further increasing the PMMA ratio produced more rigid coating that could not be effectively used for transfer via the mechanical rubbing strategy with EndoBot.

The refined coating formulation readily wets the surface of EndoBot, forming a stable, lubricant and self-supporting outer layer **(Fig 8B)**. To evaluate the impact of the coating layer on the mechanical properties of the EndoBot backbone, EB3.6^0%^ with and without coating were moved in a conical-shaped narrowing vessel phantom filled with whole blood under the standard magnetic actuation conditions **(Fig S36)**. Both configurations exhibited a maximum magnetic propulsion-mediated deformation of around 42%, demonstrating that the thin coating does not compromise the robot’s safety limits. This allows EndoBots of various sizes with transfer coating layers to be deployed in standard vascular sheaths, enabling their use in both in vitro and in vivo vessels.

To demonstrate the endoluminal drug delivery concept, we employed 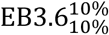 for a ∼60 mm long phantom vessel segment filled with bovine serum **(Supplementary Text 11.2, Fig 8C)**. The in vivo imaging system (IVIS) revealed that when EB3.6^10%^ traverses the vessel segment a single pass (1x), the spatial distribution of the payload demonstrates a gradual decline in radiant efficiency along the length of the segment. This decline may result from coating depletion, where the device releases its payload as it moves along the segment, leaving less material available for deposition further along the vessel. Additionally, the single pass may provide less physical interaction or contact time with distal regions, leading to lower efficiency in coating those areas. However, increasing the number of passes (1x, 3x, 5x) within the segment enables a more uniform transfer of the coating, with higher retention of radiant efficiency at distal regions. This pattern indicates that the amount of delivery can be spatially modulated by adjusting the number of passes, achieving both spatial control and enhanced distribution uniformity along the vessel.

To assess the stability of the coating against flow-induced erosion, the vessel segment was recirculated with fresh serum at 10 mL/min for 30 minutes **(Fig 8D)**. The washing step failed to remove the coating from the vessel walls, indicating that the coating is stable and flow-resistant. Surface stability of the coating was also validated against extreme vortex forces in a serum-filled environment **(Fig 8D)**. These features demonstrate that once EndoBot is delivered to the lumen, it forms a stable, dissolution- and shear-resistant drug reservoir, creating a window for the drug to be transferred across the vessel wall locally.

Fragmentation of the coating during and after the delivery could impact downstream perfusion and lead to ischemic tissue injury. To evaluate coating fragmentation behavior, we placed 10 µm pore-size filters in the circulation line **(Supplementary Text 11.3, Fig 8E)**. After 30 minutes of closed-circuit circulation, only 2.1% of the circulating coating material was retained in the filter. Considering that free Rhodamine B exhibits ∼0.6% non-specific adsorption to the filter, approximately 98.5% of the coating material passing into circulation was smaller than 10 µm, confirming the safety of this delivery procedure with minimal risk of downstream perfusion obstruction.

### Fluoroscopic navigation and drug delivery in live rat to inferior vena cava

To evaluate the safety and performance of EndoBot for navigation and drug delivery in a live animal model, we selected the inferior vena cava (IVC) of Sprague Dawley (SD) rats as the ultimate testbed. The mean peak flow velocity in the ∼300g SD rat IVC ranges from 6–10 cm/s(*57*). Previous ex vivo results with EB2.1 (Fig 7) and in vitro tests with EB3.6 (Fig 3) demonstrated successful navigation at flow velocities up to 100 cm/s in comparable sized vessels. Based on these findings, we hypothesized that EndoBot would safely perform in the normal rat IVC.

For this study, we used 300–375 g male SD rats (n=4) in a non-survival experiment (**Supplementary Text 10**). We employed the EB2.1 variant, designed to operate under compression within the range of 10–35%, applying radial pressures of < 0.25 kPa—consistent with safety thresholds derived from preliminary studies in human umbilical veins (Fig 2). Deployment of EB2.1 was performed under general anesthesia using standard 6F PINNACLE® introducer sheaths (Terumo) vascular sheaths and a micropuncture wire, under fluoroscopic guidance and magnetic control **(Supplementary movies 6,7)**.

To minimize thrombosis in the catheter/sheath used for the EndoBot, the sheath/catheter was prefilled with heparin (1000 IU/mL). During and after successful deployment of the catheter/ sheath into the IVC (the access position for the catheter/ sheath in the IVC is shown in **Fig 9A(i)**, and a close-up view post-catheterization showing the insertion site on the rat’s body is shown in **Fig 9A(ii))**, additional heparin was applied to the abdominal area to reduce thrombosis caused by blood leakage from the IVC into the abdomen. After the catheter/sheath was successfully deployed into the IVC, the rat was transferred to the working area under the C-arm and robot arm for the locomotion step, as shown in **Fig 9A(iii)**.

**Figure 9.**
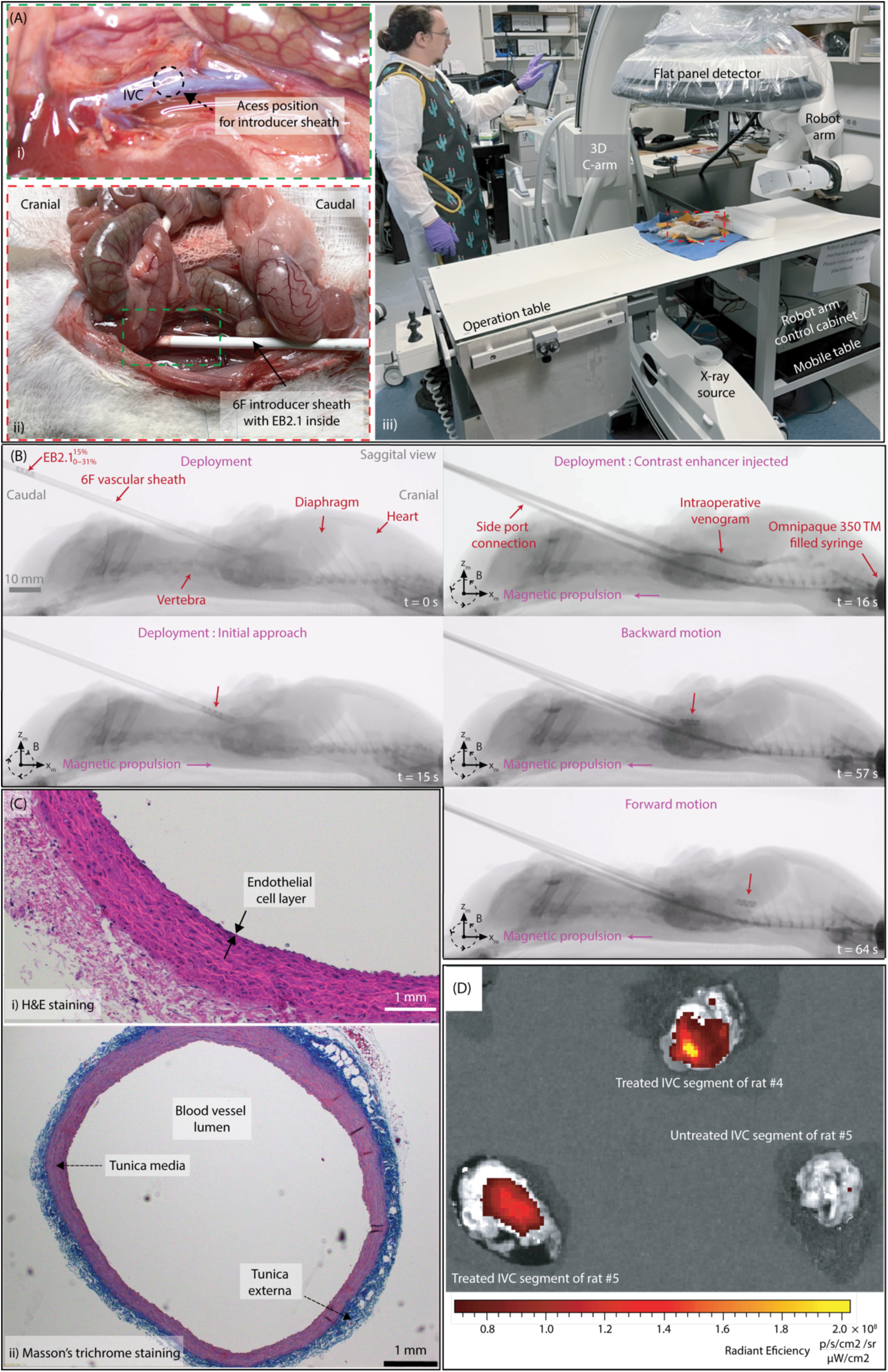
Fluoroscopic-guided navigation and drug delivery in the inferior vena cava (IVC) of live rats. (A) Experimental setup during the EndoBot intervention: (i) Positioning of the vascular introducer sheath for IVC access. (ii) Close-up view of the sheath insertion site on the anesthetized rat. (iii) Workspace configuration showing the use of the human-scale eX-MMPt together with the anesthetized rat. (B) EndoBot navigation: Sequential snapshots depict the navigation of EndoBot (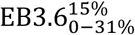) advancing and retreating along the IVC of rat #3. (C) (i) H&E, (ii) Masson’s trichrome stained transverse sections of the IVC from rat #3 after seven complete forward and backward navigation cycles, demonstrating an intact endothelial cell layer and the preserved vessel wall, respectively. (D) IVIS examination of the post-mortem extracted IVC, demonstrating the successful delivery of Rhodamine B to a 30-mm segment of the vessel in rats #4 and #5. Photo in (B) belongs to Husnu H. Alabay, a co-author of this manuscript.

For all experiments, the initial approach for deploying the EndoBot into the IVC involved using a magnetic field (|**B**| = ∼100 mT and f_m_ = ∼ 10-30 Hz) with magnetic propulsion aligned in the target movement direction. Once the EndoBot reached the end of the catheter/ sheath, which was in contact with the IVC, a second approach was employed using a contrast enhancer injection. Flushing the contrast enhancer not only facilitated the successful deployment of the EndoBot into the IVC but also cleared the IVC structure, preparing it for the subsequent locomotion step, as shown in **Fig 9B** and **Supplementary Movies 7 and 8**. EndoBot was maneuvered along a 65 mm segment of the IVC up to the diaphragm for 3–7 round trips, completed within less than 5 minutes. Total local radiation exposure during fluoroscopy was maintained below 20 Gycm^2^, ensuring minimal risk from imaging. For each rat, we performed a fluoroscopic angiography during the deployment of the vessel to understand the IVC structure. However, this is short lived (< 2 s) and there is an injection limit for a live rat, we moved EB2.1 without seeing the vessel line. As a result, for the first rat, we failed to move back the robot from the diaphragm, and it continued moving past the diaphragm into heart irreversibly (**Supplementary movie 9)**. Despite this initial failure, subsequent trials demonstrated EndoBot’s repeatability and reliability in traversing the IVC. The results confirmed that EB2.1 maintained consistent contact with the lumen wall, experiencing compression levels between 0% and ∼31%, in line with its design criteria.

Similar to the ex vivo experiment, several segments of the IVC where EB2.1 motion occurred were isolated for histological analysis. Histological analysis using H&E, Masson’s trichrome showed no evidence of endothelial cell (EC) denudation (**Fig 9C** for H&E, Masson’s trichrome, and PAS staining results). Examination of the vascular segments navigated by EndoBot confirmed that the overall tissue morphology remained intact, as was similarly observed in Umbilical Veins 1 and 3 (**Figs 7D** and **S32**). These results underscore the biocompatibility and safety of EndoBot during in vivo endovascular navigation.

To prove the transferability of our coating material, after successful EndoBot locomotion, segments of the IVC from rat #4 and #5, where EndoBot motion was observed, were isolated for IVIS imaging. Additionally, one segment without EndoBot motion was also isolated from rat #4 as a control sample. The IVIS imaging results are presented in **Fig 9F**. The results demonstrated that EndoBot successfully transferred the coating material onto the IVC surface without causing damage or affecting the endothelial cells.

## DISCUSSION

EndoBot is designed to meet several critical performance criteria for safe and effective endoluminal drug delivery. These include: (i) robust locomotion under physiological and supraphysiological flow conditions, allowing adaptive crawling along irregular tubular surfaces (**Figs 4,6,7,9**); (ii) mechanical adaptability to accommodate non-uniform vascular geometries, including vasoconstriction and periodic vessel motion (**Figs 7,9**); (iii) stability within the vessel lumen, resisting drift even in the absence of external magnetic fields (Fig 3); (iv) X-ray visibility for real-time fluoroscopic navigation (**Figs 6,7,9**); (v) atraumatic navigation without causing damage to endothelial cells (ECs) or vessel walls (**Figs 7, 9**); (vi) biocompatibility, ensuring compatibility with blood while avoiding hemolysis, coagulation, or cytotoxicity (Fig 5); (vii) wireless deployment and retrieval through standard vascular access sheaths (Fig 9) and; (viii) controlled endoluminal drug transfer with tunable deposition parameters (**Figs 8,9**).

This study demonstrates the in vitro, ex vivo and in vivo feasibility of each of these criteria, laying the groundwork for the translational development of EndoBot. Notably, this is the first study to demonstrate the safety of untethered millirobots in a live animal model using a human-scale, fluoroscopy-guided magnetic manipulation platform. EndoBot addresses a critical gap in the development of localized, atraumatic drug delivery systems that utilize untethered millirobots. By enabling early, preventive interventions that protect vascular integrity, EndoBot paves the way for a paradigm shift in vascular disease management while representing a significant advance in mobile, untethered microrobotic technologies and establishing a robust foundation for transformative innovations in interventional medicine.

