## Supplementary Information File for "Fluoroscopic-Guided Magnetic Soft Millirobot for Atraumatic Endovascular Drug Delivery"

for

**Table S1. Blood Vessels Suitable for Safe Deployment of EndoBot in Endoluminal Delivery Applications**

| | Vessel type | Description | Diameter (mm) | Blood flow rate (mL/min) | Blood flow velocity (cm/s) |
|---|---|---|---|---|---|
| Human | Middle cerebral Artery | Lateral part of the brain | 2 - 5 (*1*) | 93-149 (*2*) | 27-39 (*2*) |
| | Brachial Artery | Major artery of the upper arm. | 3 - 4 (*3, 4*) | 5-90 (*5*) | 4.2-11.13 (*6*) |
| | Coronary Artery | Superficial vein in the arm. | 2 - 4 (*7, 8*) | ~70-80 (*9*) | 13-41 (*10*) |
| | Umbilical Vein | Vein that connects the placenta to the fetus | 1.9 - 2.4 (*11*) | 63-450 (*12*) | 8-10 (*12*) |
| Rat | Inferior vena cava | Large vein from the lower body to the heart | 3.1 - 3.3 (*13*) | 10 (*14*) | 7.6-10 (*14*) |

(*9*) for 100 g myocardial tissue at rest.

(*10*) During angiography, velocities in systole and diastole

(*6*) Average of right and left radial arteries

(*14*) Peak velocity

**Supplementary Text 1: Fabrication of EndoBot**

**Fig S1** navigates the process of EndoBot fabrication. It started with the design of a master positive mold and a pin with precise dimensions using SolidWorks (Dassault Systèmes, Vélizy-Villacoublay, France), followed by 3D printing with the FormLabs 3B+ 3D printer (FormLabs, Somerville, MA). Subsequently, the master positive mold and pin were used create a negative mold made from silicon elastomer. A 10:1 weight ratio of SYLGARD™ 184 Silicone Elastomer Base and Curing Agent (base to crosslinker mass ratio) were prepared and cured in a heater at 80°C for 2 hours. Prior to the subsequent step, the surface of the negative mold was passivated using plasma treatment and alcohol passivation, as described in (*15*).

Various base:crosslinker mass ratios and NdFeB:PDMS powder mass ratios were used in the refinement process **(Supplementary text 2)**. All precursor components were mixed thoroughly followed by degassing under vacuum. Subsequently, the composite was cast into the passivated negative PDMS mold and cured inside the heater at 80°C for 2 hours. Following the curing, the EndoBot was gently demolded.

Finally, the EndoBot was magnetized in a direction perpendicular to its major helical axis using our custom-made magnetic yoke under 1.1 T **(Fig S2)**. **Fig S3** shows the repeatability of the EndoBot fabrication and scalability across a diameter range of 1.0-4.0 mm.

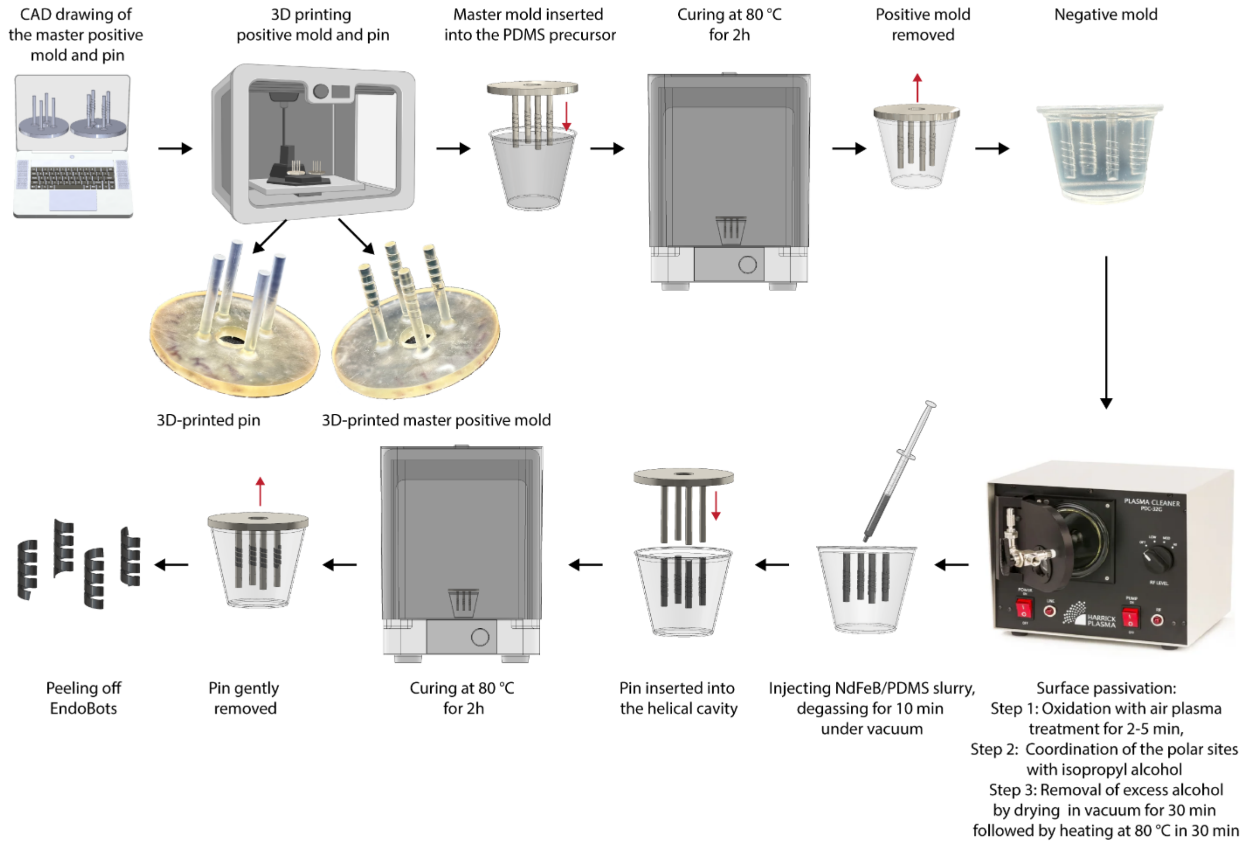

**Figure S1. Fabrication methods and production steps for EndoBot.**

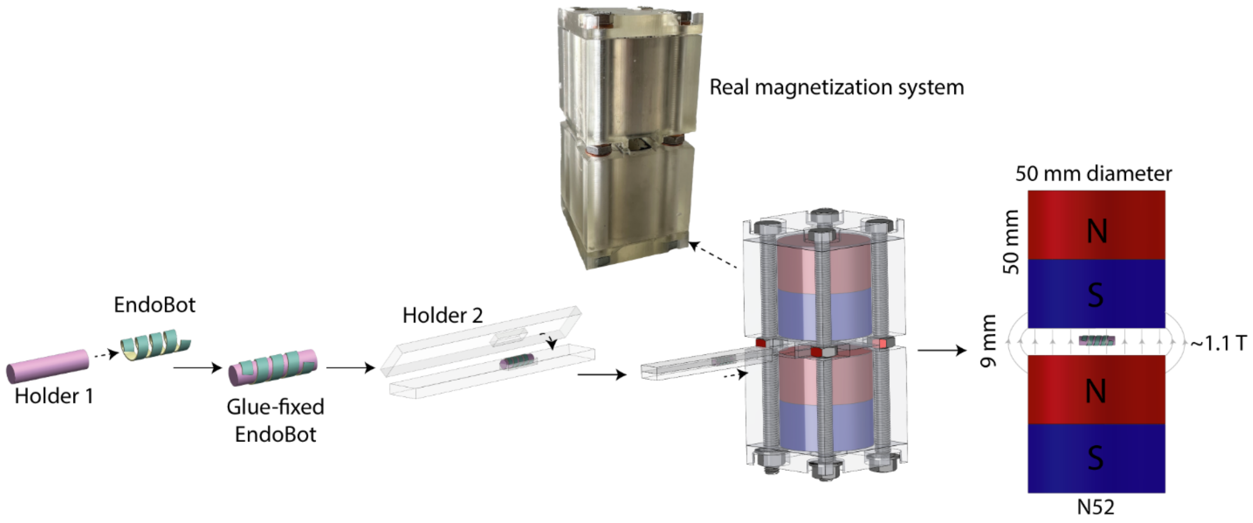

**Figure S2. Protocol for magnetization of EndoBot.** To generate a strong and directionally well-controlled $\mathbf{M_R}$, uniform magnetic field exceeding > 1 T, a pair of cylindrical N52 NdFeB magnets were positioned with their opposite poles facing each other, separated by a small gap of 9 mm. The EndoBot was secured to a rod (holder 1) using a water-soluble adhesive. This rod was then placed inside a closed container (holder 2) and exposed to the magnetic field, ensuring that $\mathbf{M_R}$ is created perpendicular to the main helical axis of EndoBot.

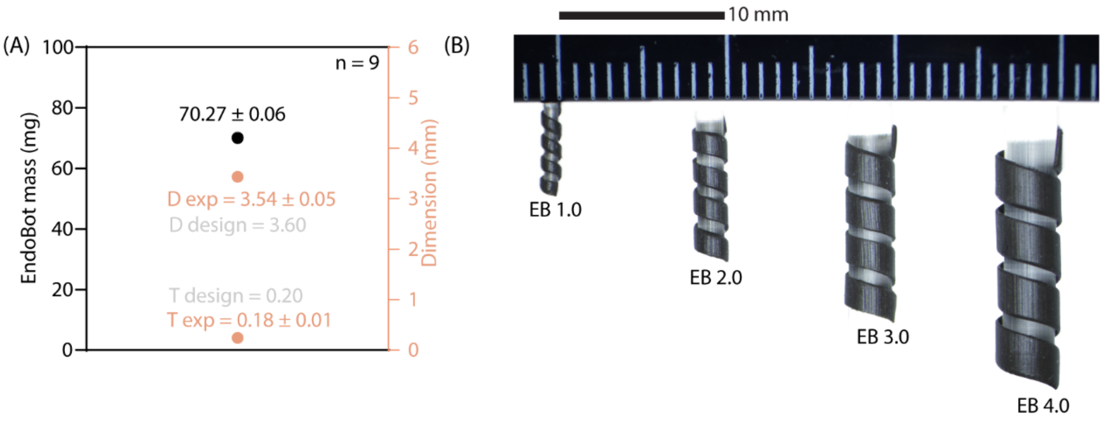

**Figure S3. Consistency and scalability of the EndoBot fabrication method.** (A) Consistency of the fabrication process. (B) Scalability in fabricating EndoBot with diameters ranging from 1.0 to 4.0 mm.

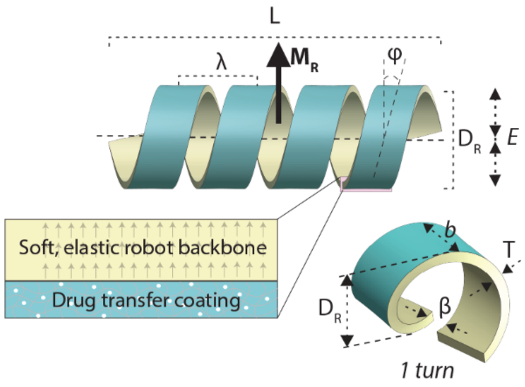

**Table S2.** Key structural and compositional parameters of the mechanically adaptive corkscrew surface-crawling mechanism of EndoBot.

| | Parameters | As-fabricated | Range of change during motion | Impact | Rationale basis |
|---|---|---|---|---|---|
| Structural parameters | *$L/D_R$ | 3.3 | 3.3–7.0 | Magnetic torque distribution | Adopted from (*16-18*) |
| | $L/\lambda$ | 4.8 | 4.8–6.7 | Redistribution of the cross-section stress | Adopted from (*16, 19, 20*) |
| | $b/\beta$ | 2.6 | 2.6–3 | Maximizing contact interaction surface of the robot | Rationally developed (this study) |
| | φ | 14° | 14° | High blood flow patency<br>High lumen contact | Rationally developed (this study) |
| | *T* | 0.2 mm | 0.2 mm | Impose mechanical cost to unintended deformation | Empirically refined (Fig.S4 and S7) |
| Compositional parameters | $\|\mathbf{M}_R\|$ | 1.8 × $10^{-3}$ (A.m$^2$) | 1.8 × $10^{-3}$ (A.m$^2$) | Maximizing magnetic torque | Empirically refined (Fig. S6) |
| | *E* | 1.99 kPa | 1.99 kPa | | Empirically refined (Fig. S4) |

* *L* = number of turns × helical pitch length + 2 × half of helical blade thickness

= number of turns × $\lambda$ + 2 × $b$/2

**Supplementary Text 2: Design and refinement of EndoBot**

**Table S2** outlines the key structural and compositional features of a scalable EndoBot prototype.

The mechanical behavior of EndoBot is primarily governed by its effective elastic modulus (*E*) across the cross-section of the helical axis. This parameter plays a pivotal role in determining: (1) the radial force exerted by EndoBot on the vessel lumen, (2) its ability and extend to conform dynamically to the vascular cross-section, and (3) its capacity for seamless surface crawling through corkscrew locomotion with magnetic actuation.

To achieve an optimal balance between adaptive deformation and vascular safety while presenting maximum magnetic actuation capacity, we systematically refined three critical structural and compositional parameters that collectively govern EndoBot's mechanical compliance, magnetic responsiveness, and structural integrity.

**(A) Refinement strategies:**

**Refinement 1:** *T* was systematically varied at 0.1, 0.2, and 0.4 mm, while maintaining a fixed NdFeB:PDMS mass ratio of 4:1 and a PDMS base:crosslinker mass ratio of 10:1.

**Refinement 2:** The NdFeB:PDMS mass ratio was varied at 1:1, 2:1, and 4:1 while maintaining a fixed PDMS:crosslinker mass ratio of 10:1 and a fixed *T* of 0.2 mm.

**Refinement 3:** The base:crosslinker mass ratio was varied at 2:1, 5:1, and 10:1 while maintaining a constant NdFeB:PDMS mass ratio of 4:1 and a fixed *T* of 0.2 mm.

**(B) Outcome Measures of Refinement Strategies:**

**1. Radial force ($\mathbf{F}_{\text{radial}}$) (Fig S4):**

$$\mathbf{F}_{\text{radial}} = \mathbf{F}_{\text{intrinsic}} + \mathbf{F}_{\text{extrinsic}} \quad (1)$$

$\mathbf{F}_{\text{intrinsic}}$: Reaction force from stored elastic energy under compressive strain ε

$|\mathbf{F}_{\text{intrinsic}}| \propto E$, thus, $|\mathbf{F}_{\text{intrinsic}}| \propto T^{2.54}$, $|\mathbf{F}_{\text{intrinsic}}| \propto (\text{NdFeB:PDMS mass ratio})^{1.08}$, and $|\mathbf{F}_{\text{intrinsic}}| \propto \varepsilon$.

$\mathbf{F}_{\text{extrinsic}}$: Radial force caused by magnetic pulling force in $z_R$-direction calculated as

$$\boldsymbol{F}_{\text{mag}}(\boldsymbol{r}_{\text{mR}}) = (\boldsymbol{M}_{\text{R}} \cdot \nabla)\mathbf{B}(\boldsymbol{r}_{\mathbf{mR}}) \quad (2)$$

where $|\boldsymbol{M}_{\boldsymbol{R}}| \propto \text{T}$ and $|\boldsymbol{M}_{\boldsymbol{R}}| \propto (\text{NdFeB:PDMS})^{0.78}$.

At maximum magnetic field condition (|**B**| = ~145 mT or $|d_z|$ = 52 mm) and maximum ε of elastic region (35%), when *T* = 0.4 mm and NdFeB:PDMS = 4:1, $|\mathbf{F}_{\text{radial}}|$ reaches its maximum value (~421 mN). In this configuration, the radial pressure exerted on the lumen surface is **~5.4 kPa**, below the endothelial damage threshold.

**2. Recovery rate after compression (Fig S5).** Recovery was tested through cyclic deformation ($\varepsilon$ = 10–35%) and measured by the recovered diameter ($D_{rec}$):

$$Recovery\ Rate\ (\%) = 100\ \times\ (1 - \frac{D_R - D_{rec}}{D_R}) \tag{3}$$

At *T* = 0.1 mm, recovery was limited to 87.2 ± 2.3 %. Increasing *T* to 0.2 mm and 0.4 mm improved recovery rates to 94.0 ± 0.6 % and 97 ± 0.8%, respectively. NdFeB:PDMS and base:crosslinker ratios minimally influenced recovery.

**3. Conformal deformation capacity under magnetic propulsion (Fig S6).** Using a blood-filled conical vessel model (A-labelled, **Fig S6**), the conformal deformation capacity (*C*) and reversibility of EB3.6 were assessed under constant magnetic actuation. Parameters yielding the highest reversible deformation were identified and summarized in **Table S3-5**. We can see that $C \propto 1/T$ and $\propto$ log(NdFeB:PDMS).

**4. Vascular patency (Fig S7A,B).** The impact of *T* on vascular patency (*S*) was defined as the percentage of the lumen cross-sectional area that remains unoccupied by the EndoBot, indicating the extent to which blood can flow freely through the vessel.

$$S\ (\%) = 100\ \times\ (\frac{S_{lumen} - S_{project}}{S_{lumen}}) \tag{4}$$

where $S_{project}$ is cross-sectional area occupied by EndoBot and $S_{lumen}$ is the total cross-sectional area of the lumen (**Fig S7A)**.

The *S* results shown in **Fig S7B**. $S \propto T^x$, with x being -0.28, -0.32 and -0.49 for $\mathrm{EB3.6}_{0\%}^{0\%}$, $\mathrm{EB3.6}_{10\%}^{10\%}$ and $\mathrm{EB3.6}_{35\%}^{35\%}$, respectively. This indicates that $S \propto 1/T$ and also indicates that the S has a high sensitivity to compression.

**5. Stability under blood flow (Table S3-5).** EndoBot's stability was evaluated under increasing flow rates to define operational safety limits. The stability of the EndoBot improved with increasing *T*, as the resulting higher radial force provided enhanced anchoring against the blood flow.

<u>**Comprehensive evaluation:**</u> The interplay between *T*, NdFeB:PDMS ratio, and base:crosslinker ratio underpins the balance between elasticity, structural robustness, and magnetic actuation. These parameters directly influence deformation recovery, radial pressure, and propulsion efficiency, all of which are critical for EndoBot's safe and effective performance across diverse

vascular conditions. To determine the optimal EndoBot design, we conducted a comprehensive evaluation of key outcome measures, summarizing findings in **Tables S3–S5**.

To determine the optimal *T*, we assessed designs at T = 0.1 mm, 0.2 mm, and 0.4 mm while maintaining fixed ratios for other parameters (Refinement 1). The results clearly indicate that *T* is a critical variable for tuning the helix's deformation behavior. The effective elastic modulus (*E*) across the cross-section scales nearly cubically with *T* ($E \propto T^{2.54}$), so even moderate increases in *T* substantially raise $\mathbf{F}_{\mathbf{intrinsic}}$ arising from stored elastic energy under compressive strain ($\mathbf{F}_{\mathbf{intrinsic}} \propto T^{2.54}$). At the same time, increasing *T* reduces both the *C* during magnetic propulsion ($T \propto 1/C$) and *S* ($T \propto 1/S$). However, increasing *T* also raises $\mathbf{F}_{\mathbf{radial}}$ and $\mathbf{M}_{\mathrm{R}}$ ($|\boldsymbol{M_R}| \propto T$). Consequently, *T* was judiciously optimized to balance robust structural integrity and mechanical compliance for adaptive crawling, while minimizing radial pressure on the vessel wall and preserving vessel patency. With *T* = 0.1 mm, EndoBot has a low radial force demonstrated an enhanced *C* (up to 47%), which expanded adaptability to varying vascular diameters. However, this came at the cost of reduced structural integrity, leading to uncontrolled intra-structural deformations under magnetic interactions (**Fig S7**, *T* = 0.1 mm in Refinement 1) and a low recovery rate after compression (~87.2%). The low radial force also diminished lower stability (90 mL/min at *T* = 0.1 mm). Additionally, the lower $|\boldsymbol{M_R}|$ required a stronger external magnetic field for effective propulsion. With *T* = 0.4, EndoBot has a high radial force exhibited enhanced deformation recovery rate, exceptional stability under blood flow rate (~98% recovery, >150 mL/min at T = 0.4 mm in Refinement 1) and high $|\boldsymbol{M_R}|$. However, this came at a significant cost, including a substantial reduction in *C* (only 28% at *T* = 0.4 mm in Refinement 1), limiting adaptability to varying vascular diameters. Additionally, achieving high radial force by increasing *T* led to a significant reduction in *S*, which may severely restrict the blood vessel. Furthermore, the high radial force at *T* = 0.4 mm resulted in elevated radial pressure; although below the endothelial damage threshold (5.4 kPa vs. 12 kPa), it remained relatively high, warranting further reduction to enhance long-term safety. Given these trade-offs, *T* = 0.2 mm was selected as the optimal design parameter.

While *T* was the primary tuning parameter, additional refinements were required to optimize mechanical properties and magnetic responsiveness. Similarly, to systematically determine the optimal NdFeB:PDMS ratio, we evaluated designs with NdFeB:PDMS ratio at 1:1, 2:1, and 4:1 while keeping other ratios fixed (Refinement 2). The NdFeB:PDMS ratio played a crucial role in determining key performance metrics, directly influencing $\mathbf{M}_{\mathrm{R}}$ ($|\boldsymbol{M_R}| \propto (\mathrm{NdFeB{:}PDMS})^{0.78}$), the intrinsic radial force ($|\mathbf{F}_{\mathrm{intrinsic}}| \propto (\text{NdFeB:PDMS mass ratio})^{1.08}$) and *C* ($C \propto \log(\mathrm{NdFeB{:}PDMS})$), with minimal impact on recovery rate, stability and *S*. Increasing the NdFeB:PDMS mass ratio significantly enhanced magnetic force, torque, and propulsion efficiency, leading to improved maneuverability and control. Although a higher NdFeB:PDMS mass ratio increased intrinsic radial pressure, it remained well below the endothelial damage threshold (maximum radial pressure is only 1.0 kPa). Meanwhile, increasing the NdFeB:PDMS ratio enhanced stability (reaching 110 mL/min at the NdFeB:PDMS mass ratio of 1:1 and 120 mL/min at the NdFeB:PDMS mass ratio of 4:1, both measured at $\varepsilon$ = 10%). However, this increase came at the expense of reduced C, with values of 42%, 44%, and 50% for NdFeB:PDMS mass ratios of 1:1, 2:1, and 4:1, respectively. Given the substantial benefits of higher propulsion efficiency, stability while maintaining sufficient *C*, we prioritized maximizing the NdFeB:PDMS ratio and selected an optimal value of 4:1.

Finally, to identify the optimal base-to-crosslinker ratio, we assessed designs with base-to-crosslinker ratios of 2:1, 5:1, and 10:1, while keeping other ratios fixed (Refinement 3).The base-to-crosslinker ratio played a crucial role in $|\mathbf{F}_{intrinsic}|$, and *C*, with minimal impact on stability, recovery rate, $|\mathbf{M}_R|$, and *S*. Since all EndoBot designs demonstrated radial pressures well below the endothelial damage threshold, we only focused on evaluation of *C*. Increasing the base-to-crosslinker ratio from 2:1 to 5:1 (a 2.5-fold increase) led to a slightly reducing *C* (from 36% to 33%). However, further increasing the ratio from 5:1 to 10:1 (a 2-fold increase) resulted in a significantly improving *C* (from 33% to 42%). Given the substantial benefits of enhanced *C* while maintaining sufficient stability, recovery rate, $|\mathbf{M}_R|$, and *S*, we selected the optimal ratio at 10:1.

As a result, the chosen design (T = 0.2, NdFeB:PDMS = 4:1, and base:crosslinker = 10:1) fulfills the requirements for a high stability under blood flow (> 150 mL/min), high *C* (42%), high *S* (68.2% under maximum $\varepsilon$ of 35%) and a high recovery rate (94%). This ensures compatibility with almost all types of human blood vessels, as shown in **Table S1**.

A similar strategy was applied to EB2.1, yielding comparable performance. The final optimal designs for EB3.6 and EB2.1 are summarized in **Table S6**.

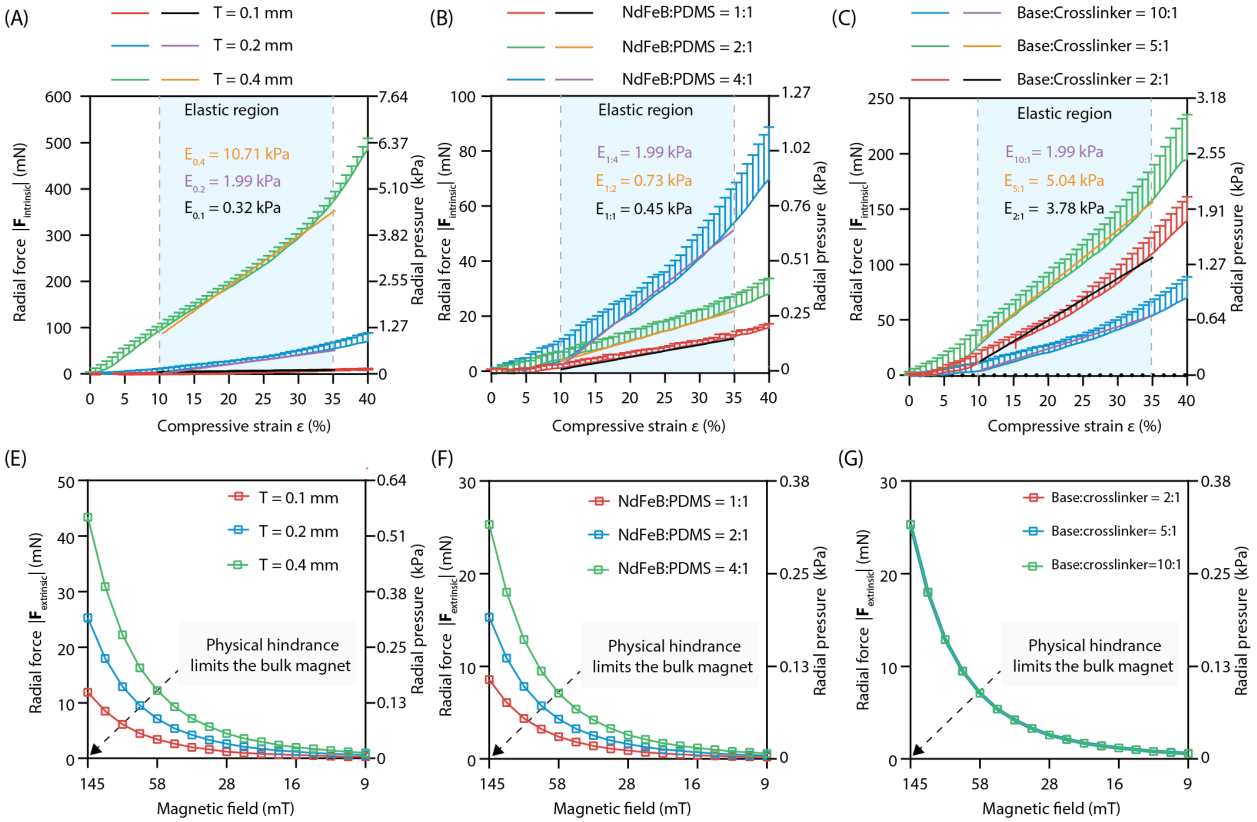

**Figure S4. Intrinsic radial force |$F_{intrinsic}$| and extrinsic radial force |$F_{extrinsic}$| dynamics of refined EB3.6.** (A) and (E) Radial forces |$F_{intrinsic}$| and |$F_{extrinsic}$| for Refinement 1, respectively. (B) and (F) Radial force |$F_{intrinsic}$| and |$F_{extrinsic}$| for Refinement 2, respectively. (C) and (G) Radial forces |$F_{intrinsic}$| and |$F_{extrinsic}$| for Refinement 3, respectively. All fitting lines in Fig S4 has $R^2$ > 0.98.

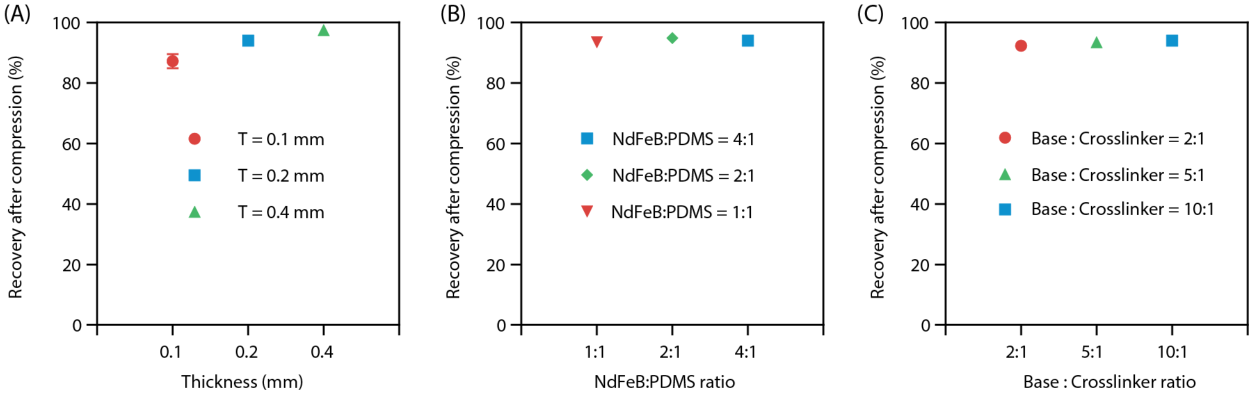

**Figure S5. Recovery rate after cyclic compressive strain ε from 0%-35%.** (A) Refinement 1. (B) Refinement 2. (C) Refinement 3.

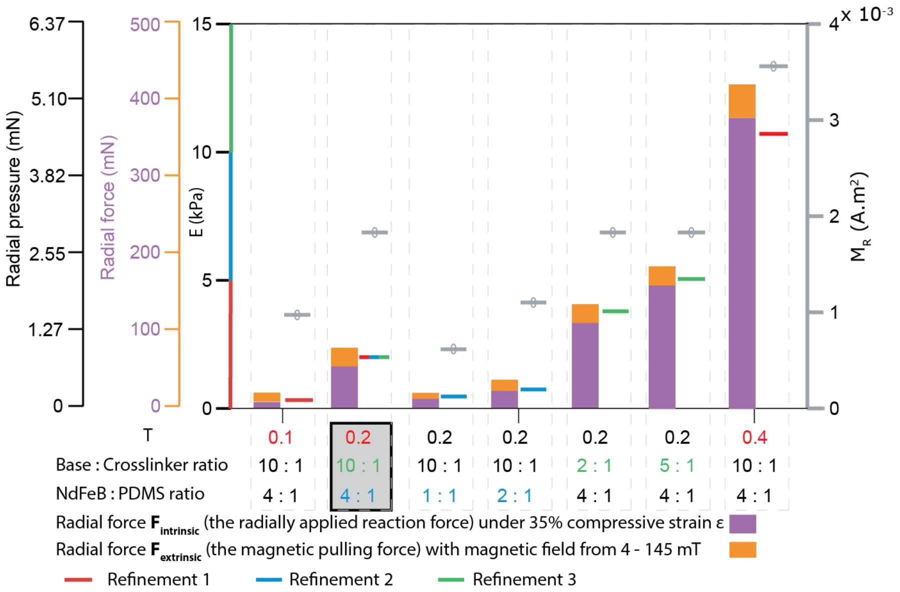

**Figure S6. Summary of the cyclic compressive deformation tests for the three refinement groups elucidating the key structural and compositional components of EndoBot.**

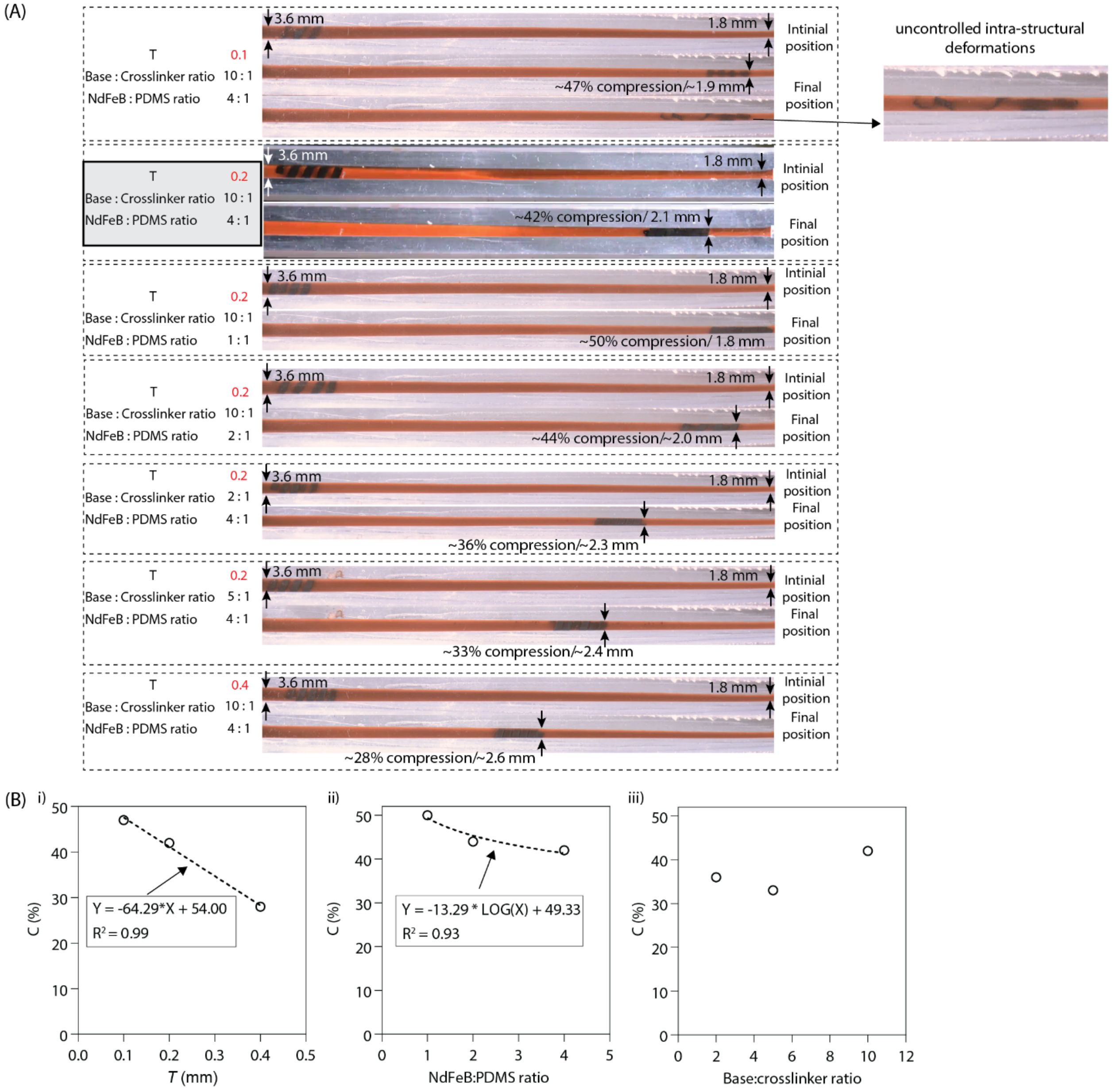

**Figure S7. Outcome measurement of the conformal deformation capacity under magnetic propulsion.** (A) EB 3.6 designs with varying *T*, base-to-crosslinker ratios, and NdFeB:PDMS mass ratios were tested under a rotating magnetic field of 4–145 mT within vessel of diameters ranging from 3.6 to 1.8 mm to assess their *C*. The initial magnetic field was set at 4 mT and gradually increased as the vessel diameter decreased. (B) Plots of *C* as a function of (i) *T*, (ii) NdFeB:PDMS mass ratios, and (iii) base-to-crosslinker mass ratios.

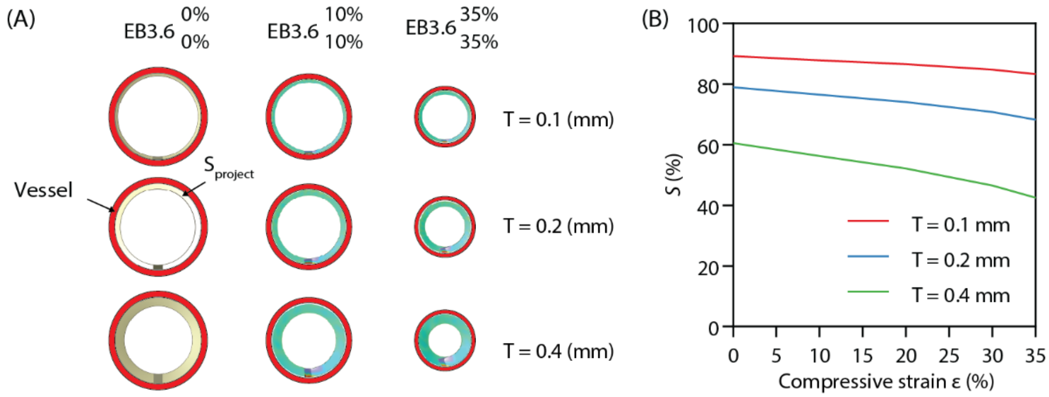

**Figure S8. The impact of *T* on vascular patency *S*.** (A) Demonstration of *S* for different EB3.6 designs with varying *T* and compressive strain ε. (B) *S* of different EndoBot designs for various *T* values and compressive strain ε.

**Table S3. EndoBot performance for Refinement 1**

| **Thickness (T):** | **0.1** | | **0.2** | | **0.4** | |
|---|---|---|---|---|---|---|
| **EndoBot:** | $\mathbf{EB3.6}_{10\%}^{10\%}$ | $\mathbf{EB3.6}_{35\%}^{35\%}$ | $\mathbf{EB3.6}_{10\%}^{10\%}$ | $\mathbf{EB3.6}_{35\%}^{35\%}$ | $\mathbf{EB3.6}_{10\%}^{10\%}$ | $\mathbf{EB3.6}_{35\%}^{35\%}$ |
| Max average radial pressure (kPa) | 0.2 | 0.3 | **0.4** | **1.0** | 1.8 | 5.4 |
| Stability under blood flow rate without the assistance of external magnetic fields (mL/min) | 90 | 100 | **120** | **>150** | >150 | >150 |
| Conformal deformation capacity *C* (%) | 47 | | **42** | | 28 | |
| Recovery rate after compression (%) | 87.2 | | **94.0** | | 97.5 | |
| Blood patency *S* (%) | 87.9 | 83.4 | **76.6** | **68.2** | 56.2 | 42.5 |

**Table S4. EndoBot performance for Refinement 2**

| **NdFeB : PDMS ratio:** | **1:1** | | **2:1** | | **4:1** | |
|---|---|---|---|---|---|---|
| **EndoBot:** | $\mathbf{EB3.6}_{10\%}^{10\%}$ | $\mathbf{EB3.6}_{35\%}^{35\%}$ | $\mathbf{EB3.6}_{10\%}^{10\%}$ | $\mathbf{EB3.6}_{35\%}^{35\%}$ | $\mathbf{EB3.6}_{10\%}^{10\%}$ | $\mathbf{EB3.6}_{35\%}^{35\%}$ |
| Max average radial pressure (kPa) | 0.1 | 0.3 | 0.2 | 0.5 | **0.4** | **1.0** |
| Stability under blood flow rate without the assistance of | 110 | 120 | 115 | >150 | **120** | **>150** |

| | | | | | | |
|---|---|---|---|---|---|---|
| external magnetic fields (mL/min) | | | | | | |
| Conformal deformation capacity *C* (%) | 50% | | 44% | | **42%** | |
| Recovery rate after compression (%) | 93.5 | | 94.9 | | **94.0** | |
| Blood patency *S* (%) | 76.6 | 68.2 | 76.6 | 68.2 | **76.6** | **68.2** |

**Table S5. EndoBot performance for Refinement 3**

| **Base:crosslinker** | **2:1** | | **5:1** | | **10:1** | |
|---|---|---|---|---|---|---|
| **EndoBot:** | $\mathbf{EB3.6}_{10\%}^{10\%}$ | $\mathbf{EB3.6}_{35\%}^{35\%}$ | $\mathbf{EB3.6}_{10\%}^{10\%}$ | $\mathbf{EB3.6}_{35\%}^{35\%}$ | $\mathbf{EB3.6}_{10\%}^{10\%}$ | $\mathbf{EB3.6}_{35\%}^{35\%}$ |
| Max average radial pressure (kPa) | 0.5 | 1.7 | 0.7 | 2.4 | **0.4** | **1.0** |
| Stability under blood flow rate without the assistance of external magnetic fields (mL/min) | >150 | >150 | >150 | >150 | **120** | **>150** |
| Conformal deformation capacity *C* (%) | 36% | | 33% | | **42%** | |
| Recovery rate after compression (%) | 92.3 | | 93.4 | | **94.0** | |
| Blood patency *S* (%) | 76.6 | 68.2 | 76.6 | 68.2 | **76.6** | **68.2** |

**Table S6.** Blueprints of EB2.1 and EB3.6 used in this study

| **Physical parameters** | **EB2.1** | **EB3.6** |
|---|---|---|
| Overall length, *L* (mm) | 8.0 | 12.0 |
| Fabricated diameter, $D_R$ (mm) | 2.1 | 3.6 |
| Thickness, *T* (mm) | 0.2 | 0.2 |
| Pitch length, $\lambda$ (mm) | 1.7 | 2.5 |
| Groove, $\beta$ (mm) | 0.5 | 0.7 |
| Helical blade thickness *b* (mm) | 1.2 | 1.8 |
| Helical angle, $\varphi$ (°) | 14 | 14 |
| Surface area ($mm^2$) | 32.4 | 78.5 |
| PDMS base to crosslinker ratio | 10:1 | 10:1 |
| NdFeB : PDMS ratio | 4:1 | 4:1 |

| | | |
|---|---|---|
| Magnetic moment × $10^{-3}$, $|\mathbf{M_R}|$ ($A.m^2$) | 0.7 | 1.8 |

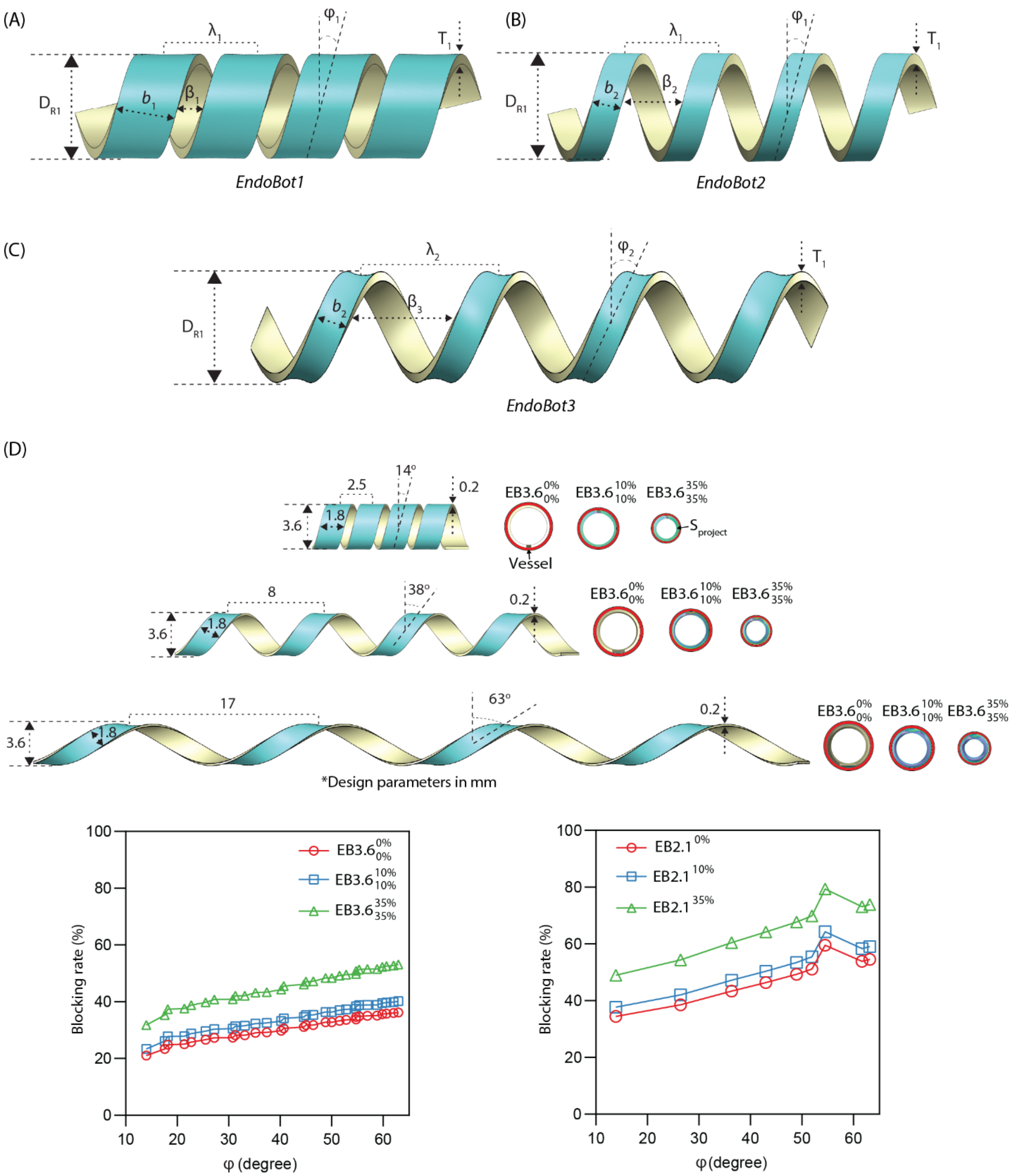

**Figure S9. Design optimization of EndoBot to enhance contact with the vessel lumen while reducing the cross-sectional area obstructed by EndoBot.** (A-C) Different EndoBot designs with the same $D_R$, T while changing *b*, $\lambda$, β and φ. To enhance contact with the vessel lumen the EndoBot design should maximize b and minimize β and φ, enabling robust crawling locomotion and efficient drug transfer. For example, when $\lambda$ is constant, reducing *b* and increasing $\beta$ leads to decrease the contact between EndoBot and the vessel lumen due to a reduction of the drug transfer coating area (Fig S9A and Fig S9B). In aother example, for the same *b*, increasing both $\lambda$ and $\beta$, as well as increasing φ, although the same drug transfer coating area is maintained but still reduces contact between EndoBot and the vessel lumen due to the EndoBot surface becoming more concave (Fig S9B and FigS9C). In the figure, $\lambda_1 < \lambda_2$, $b_1 > b_2$, $\beta_1 < \beta_2 < \beta_3$, and $\varphi_1 < \varphi_2$. (D) Different EndoBot designs with the same $D_R$, T and b while changing φ. We can see that the cross-sectional area obstructed by EndoBot decreases when φ is reduced, thereby enhancing vessel patency during deployment and navigation.

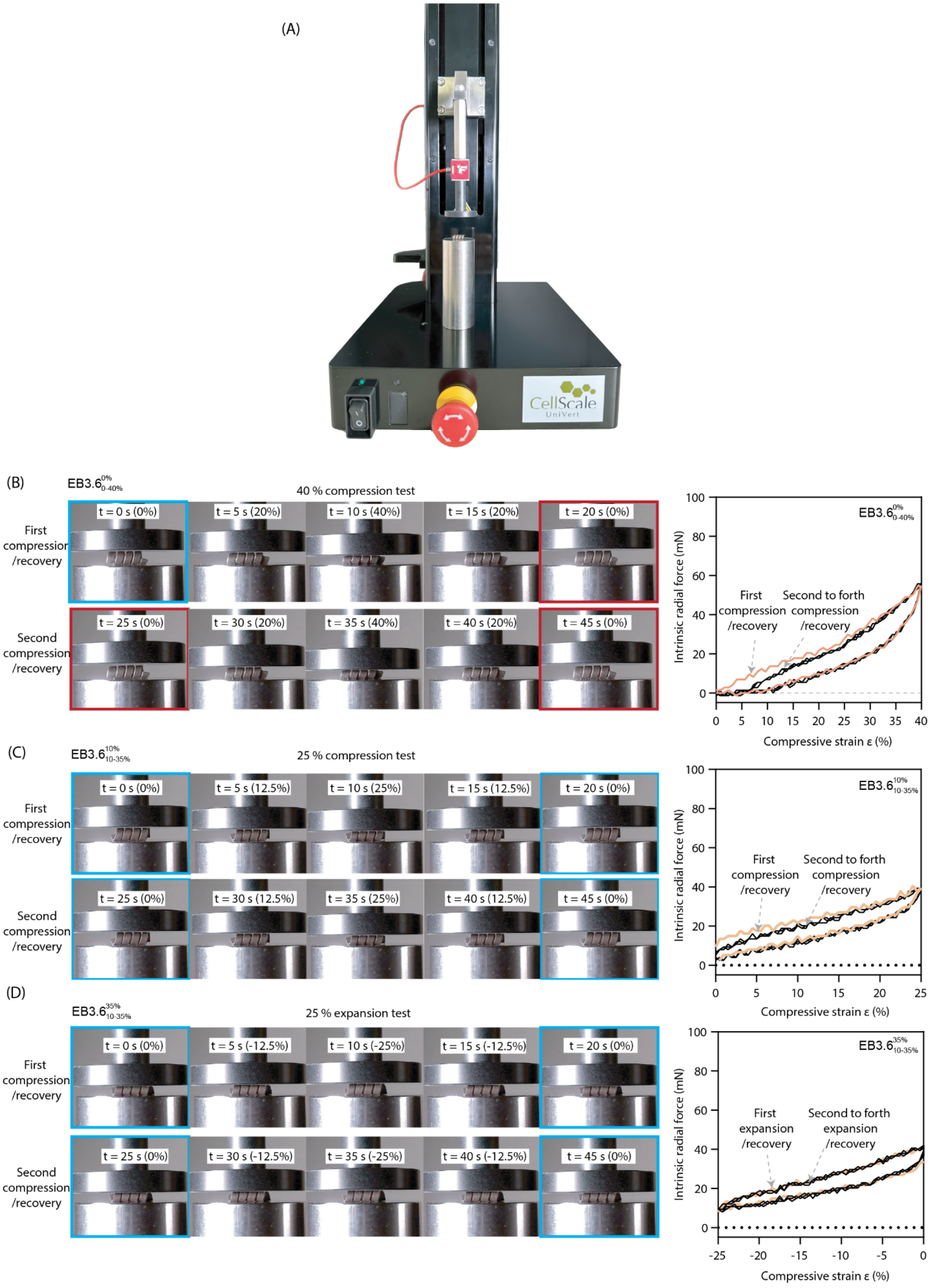

(A)
CellScale
UniVert
(B)
$EB3.6^{0\%}_{0\text{-}40\%}$
40 % compression test
First compression /recovery
Second compression /recovery
t = 0 s (0%)
t = 5 s (20%)
t = 10 s (40%)
t = 15 s (20%)
t = 20 s (0%)
t = 25 s (0%)
t = 30 s (20%)
t = 35 s (40%)
t = 40 s (20%)
t = 45 s (0%)
First compression /recovery
Second to forth compression /recovery
Intrinsic radial force (mN)
Compressive strain ε (%)
(C)
$EB3.6^{10\%}_{10\text{-}35\%}$
25 % compression test
First compression /recovery
Second compression /recovery
t = 0 s (0%)
t = 5 s (12.5%)
t = 10 s (25%)
t = 15 s (12.5%)
t = 20 s (0%)
t = 25 s (0%)
t = 30 s (12.5%)
t = 35 s (25%)
t = 40 s (12.5%)
t = 45 s (0%)
First compression /recovery
Second to forth compression /recovery
Intrinsic radial force (mN)
Compressive strain ε (%)
(D)
$EB3.6^{35\%}_{10\text{-}35\%}$
25 % expansion test
First compression /recovery
Second compression /recovery
t = 0 s (0%)
t = 5 s (-12.5%)
t = 10 s (-25%)
t = 15 s (-12.5%)
t = 20 s (0%)
t = 25 s (0%)
t = 30 s (-12.5%)
t = 35 s (-25%)
t = 40 s (-12.5%)
t = 45 s (0%)
First expansion /recovery
Second to forth expansion /recovery
Intrinsic radial force (mN)
Compressive strain ε (%)

**Figure S10. Mechanical adaptability of EndoBot under repetitive compressive stress.** (A) Mechanical test instrument (Cellscale Univert with load cell 2.5 N) used for measuring the intrinsic radial force exerted by EndoBot under compression. (B) Intrinsic radial force dynamics of $\mathrm{EB}_{0-40\%}^{0\%}$ under compression from 0% to 40% compressive strain ε. (C) Intrinsic radial force dynamics of $\mathrm{EB}_{10-35\%}^{10\%}$ under compression from 0% (10% when including the initial 10% bias compression) to 25% (35% total when including the initial 10% bias compression) compressive strain ε. (D) Intrinsic radial force dynamics of $\mathrm{EB}_{10-35\%}^{35\%}$ under expansion from 0% (35% when including the initial 35% bias compression) to -25% (10% total when including the initial 35% bias compression) compressive strain ε. Four cycles of testing were conducted for each sample (Fig 2Cii). Within each cycle, there was a preload phase of 0 s, a stretch phase of 10 s, a hold phase of 0 s, a recovery phase of 10 s, and a rest phase of 5 s.

**Supplementary Text 3:** ***Vessel phantom design and fabrication***

This paper focuses on demonstrating a drug delivery method targeting the surface of the lumen wall. To achieve this, vessels without bifurcation and with lumen diameter around 3 mm (the vessel types with lumen diameter around 3 mm is shown detail in **Table S1** were selected to validate our concept. For vitro experiment, we prepared two types of phantoms: one using PDMS-based vessels and the other using platinum-cured silicone tubing-based vessels. Detailed design parameters for each vessel phantom are provided in **Table S7**.

**Table S7. Phantom vessels used in this article**

| Figure | Label | Design | Material |
|---|---|---|---|
| Fig. 2F | A | 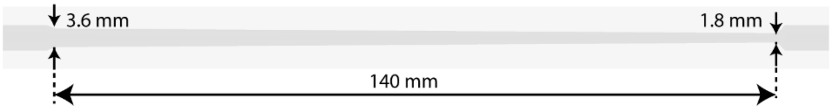 | PDMS |
| Fig. 2G, 3C | B | 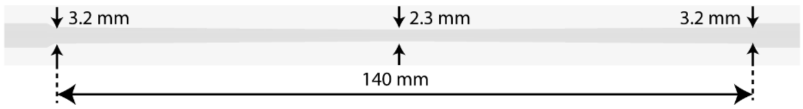 | PDMS |
| Fig. 3A Fig. 4B-C, Fig. 6C-D | C | 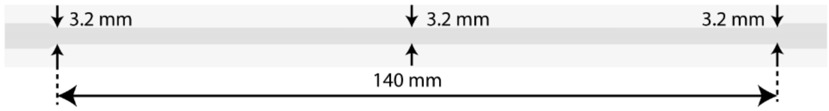 | PDMS |
| Fig. 3B | D | 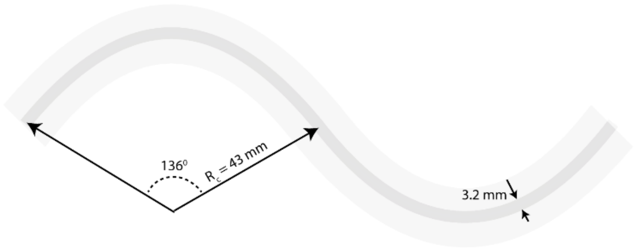 | PDMS |
| Fig.4B -C | E | 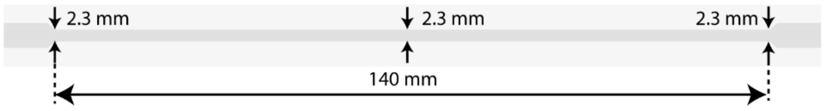 | PDMS |
| Fig. 3D | F | | platinum-cured silicone tubing |

| | | |
|---|---|---|
| | | 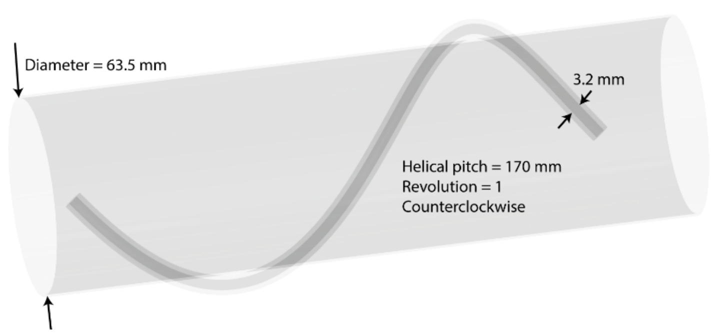 |

3.1 *PDMS-based vessel phantom*

Due to its mechanical stability, similar mechanical properties (with a Young's modulus of 2.6 MPa for PDMS (*21*) compared to approximately 1-3 MPa for human vessels (such as arteries(*22, 23*)), and its ability to easily change diameter as desired, along with its widespread acceptance in biomedical applications(*24*), PDMS elastomer was primarily used for constructing our vessel phantoms.

The fabrication process for PDMS-based vessels is illustrated in **Fig S11**. Initially, a CAD mold was designed using SolidWorks. The mold, consisting of an internal and external part, was then 3D printed using the FormLabs 3B+ 3D printer. Subsequently, the mold was coated with non-adhesive ease release 200 spray (Mann Release Technologies, Macungie, PA, USA). The mold was then filled with a mixture of SYLGARD™ 184 Silicone Elastomer Base and Curing Agent, combined at a 10:1 mass ratio. After curing at 80°C for 2 hours, the internal mold was removed, and the cured PDMS was demolded from the external mold, resulting in the vessel phantom in its final form. Before experiment, the PDMS-based phantoms were placed in the plasma cleaner for 5 minutes, cleaned with ethanol for 30 minutes and then dried at 80°C for 30 minutes, which removed the wax residuals.

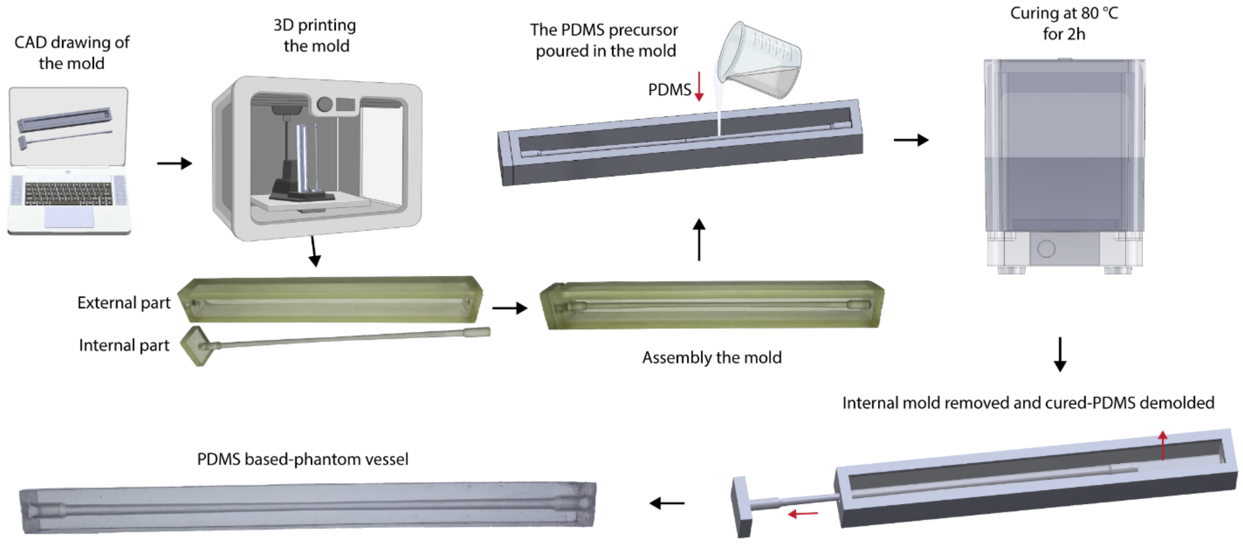

**Figure S11. Fabrication methods and production steps for PDMS-based phantom vessels.**

3.2 *Platinum-cured silicone tubing-based vessel phantom*

To showcase the capabilities of the EndoBot and the magnetic actuator in three dimensions (3D), we constructed a three-dimensional helical vessel labeled as F-labeled vessel phantom in **Table S7**. Given the challenges associated with using PDMS material for creating the 3D vessel phantom in this context, we opted for platinum-cured silicone tubing (16# LG-SIL-16#, Longer Pump, USA) with a diameter of 3.2 mm which also exhibits similar mechanical properties to those of vessels (with a Young's modulus of around 0.2-1.2 MPa (*25*)). This silicone tube was wrapped around a transparent plastic tube (McMaster-Carr, USA) with a diameter of 63.5 mm to form the 3D vessel. The dimensions of the vessel are also detailed in **Table S7**.

### Supplementary Text 4: Friction force modeling

The friction force is given by:

$$\boldsymbol{F}_{\text{friction}} = \boldsymbol{F}_{\text{friction},||} + \boldsymbol{F}_{\text{friction},\perp} = \mu_{||}|\boldsymbol{F}_{\text{radial}}|\boldsymbol{e}_{||} + \mu_{\perp}|\boldsymbol{F}_{\text{radial}}|\boldsymbol{e}_{\perp} \tag{5}$$

where $\mathbf{F}_{\text{friction},||}$ and $\mathbf{F}_{\text{friction},\perp}$ represent the frictional forces parallel and perpendicular to the helix, respectively; $\mathbf{e}_{||}$ and $\mathbf{e}_{\perp}$ are the unit vectors in the parallel and perpendicular directions of the helix, respectively; $\mu_{||}$ and $\mu_{\perp}$ are coefficient of friction (CoF) in the parallel and perpendicular directions, respectively. $\mathbf{F}_{\text{radial}}$ is shown in equation (1). The radial force, $\mathbf{F}_{\text{intrinsic}}$ is a radial force without the effect of magnetic force which was measured by using CellScale UniVert biomaterials testing machine as mentioned in **Fig 2C**. The CoFs were got from(*26*). Depending on the condition of EndoBot, different CoFs will be used either static or kinetic friction friction.

An example of the static friction forces on the optimal $\mathrm{EB3.6}_{0-40\%}^{0\%}$ at different compressive stress with different type of vessel (PDMS-based vessel and human vein vessel) is shown in **Fig S12**. We observed that the intrinsic friction force between the EndoBot and the PDMS-based vessel is approximately twice as high as that between the EndoBot and a human vein vessel. Consequently, the EndoBot exhibits greater conformal deformation capacity in the human vein vessel than in a PDMS-based vessel.

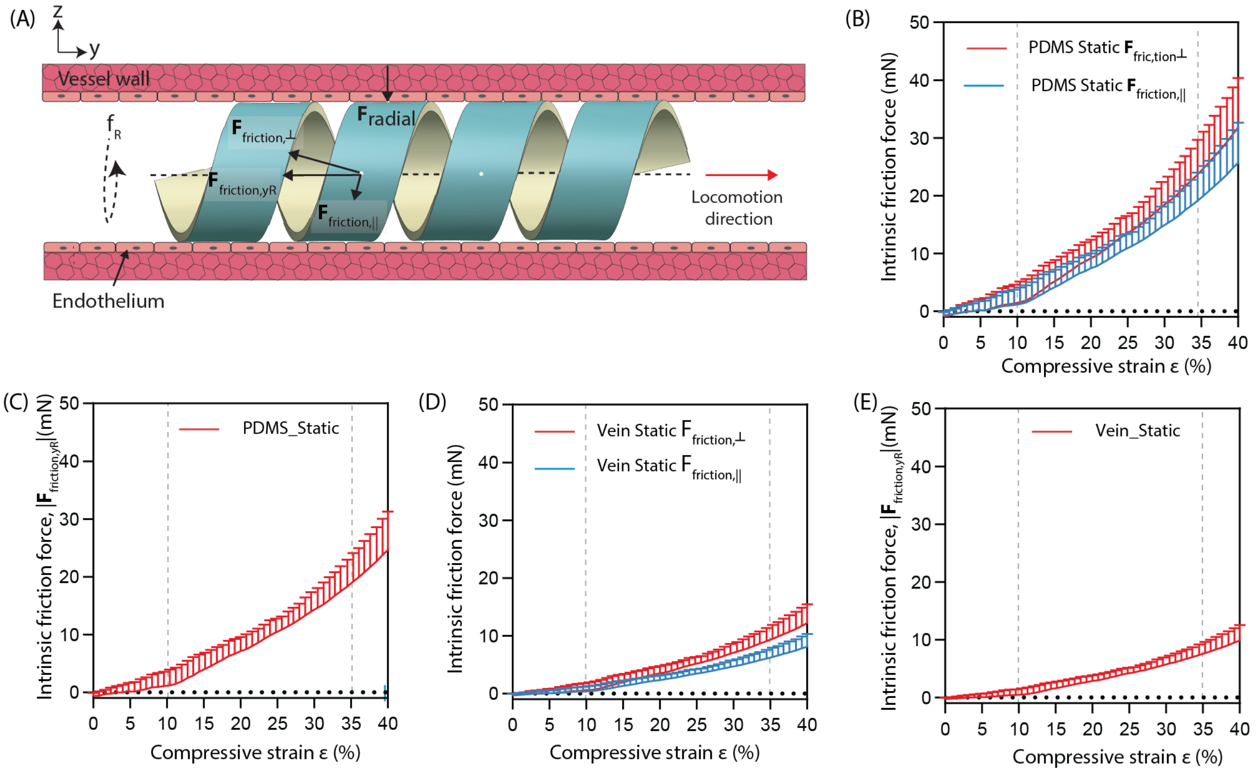

**Figure S12. Intrinsic friction force of the selected $\mathbf{EB3.6_{0-40\%}^{0\%}}$ in different type of vessels.** (A) Description of the intrinsic friction force when the EndoBot inside a vessel. (B) Intrinsic friction force $\mathbf{F}_{friction,||}$ and $\mathbf{F}_{friction,\perp}$ in a PDMS-based vessel. (C) Intrinsic friction force $\mathbf{F}_{friction,yR}$ along the PDMS-based vessel. (D) Intrinsic friction force $\mathbf{F}_{friction,||}$ and $\mathbf{F}_{friction,\perp}$ in a human vein vessel. (E) Intrinsic friction force $\mathbf{F}_{friction,yR}$ along the human vein vessel.

### Supplementary Text 5: Drag force modeling

In this paper, two models were used to predict the drag force $\mathbf{F}_{drag}$.

**Fist model:**

The first model of the drag force was given by the following equation:

$$\mathbf{F}_{drag} = \nabla P \times (S_{lumen} - S_{project}) \quad (6)$$

where ∇P, based on Hagen-Poiseuille Equation (*27*), represents the pressure drop due to viscous blood flow:

$$\nabla P = \frac{8\mu_{blood} L Q}{\pi\left(\frac{D_R - T}{2}\right)^4} \quad (7)$$

where μ is dynamic viscosity of the fluid (Pa·s), $\mu = 4.4\times10^{-3}$ (Pa·s)(*26*).

Q is blood flow rate ($m^3$/s)

*L* is length of EndoBot and will be changed under different lumen vessel (m).

**Second model:**

The $\mathbf{F}_{drag}$ was modeled by using Fluid flow module (Laminar flow analysis) in COMSOL Multiphysics 6.1(*28*).

$$\mathbf{F}_{drag} = \int_S \sigma_{y_R} \, dS \quad (8)$$

where $\sigma_{y_R}$ is the stress in $y_R$ direction, and *dS* is the EndoBot surface.

**Figure 13** shows the drag force under different blood flow rate and different compressive strain ε of both models. Both models produced similar results. Since COMSOL accounts for the actual geometry and the entire surface of the EndoBot, the drag force in the COMSOL model appears to be slightly higher compared to the other model. We found that:

**First model:** $|\mathbf{F}_{drag}| \propto \sim 1/D_R^{3.0}$. **Second model:** $|\mathbf{F}_{drag}| \propto \sim 1/D_R^{3.2}$.

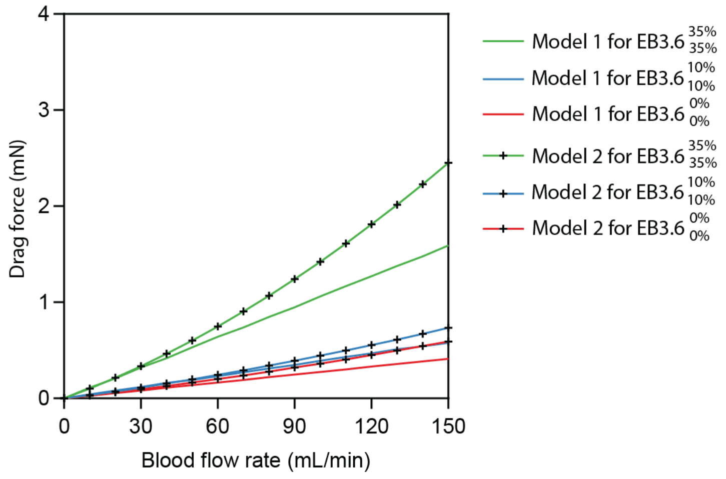

**Figure S13. The drag force of the selected EB3.6 under different compressive strain ε for both models.**

**Supplementary Text 6: System setup for mimicking the anatomical and physiological conditions of blood vessels**

To mimic the anatomical and physiological conditions of blood vessels, with highly tunable flow parameters, we developed a system that shown in **Fig 3A**. This system was constructed by connecting a peristaltic pump (G100-2J, Longer Pump, USA) with platinum-cured silicone tubing (3.2 mm diameter) to a liquid flow sensor (SLFS-4000B, Sensirion AG, Stäf, Switzerland), along with the vessel phantoms outlined in **Table S7**. In vitro experiment, all vessel phantoms were filled with cow blood (registration lot: 45991, Innovative Research, Inc., USA). The blood flow rate of fresh blood using the peristaltic pump at maximum pump capacity is depicted in **Fig S14A**. As the blood flow rate is not constant, the peak flow rate is utilized to assess the performance of EndoBot under fluid flow conditions. Relationship between blood flow rate and blood flow velocity for different $\mathrm{EB3.6}_{10\%}^{10\%}$ and $\mathrm{EB3.6}_{35\%}^{35\%}$ are shown in **Fig S14B**. Note that, blood flow velocity was measured at the center of the vessel where EB was present as shown in **Fig S14B**.

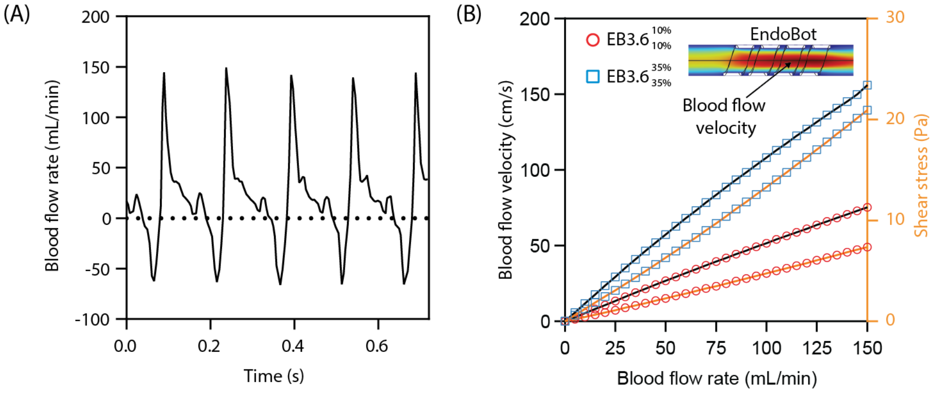

**Figure S14. Characterization of blood flow generated by the peristaltic blood pump.** (A) Blood flow behavior of pump. (B) Relationship between blood flow rate and blood flow velocity for different $\mathrm{EB3.6}_{10\%}^{10\%}$ and $\mathrm{EB3.6}_{35\%}^{35\%}$ The blood flow velocity was measured at the center of the vessel, where EB was present.

### Supplementary Text 7: Magnetic control and magnetic force and torque modeling

An LBR Med 7 R800 robot arm (KUKA) with seven degrees of freedom was employed to control the magnetic field position and orientation in the workspace of the EndoBot, allowing precise control over the direction and magnitude of the magnetic fields from a mechanically safe position **($\mathbf{r}_{mR}$)**. Our KUKA robot arm's software version is V1.5.4-2 and KUKA Robot arm programmed with its unique application (KUKA Sunrise Workbench – 2.6.5_6). Attached to the robot arm end effector, a custom-designed DC motor (EC- I30, maxon precision motors, Taunton, MA, USA)-run part facilitates the high-performance rotation of an external permanent magnet (50 mm × 50 mm cylinder permanent magnet, N52, K&J Magnetics, Inc., USA) up to ~100 Hz. The speed of the servo motor was controlled by using ESCON 70/10, and variable resistor of 10 kΩ. The servo motor was supplied by a DC Power (Tek Power TP610E (60V, 10A)). The magnet mounted-robot arm is depicted in **Fig 1A** and **Fig 6A**.

To facilitate tracking and explanation in this paper, local coordinate systems were assigned to both the magnet and EndoBot. The magnet's local coordinate system ($x_m$-$y_m$-$z_m$) was centered on the magnet, while EndoBot's local coordinate system ($x_R$-$y_R$-$z_R$) was centered on EndoBot (**Fig 1D)**. When the magnet, characterized by its magnetic dipole moment ($\mathbf{M}_m$), rotates around the $y_m$-axis, it induces EndoBot, located at $\mathbf{r}_{mR}$ and possessing its own magnetic moment ($\mathbf{M}_R$), to rotate around the $y_R$-axis in the opposite direction. During EndoBot's navigation along an arbitrary three-dimensional trajectory, the magnet translates at a velocity of $\mathbf{v}_m$, while EndoBot follows predefined trajectories by adjusting its position ($d_x$, $d_y$, $d_z$) along the $x_m$, $y_m$, and $z_m$ axes as shown in Fig 1D. The $y_m$-axis of the magnet's local coordinate system is used to orient the magnet relative to EndoBot and is consistently aligned tangentially to the vessel's centerline, ensuring effective torque transmission. To ensure safety, the magnet is positioned at least 52 mm from its center along the $z_m$-axis ($|d_z| \geq 52$ mm), enabling the delivery of magnetic fields ranging from 4-145 mT to depths up to $|d_z|$=120 mm within the human body.

The magnetic force on the EndoBot was shown in equation (2) and the magnetic torque on EndoBot is given by:

$$\boldsymbol{T}_{\text{mag}}(\boldsymbol{r}_{\text{mR}}) = \boldsymbol{M}_{\text{R}} \times \mathbf{B}(\boldsymbol{r}_{\text{mR}}) \tag{8}$$

where the magnetic field was modeled by using AC/DC magnetic field module in COMSOL Multiphysics 6.1 and validated by using DC Gauss-meter (Model GM1-ST; AlphaLab, Inc., Salt Lake City, UT). The measurements agree well with the simulation results (**Fig S15**). Since the magnet is rotated during the experiment, the magnetic field at the EndoBot's position is not constant. Examples of the changing magnetic field when the magnet is rotated around the $y_m$-axis from 0° to 360° at several positions are shown in **Fig S16.** **|$M_R$|** of EndoBot was get based on the magnetization curve of NdFeB D50 in (*29*), the magnetic moment values of each EndoBot were shown in **Table S6.**

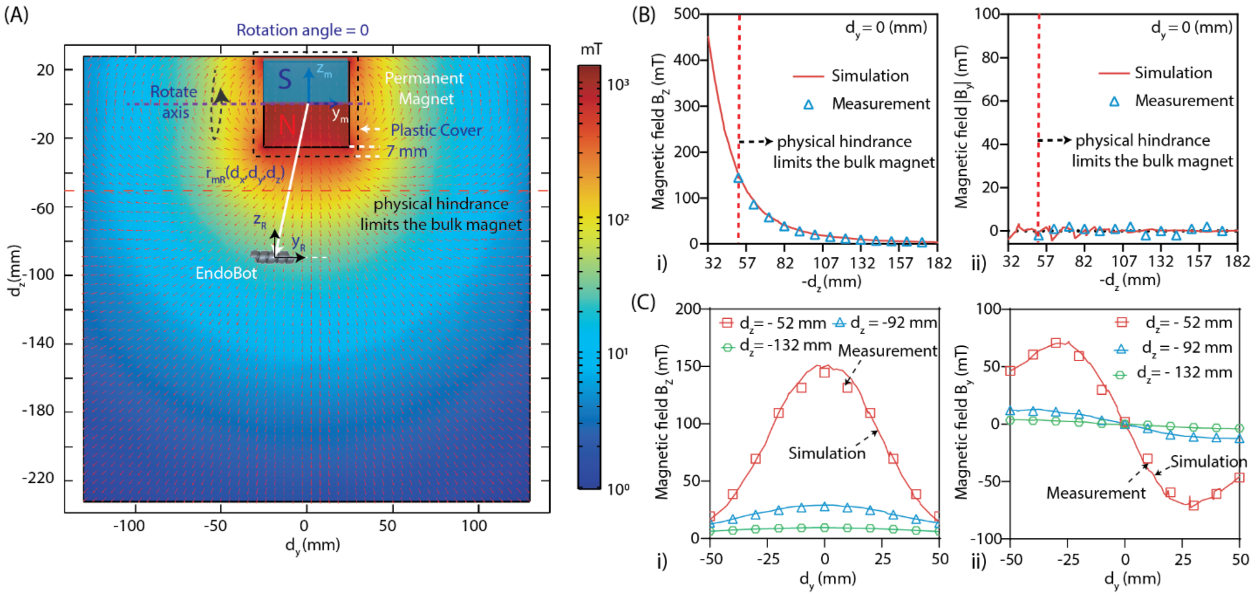

**Figure S15. Magnetic field lines around EndoBot when the permanent magnet is on standby.** (A) Snapshot of the magnetic field lines on the yz-plane with the magnet fixed at a rotation angle of 0° (**r**$_{mR}$($d_x$, $d_y$, $d_z$) represents the position of the EndoBot's center in $x_m$,$y_m$,$z_m$ coordinate system). (B) Comparison of the magnetic field components $B_y$ and $B_z$, where $B_y$ and $B_z$ represent the magnetic field in the y- and z-directions, respectively. (C) Comparison of the magnetic field components $B_y$ and $B_z$ at varying $d_y$ with $d_z$ fixed at -52, -92, and -132 mm. When the external magnet and EndoBot are centrally aligned, torque is maximized, resulting in the most efficient propulsion of the robot. To maintain these optimal conditions, the robot arm must be continuously repositioned and reoriented as EndoBot moves, guided by continuous visual feedback. If EndoBot accelerates along the $y_m$- axis, for example, due to external flow or suboptimal response from the robot arm carrying the magnet, a corrective pulling magnetic field $B_y$ realigns EndoBot.

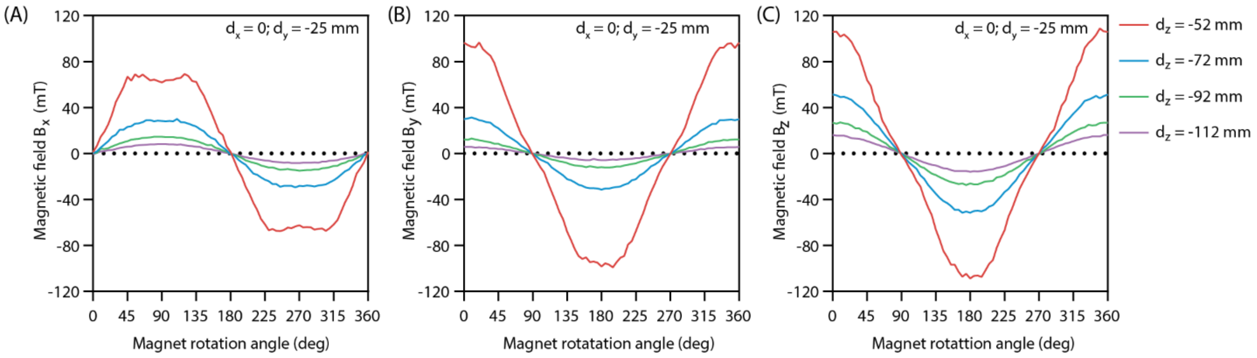

**Figure S16. Magnetic field distribution of $B_x$, $B_y$ and $B_z$, where $B_x$, $B_y$ and $B_z$ are magnetic field components of $B(r_{mR})$ in x-,y- and z-directions, respectively, at various positions when magnet is rotated around $y_m$-axis from 0° to 360$^0$.** (A) Magnetic field x-component $B_x$. (B) Magnetic field y-component $B_y$, and (C) Magnetic field z-component $B_z$.

The absolute magnetic forces $|\mathbf{F}_{mag,zR}|$ by COMSOL simulation with different $|d_z|$ or magnetic field were validated by using a customer design combine with a precision balance (BCE822-1S, Sartorius Lab Instruments GmbH & Co. KG, 37070 Goettingen, Germany) as shown in **Fig S17A**. The magnetic force measurements closely match to the simulation results (**Fig S17B**).

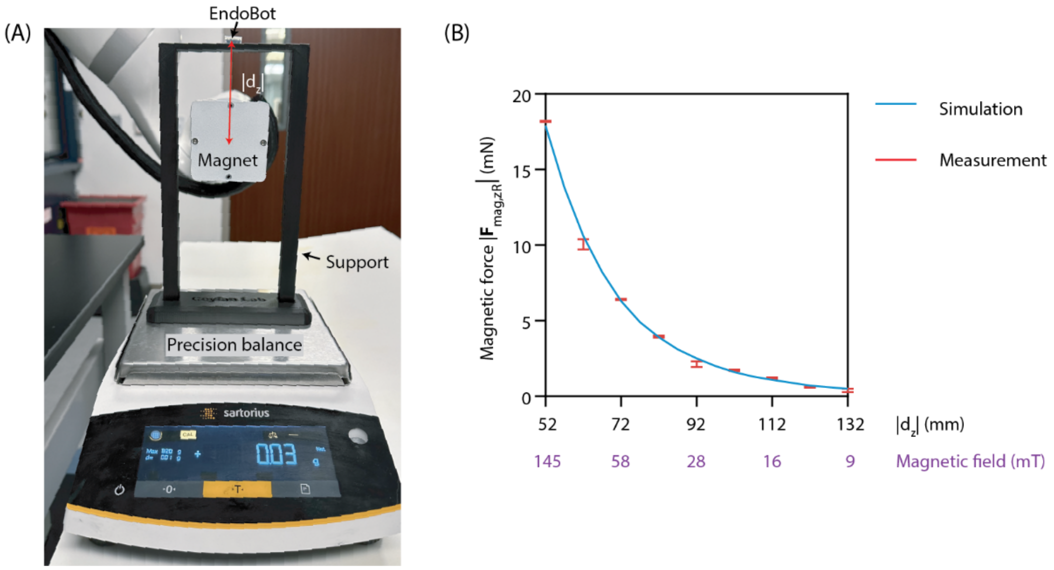

**Figure S17. Magnetic Force Evaluation.** (A) Customer-designed setup for measuring magnetic force on EndoBot EB3.6 with different magnetic field or $|d_z|$, using a precision balance. (B) Absolute magnetic force $|\mathbf{F}_{mag,z}|$ along $z_m$ axis, demonstrating a close match between measured values and simulation results.

The maximum absolute force and torque of EndoBot with variable $|d_z|$ and $|d_y|$ are shown in **Fig S18**.

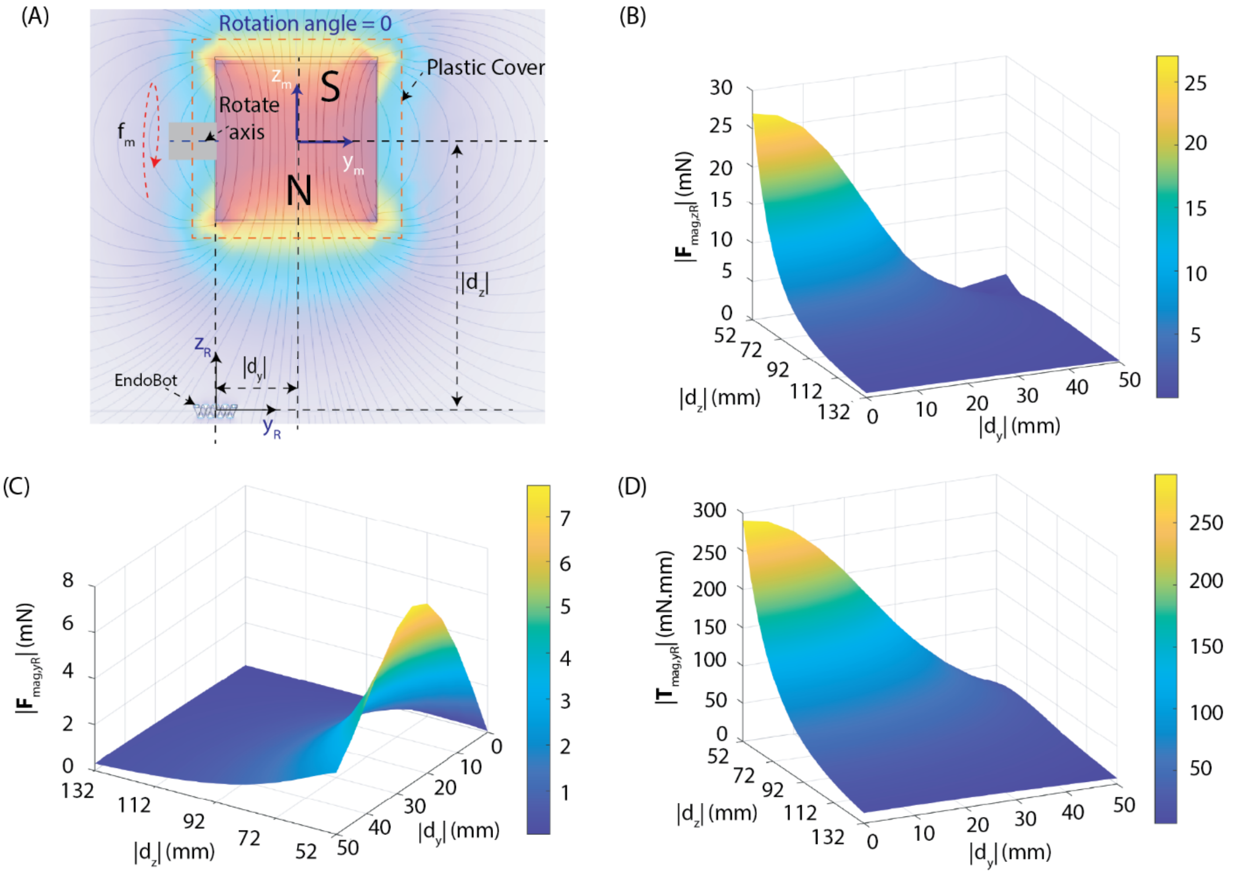

**Figure S18. Maximization of stabilizer force under flow and magnetic torque for propulsion.** Maximum achieved absolute magnetic forces and torques with different $|d_y|$ and $|d_z|$. (A) Definition of $|d_y|$ and $|d_z|$ (Note that $d_x$, $d_y$, $d_z$ are components of $\mathbf{r}_{mR}$ which represents the position of the EndoBot's center in $x_m$,$y_m$,$z_m$ coordinate system). (B) Maximum absolute magnetic retraction force $|\mathbf{F}_{mag,zR}|$ in $z_R$ direction. (C) Maximum absolute magnetic retraction force $|\mathbf{F}_{mag,yR}|$ in $y_R$ direction. (D) Maximum absolute magnetic torque $|\mathbf{T}_{mag,yR}|$ in $y_R$ direction.

To achieve highest locomotion performance for EndoBot, the absolute magnetic torque and force should be maximized. From the results in **Fig S18**, we can define propulsion optimization as follow:

- To maximize the absolute magnetic torque $|\mathbf{T}_{mag,yR}|$ in $y_R$ direction and to maximize the absolute magnetic force $|\mathbf{F}_{mag,yR}|$ in $z_R$ direction, $|d_z|$ should be minimized for a given $|d_y|$.
- For a fixed $|d_z|$, the maximum absolute magnetic force $\mathbf{F}_{mag,yR}$ in $y_R$ direction is achieved when $|d_y|$ is close to 25 mm. The value of $d_y$ depends on the desired movement direction of the EndoBot. For example, to move the EndoBot from left to right ($y_m$ direction), set $d_y$ close to -25 mm and

move the magnet in the same direction as the movement. Similarly, to move the EndoBot from right to left, set $d_y$ close to +25 mm and move the magnet in the corresponding direction.

- The rotation of the EndoBot must adapt to both the strength and direction of blood flow to ensure effective movement.

  i) When the frictional force dominates over the drag force (Such as in case without/small blood flow and EndoBot under high compression), in addition to defining $d_y$ and magnet movement direction as previously explained, the EndoBot's rotation determines the propulsion direction. To move along the blood flow direction (left to right, as shown in **Fig S19A**), the EndoBot should rotate counterclockwise. In this paper, the rotation direction is always considered as viewed in the positive $y_m$-axis direction, with the coordinate system defined in **Fig 1D**. Conversely, to move against the blood flow direction (right to left, as shown in **Fig S19A**), the EndoBot should rotate clockwise.

  ii) When the drag force becomes comparable to or greater than the frictional force, the EndoBot's rotation should always remain clockwise to maintain control and stability. In this case, the direction of movement is determined solely by adjusting $d_y$ and magnet movement direction as previously described.

To maintain these optimal conditions, the robot arm must be continuously repositioned and reoriented as EndoBot moves, guided by continuous visual feedback.

An example of the EndoBot's magnetic actuation mechanism is shown in **Fig S19**. The maximum velocity of the EndoBot is achieved when moving forward from $d_y$=25 mm to $d_y$=0 mm, as illustrated in **Fig S19C**. In detail, the maximum speed of the EndoBot $\mathrm{EB3.6}_{10\%}^{10\%}$ within a 3.2 mm diameter vessel is 14.3 ± 2.9 mm/s, achieved at 30 Hz, $d_y$ = ~ 25 mm and $d_z$ = -52 mm (magnetic field ~145 mT), while the maximum speed of $\mathrm{EB3.6}_{35\%}^{35\%}$ within a 2.3 mm diameter vessel is 14.5 ± 7.9 mm/s, also achieved at 30 Hz, $d_y$ = ~ 25 mm and $d_z$ = -52 mm. No step-out frequency was observed under the magnetic actuation conditions prescribed in this study.

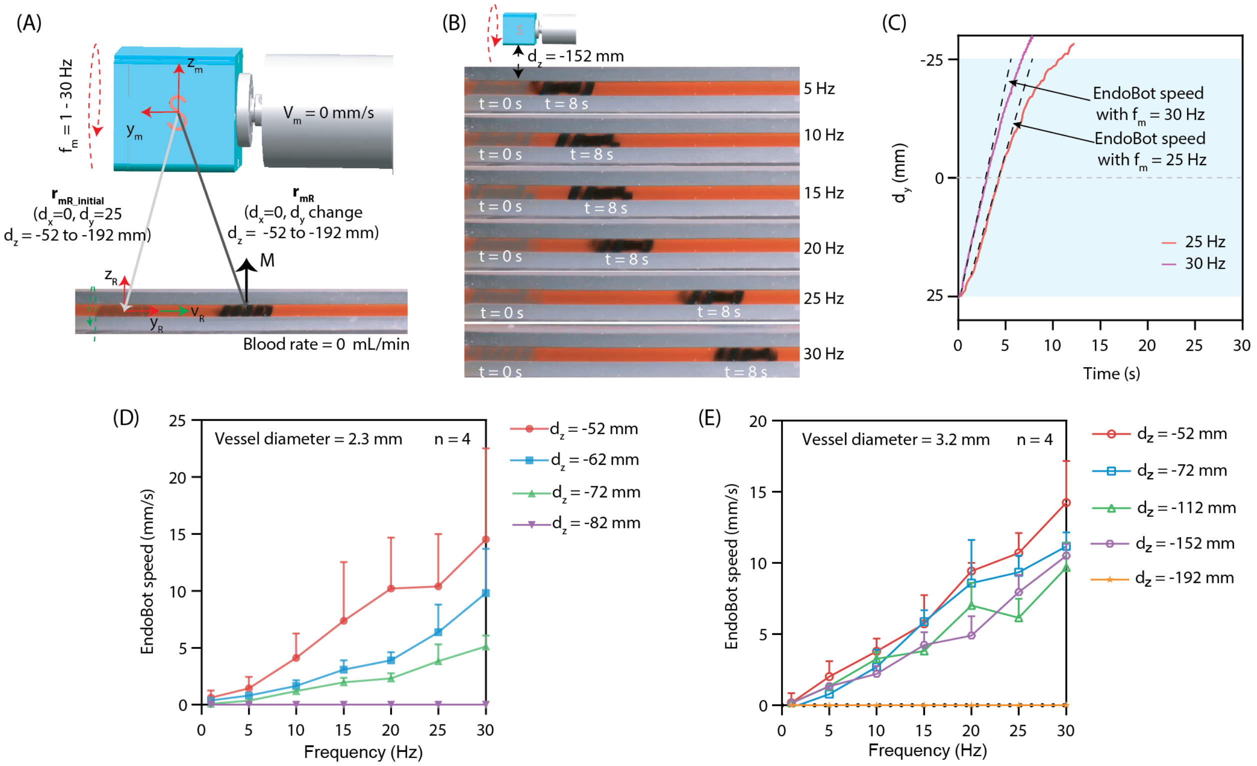

**Figure S19**. **EndoBot magnetic actuation mechanism.** The external magnet is kept at a certain vertical distance, $|d_z|$ and an initial horizontal distance, $|d_y|$ (where $d_x$, $d_y$, $d_z$ are components of $\mathbf{r}_{mR}$ which represents the position of the EndoBot's center in $x_m$,$y_m$,$z_m$ coordinate system). Since EndoBot is confined in a straight-line vessel, $d_x$ remained constant during the experiment. (A) Description of experimental condition. (B) An example of an EndoBot $\mathrm{EB3.6}_{10\%}^{10\%}$ moving within a 3.2 mm diameter vessel channel under different frequencies ranging from 5 to 30 Hz. (C) Definition of EndoBot speed. The speed results of (D) the EndoBot $\mathrm{EB3.6}_{35\%}^{35\%}$moving within vessel diameters of 2.3 mm and (E) the EndoBot $\mathrm{EB3.6}_{10\%}^{10\%}$ moving within vessel diameters of 3.2 mm.

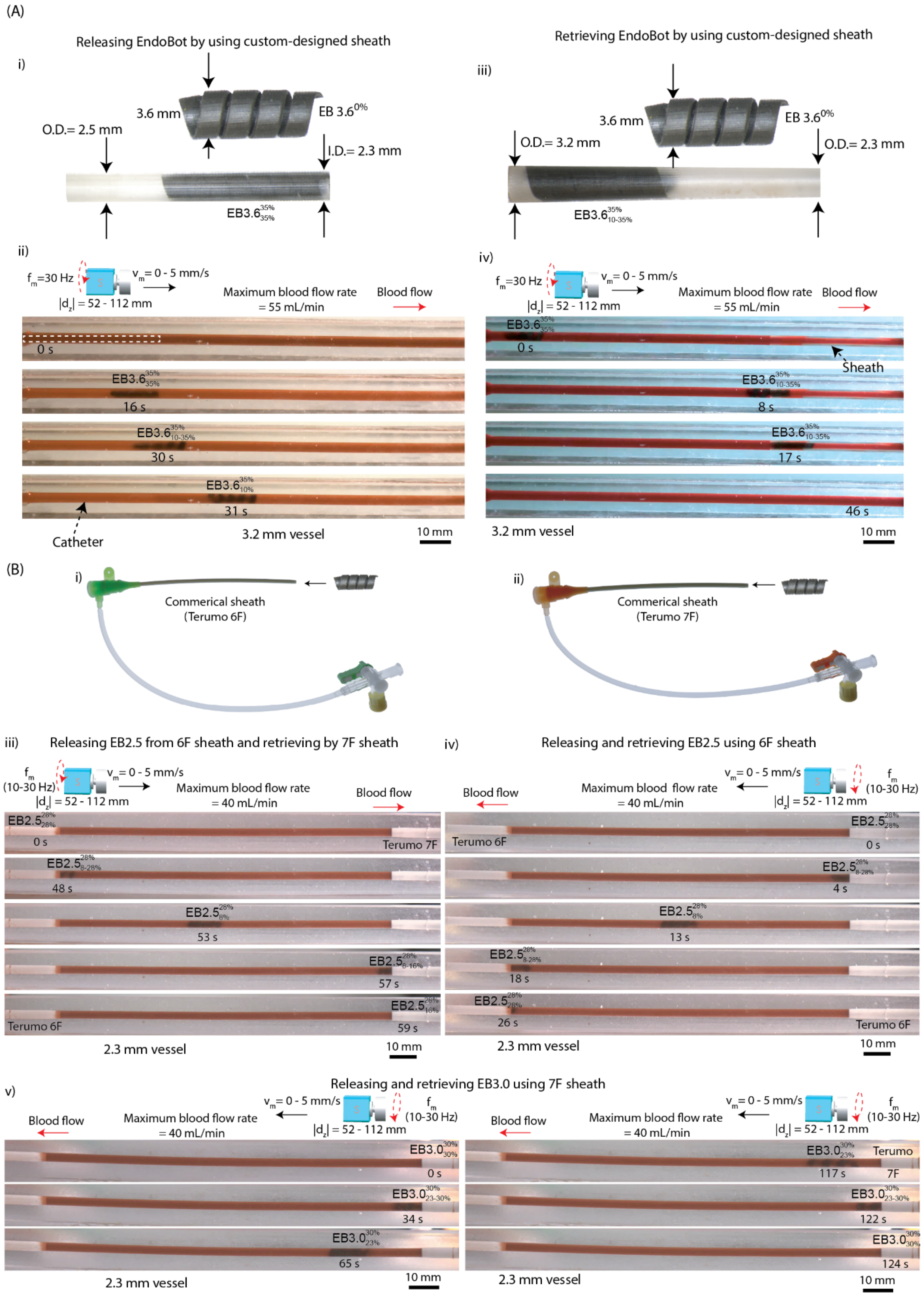
(A)
Releasing EndoBot by using custom-designed sheath
i)
3.6 mm
$EB\ 3.6^{0\%}$
O.D.= 2.5 mm
I.D.= 2.3 mm
$EB3.6^{35\%}_{35\%}$
Retrieving EndoBot by using custom-designed sheath
iii)
3.6 mm
$EB\ 3.6^{0\%}$
O.D.= 3.2 mm
O.D.= 2.3 mm
$EB3.6^{35\%}_{10-35\%}$
ii)
$f_m$=30 Hz
$v_m$ = 0 - 5 mm/s
$|d_z|$ = 52 - 112 mm
Maximum blood flow rate = 55 mL/min
Blood flow
0 s
$EB3.6^{35\%}_{35\%}$
16 s
$EB3.6^{35\%}_{10-35\%}$
30 s
$EB3.6^{35\%}_{10\%}$
31 s
Catheter
3.2 mm vessel
10 mm
iv)
$f_m$=30 Hz
$v_m$ = 0 - 5 mm/s
$|d_z|$ = 52 - 112 mm
Maximum blood flow rate = 55 mL/min
Blood flow
$EB3.6^{35\%}_{35\%}$
0 s
Sheath
$EB3.6^{35\%}_{10-35\%}$
8 s
$EB3.6^{35\%}_{10-35\%}$
17 s
46 s
3.2 mm vessel
10 mm
(B)
i)
Commerical sheath (Terumo 6F)
ii)
Commerical sheath (Terumo 7F)
iii) Releasing EB2.5 from 6F sheath and retrieving by 7F sheath
$f_m$ (10-30 Hz)
$v_m$ = 0 - 5 mm/s
$|d_z|$ = 52 - 112 mm
Maximum blood flow rate = 40 mL/min
Blood flow
$EB2.5^{28\%}_{28\%}$
0 s
Terumo 7F
$EB2.5^{28\%}_{8-28\%}$
48 s
$EB2.5^{28\%}_{8\%}$
53 s
$EB2.5^{28\%}_{8-16\%}$
57 s
$EB2.5^{28\%}_{16\%}$
59 s
Terumo 6F
2.3 mm vessel
10 mm
iv) Releasing and retrieving EB2.5 using 6F sheath
Blood flow
Maximum blood flow rate = 40 mL/min
$v_m$ = 0 - 5 mm/s
$|d_z|$ = 52 - 112 mm
$f_m$ (10-30 Hz)
Terumo 6F
$EB2.5^{28\%}_{28\%}$
0 s
$EB2.5^{28\%}_{8-28\%}$
4 s
$EB2.5^{28\%}_{8\%}$
13 s
$EB2.5^{28\%}_{8-28\%}$
18 s
$EB2.5^{28\%}_{28\%}$
26 s
Terumo 6F
2.3 mm vessel
10 mm
v) Releasing and retrieving EB3.0 using 7F sheath
Blood flow
Maximum blood flow rate = 40 mL/min
$v_m$ = 0 - 5 mm/s
$|d_z|$ = 52 - 112 mm
$f_m$ (10-30 Hz)
$EB3.0^{30\%}_{30\%}$
0 s
$EB3.0^{30\%}_{23-30\%}$
34 s
$EB3.0^{30\%}_{23\%}$
65 s
2.3 mm vessel
10 mm
$EB3.0^{30\%}_{23\%}$
Terumo 7F
117 s
$EB3.0^{30\%}_{23-30\%}$
122 s
$EB3.0^{30\%}_{30\%}$
124 s
2.3 mm vessel
10 mm

**Figure S20. Wireless deployment and release of EndoBot.** (A) Using custom-designed sheath: (i) Releasing EndoBot $\mathrm{EB3.6}_{10-35\%}^{35\%}$, (ii) Retrieving EndoBot $\mathrm{EB3.6}_{10-35\%}^{35\%}$. (B) Using commercial sheaths: (i) Terumo 6F sheath, (ii) Terumo 7F sheath, (iii) Releasing $\mathrm{EB2.5}_{8-28\%}^{28\%}$ from the 6F sheath and retrieving with the 7F sheath, (iv) Releasing $\mathrm{EB2.5}_{8-28\%}^{28\%}$ from the 6F sheath and retrieving with the same 6F sheath, (v) Releasing $\mathrm{EB3.0}_{23-30\%}^{30\%}$ from the Terumo 7F sheath and retrieving with the same sheath.

**Supplementary Text 8: Biocompatibility assessment of EndoBot**

As a blood-contacting device, EndoBot and its corkscrew locomotion must result in minimal adverse interactions with blood and endothelial cells (ECs) lining the vessel intima. Although the thrombogenicity of such devices is difficult to predict, we conducted a comprehensive series of tests to evaluate coagulative properties of EndoBots, hemolytic potential, platelet activation, and cytotoxicity. These tests followed commonly accepted in vitro protocols aligned with ISO-10993-Part 4 (Biological evaluation of medical devices: Selection of tests for interactions with blood) guidelines and customized procedures tailored to our application context(*36, 37*).

*8.1 Evaluate the effect of blood on EndoBot's locomotion*

The envisioned operational timescale of EndoBot within a blood vessel is short-term (<15 min). During this period, its surface material composition and locomotion could potentially trigger blood coagulation. To assess this risk, we continuously monitored the real-time translational velocity of $\mathrm{EB3.6}_{10\%}^{10\%}$ moving in a phantom vessel filled with whole bovine blood under constant magnetic actuation (~12-15 mT, 10 Hz) for 30 min (**Fig 5A**). In detail, the $\mathrm{EB3.6}_{10\%}^{10\%}$ was moved from left to right, the magnet and the $\mathrm{EB3.6}^{10\%}$ was positioned at $\mathbf{r}_{mR}$ of $d_x$ = 0 mm, $d_y$ = -80 mm, and $d_z$ = -112 mm at a clockwise rotation frequency of $f_m$ =10 Hz, without using a movement magnet (referred to as a forward lap). Similarly, to move the $\mathrm{EB3.6}^{10\%}$ from right to left, the magnet and the $\mathrm{EB3.6}_{10\%}^{10\%}$ was positioned at $\mathbf{r}_{mR}$ coordinates of $d_x$ = 0 mm, $d_y$ = 80 mm, and $d_z$ = -112 mm with a counterclockwise rotation frequency of $f_m$ =10 Hz, also without a movement magnet (referred to as a backward lap). The blood was pre-heparinized at 0.5 U/mL, approximately matching the prophylactic dose recommended for adults undergoing endovascular procedures for deep vein thrombosis(*38*). A decrease in the robot velocity over ~40 forward-backward traversals (~560 cm total distance) would suggest coagulation-induced drag. However, mean velocities remained stable (5.9 ± 0.9 mm/s forward and 5.9 ± 0.8 mm/s backward), demonstrating no signs of coagulation-induced drag. This key observation provides translatable evidence for future animal and human procedures.

*8.2 Coagulation*

Before assessing potential coagulation risks, EndoBot samples were first cleaned with alcohol and phosphate-buffered saline (PBS), then placed in a plasma cleaner for 30 minutes. After cleaning, the samples were coated with tributyl O-acetylcitrate. The coated surfaces were then treated with three different methods over 24 hours:

- Positive Treatment Method: The coated EndoBots were treated with collagen (1 mg/mL of gelatin in PBS) and are referred as Collagen-treated EndoBot.

- Negative Treatment Method: The coated EndoBots were treated with 1 mL heparin (10000 units/10 mL) and are referred as Heparin-treated EndoBot.
- No Treatment Method: The coated EndoBots were placed in 1 mL distilled water (dH2O) and are referred as Untreated EndoBot.

The surface-treated EndoBots were placed in low protein binding microcentrifuge tubes, with each tube containing 900 µL of fresh human blood (registration lot: ND0809-49877 and ND0337-49876 from Innovative Research, Inc., USA). The blood was exposed to these surface-treated EndoBot groups overnight before measuring the activated clotting time (ACT). To reduce the ACT time of the pre-heparinized single-donor human blood, whole blood was initiated by adding calcium chloride ($CaCl_2$) solution. We used two different concentrations of $CaCl_2$: Blood volume ratios for this test a 1:50 v/v ratio (3.6 mM $CaCl_2$) and 1:65 v/v ratio of 0.2 M calcium chloride ($CaCl_2$) solution (2.8 mM $CaCl_2$). Whole blood samples, with and without $CaCl_2$, were used as control and negative control samples, respectively. Upon adding the $CaCl_2$, the blood was used immediately for the ACT test. The ACT of the samples was recorded using an i-STAT blood analyzer (Abbott Laboratories, Chicago, IL) and a Celite ACT cartridge (Abbott Laboratories, Chicago, IL). Under these conditions, ACT values for the untreated EndoBot were comparable to both controls, indicating no heightened or reduced procoagulant effect.

Additionally, a complete blood count (CBC) test (HESKA CBC) was performed to measure hematological parameters, including hemoglobin, red blood cells (RBCs), hematocrit, platelet count, white blood cell count, and differential leukocyte counts. To conduct this test, each surface-treated EndoBot was placed in 1 mL of human blood (registration lots ND0752-49725 and ND0752-49723 from Innovative Research, Inc., USA). A control sample consisting of 1 mL of blood without an EndoBot was also prepared. After 30 minutes of incubation, the blood samples were analyzed using the CBC test. The results, shown in **Fig S21**, indicate that all measured hematological parameters remained within normal ranges, with no adverse changes in blood cell morphology or counts. Furthermore, the absence of detectable pro-coagulant behavior is consistent with the U.S. National Heart, Lung, and Blood Institute's designation of silica filler-free PDMS as a reference material for blood compatibility(*39*).

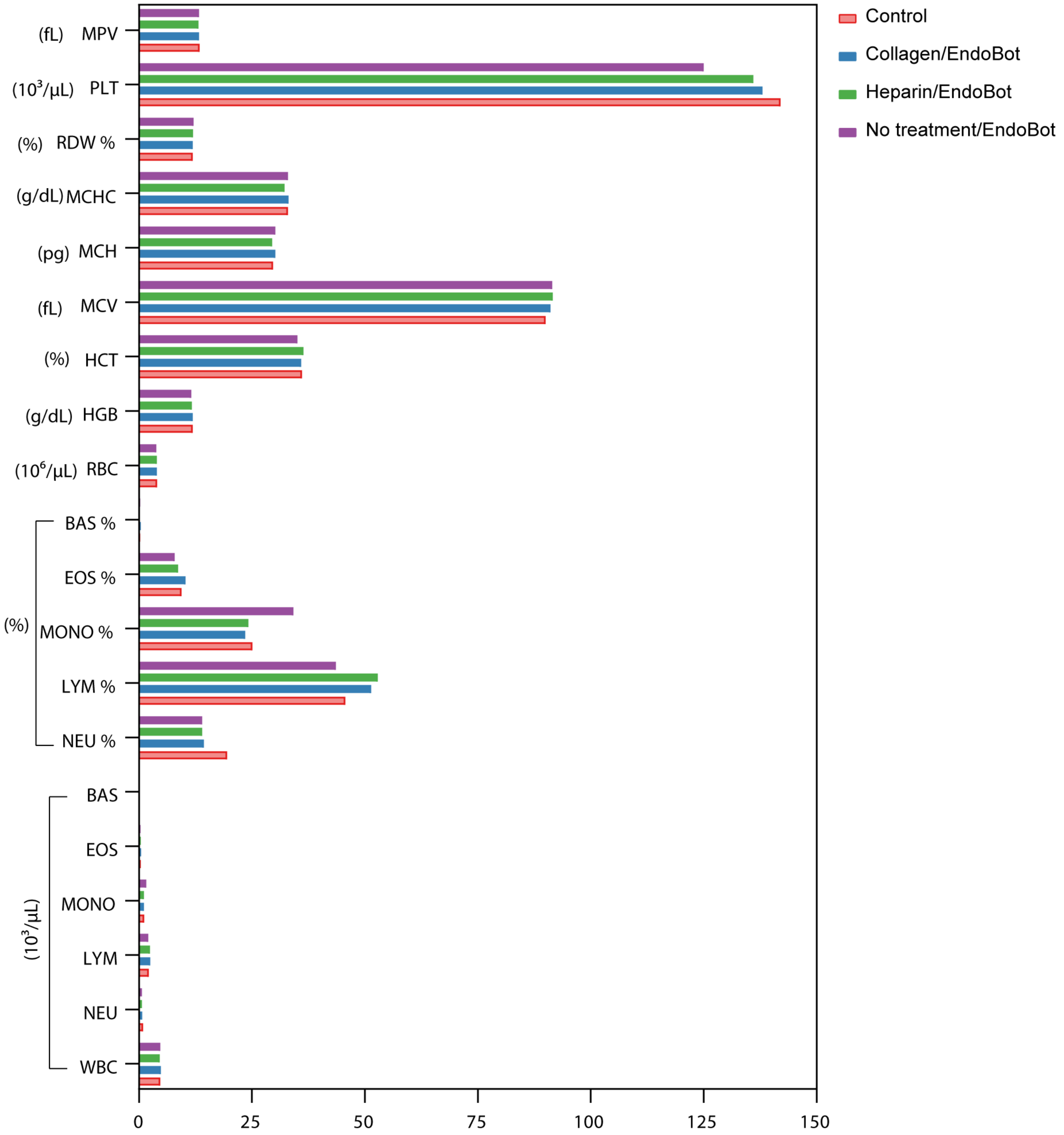

**Figure S21. Complete blood count of human blood after exposure to overnight surface-treated EndoBots.**

*8.3 Hemolysis studies*

Hemolysis, the rupture of RBCs and subsequent release of their intracellular components, can be induced by mechanical stress or surface interactions. Such released molecules may, in turn, trigger thrombogenesis by activating platelets(*40*).

To evaluate potential hemolytic effects, the first hemolysis study focused on the impact of locomotion. EndoBot samples were thoroughly cleaned with alcohol and phosphate-buffered saline (PBS) before being treated in a plasma cleaner for 30 minutes. After cleaning, the samples were coated with tributyl O-acetylcitrate, which serves as a model for a drug excipient coating layer. To evaluate the effect of locomotion frequency on hemolysis, we quantified free hemoglobin released from RBCs after exposing fresh, heparinized bovine blood (registration lot: 48507 from Innovative Research, Inc., USA) to $\mathrm{EB3.6}_{10\%}^{10\%}$ in a straight vessel phantom at varying locomotion frequencies (1, 10, and 30 Hz) for 15 min. The vessel phantom used in this experiment was a 20 cm long platinum-cured silicone tubing, containing 1.6 mL of blood. We prepared three EndoBots for each locomotion group. The experimental setup is illustrated in **Fig S22**. A negative control was prepared using a vessel phantom without EndoBot movement. After the experiment, 1 mL blood samples were collected from the four groups' vessel phantoms into low protein binding microcentrifuge tubes (Thermo Scientific, USA). These tubes were agitated for 20 minutes at 1500 revolutions per minute (rpm) at 4°C temperature to separate the red blood cells. Following the separation, 20 µL from each sample group was used for hemolysis analysis using a hemoglobin assay kit (colorimetric) (ab234046). For the positive control in the hemolysis analysis, 1 mL of fresh cow blood was mixed with 9 mL of distilled water (dH2O) in a 1:10 volume ratio to fully activate the hemoglobin in the blood, then 20 µL of this mixture was used with the hemoglobin assay kit. According to the kit's protocol, the 20 µL samples from all five groups were mixed with the hemoglobin detector, and the hemoglobin signal at 575 nm was recorded. The hemoglobin concentration was calculated based on the standard curve provided with the kit. The hemolysis rate was then calculated using the following equation:

$$\text{Hemolysis rate } (\%/\mathrm{L}) = \frac{H_{sample,1.6\text{ mL}} - H_{neg,1.6\text{ mL}}}{H_{pos,1\mathrm{L}}} \times 100\% \quad (17)$$

Where $H_{\text{sample, 1.6 mL}}$ is the amount of activated hemoglobin in the 1.6 mL blood sample exposed to the EndoBot, $H_{\text{neg, 1.6 mL}}$ is the amount of activated hemoglobin in the 1.6 mL blood sample not exposed to the EndoBot., and $H_{\text{pos, 1L}}$ is the amount of activated hemoglobin in 1 L of blood used as the positive control.

Since this experiment was conducted without blood flow and the $\mathrm{EB3.6}_{10\%}^{10\%}$ moved forward and backward multiple times, the red blood cells in the vessel channel were repeatedly exposed to the EndoBot's locomotion. This setup represents a scenario of maximum potential hemolysis. In real conditions with blood flow, during the 15 minutes of $\mathrm{EB3.6}^{10\%}_{10\%}$ operation, individual red blood cells would be exposed to the EndoBot's locomotion only briefly before being carried away by the blood flow. Therefore, the hemolysis percentage in this study is compared to a 1 L volume of blood, rather than the blood volume present in the vessel channel used in the experiment.

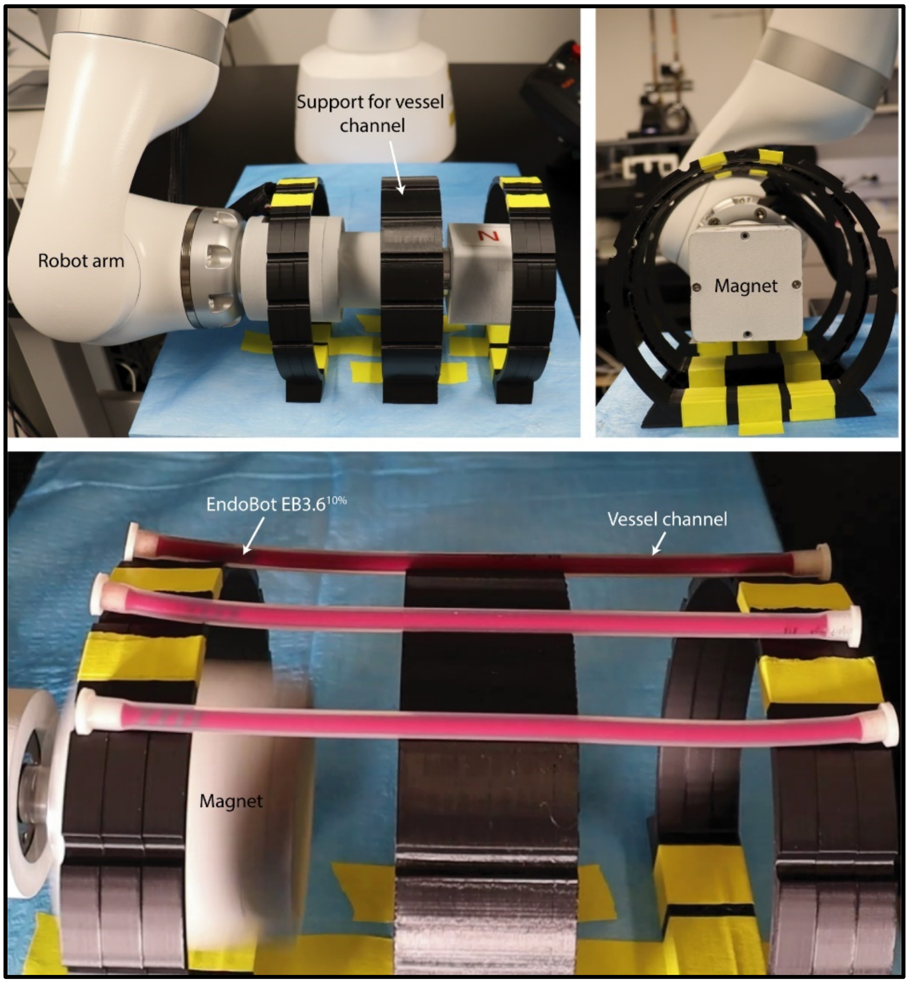

**Figure S22. Experimental setup for the blood compatibility experiments described in Figs 5C, D, and E.** A custom-designed support was used to position three platinum-cured silicone tubing-based vessel phantoms, enabling the simultaneous movement of three $\mathrm{EB3.6}^{10\%}_{10\%}$ samples under similar magnetic field conditions, achieving n = 3 for sufficient replication of the results.

The hemolysis results are shown in **Fig 5C**. The amount of activated hemoglobin at 30 Hz is 1.5 times more than that at 10 Hz and over 18 times that at 1 Hz. Higher magnetic actuation frequencies correlated with increased hemolysis, which we attribute to intensified mechanical shear forces at the blood-robot interface. However, when compared to the total amount of hemoglobin in 1 liter of blood, the activated hemoglobin at 30 Hz is significantly smaller (0.018%

compared to 100%). Therefore, we can conclude that the locomotion of EndoBot does not adversely affect the blood.

In the second study, we evaluated the effects of different treated EndoBot samples under locomotion on hemolysis. The EndoBot samples underwent the same cleaning protocol described in the first study. After cleaning, the samples were coated with tributyl O-acetylcitrate and treated using three different methods, referred to as Collagen-treated EndoBot, Heparin-treated EndoBot, and Untreated EndoBot, over a 24-hour period, as detailed in **Supplementary Text 8.2**. The treated EndoBot ($\mathrm{EB3.6}_{10\%}^{10\%}$) samples were then moved inside a heparin-treated straight vessel phantom filled with human blood (registration lot: ND0694-48858, Innovative Research, Inc., USA) for 15 minutes at a locomotion frequency of 30 Hz. The vessel phantom, as in the first experiment, consisted of a 20 cm long platinum-cured silicone tubing containing 1.6 mL of blood. Three EndoBot samples were prepared for each treatment group. Negative and positive controls for human blood, following the same protocol as in the first hemolysis study, were also included. Blood samples (1 mL each) were collected into 1.5 mL low protein-binding microcentrifuge tubes, agitated for 20 minutes at 1500 rpm at 4°C to separate red blood cells. Hemolysis analysis was performed using the same hemoglobin assay kit (colorimetric) (ab234046), and the hemolysis rate was calculated using equation 17. All EndoBot samples from the second hemolysis study were subsequently collected for thrombogenicity (platelet activation) studies. The hemolysis results, shown in **Fig 5D**, revealed that Collagen-treated EndoBot samples exhibited over double the amount of activated hemoglobin compared to Heparin-treated EndoBots. The activated hemoglobin levels for Heparin-treated EndoBots were similar to those for Untreated EndoBots. When compared to the total hemoglobin in 1 liter of blood, the activated hemoglobin with Collagen treatment remained significantly low (0.054% compared to 100%). These findings indicate that surface treatments influenced hemolytic outcomes: Heparin-treated EndoBots produced slightly lower levels of hemolysis compared to untreated devices, while Collagen-treated EndoBots exhibited the highest hemolysis (**Fig 5D**). However, according to ASTM F756-17 (Standard Practice for Assessment of Hemolytic Properties of Materials), materials are classified as non-hemolytic if their hemolysis rate remains below 2% following blood contact(*41*). Under these criteria, the EndoBot and its locomotion parameters demonstrated acceptable hemolytic behavior, with hemolysis rates below 0.1% under all tested conditions.

8.4 *Thrombogenicity with platelet activation study*

As detailed in Supplementary Text 6.3, EndoBot samples were retrieved upon completion of the second hemolysis study and immediately fixed in 4% paraformaldehyde for 48 hours to preserve their structural integrity. Following fixation, the samples were stained with PE Mouse Anti-Human CS62P antibody (BD Pharmingen™, USA) to detect platelet activation markers. The blood-contacting inner surfaces of the EndoBots were meticulously analyzed for platelet activation, adhesion, and aggregation—critical indicators of thrombogenicity(*37*). We used platelet P-selectin expression as a reliable marker to characterize platelet activation and aggregation(*42*) **(Fig. 5E)**. High-resolution fluorescence imaging of P-selectin proteins was conducted using an Olympus IX73 fluorescence microscope, providing detailed visualization of platelet activation and aggregation. Consistent with the hemolysis rate patterns, collagen-treated EndoBots exhibited significantly elevated levels of P-selectin-expressing platelets attached to the robot surface, forming denser and larger aggregates. This outcome aligns with the known properties of collagen I, which presents epitopes for platelet-binding and activation(*43*). In contrast, heparin-treated EndoBots displayed substantially lower levels of P-selectin-positive platelet adhesion and aggregation. Although untreated EndoBots induced slightly more platelet activation and aggregation than the heparin-treated variant, these levels remained closer to the heparin-treated group, suggesting potential thrombogenic potential over long-term exposure.

*8.5 Cytotoxicity*

To evaluate the potential cytotoxicity of EndoBot, we tested the impact of any leachable substances from its body adversely affect human umbilical vein endothelial cells (HUVECs). HUVECs were expanded in T-225 cell culture flasks containing trypsinization media. For the assay, the cells were collected and seeded into 96-well plates at a concentration of $2 \times 10^3$ cells/well in triplicates, with each well containing the EndoBot sample. In this study, EndoBots were fabricated with different NdFeB:PDMS ratios of 0.5 : 1, 1:1, 2:1 and 4:1; and different the base to crosslinker mass ratio of 10:1 and 5:1. Before experiment, EndoBot samples were cleaned with alcohol, phosphate-buffered saline (PBS) and then placed in the plasma cleaner for 30 mins. A negative control without the EndoBot sample was included for comparison. The cells were incubated at 37 °C in a humidified atmosphere of 5% $CO_2$ for up to 24 hours.

The cell viability after exposure to the different samples for 1 week was assessed using the Live/Dead Cell Imaging Kit (Invitrogen, Thermo Fisher Scientific, Inc.). Fluorescence microscopy images were captured using the Olympus IX73 fluorescence microscope. Live and dead cells were counted using the integrated hybrid cell count analysis module. For each sample, n = 4, ensuring sufficient statistical power to evaluate the cytotoxic effects accurately. Viability was quantified by calculating the percentage of live cells relative to the total cell count in each well, providing a comprehensive understanding of the cytotoxic effects of the EndoBot samples.

The cell viability was calculated according to the following equation:

$$\text{Cell viablitiy } (\%) = \frac{C_{sample} - C_{blank}}{C_{control} - C_{blank}} \times 100\% \quad (18)$$

Where $C_{sample}$ is absorbance of sample exposing with EndoBot, $C_{control}$ is absorbance of control (without exposing with EndoBot), and $C_{control}$ is absorbance of blank group.

The cytotoxicity results of the optimized EndoBot are shown in **Fig 5F**, and the cell viability data for various materials used in its fabrication are presented in **Table S8**. As shown in Figure 5E, compared to no-treatment cultures, HUVECs exposed to EndoBot-conditioned medium showed no significant changes in cell viability or morphology. Additionally, EndoBots with other material composition ratios also exhibited no adverse effects on cell viability, indicating their potential suitability for human applications.

**Table S8. Human umbilical vein endothelial cell viability after exposure to EndoBot-exposed culture medium for 1 week.**

| PDMS:Curing agent (weight ratio) | PDMS:NedFeB ratio (By weight) | Live cell count (per field of 5x field) | Percentage (%) |
|---|---|---|---|
| 10:1 | 1:0.5 | 2648 ±134 | 105.3% ±5.3% |
| 10:1 | 1:1 | 2535 ±220 | 100.8% ±8.8% |
| 10:1 | 1:2 | 2598 ±243 | 103.3% ±9.7% |
| 10:1 | 1:4 | 2633 ±151 | 104.7% ±6.0% |
| 5:1 | 1:0.5 | 2552 ±99 | 101.5% ±3.9% |
| 5:1 | 1:1 | 2602 ±199 | 103.5% ±7.9% |
| 5:1 | 1:2 | 2569 ±167 | 102.2% ±6.6% |
| 5:1 | 1:4 | 2567 ±176 | 102.1% ± 7.0% |
| No treatment | No treatment | 2515 ±256 | 100.0% ±10.9% |

**Supplementary Text 9: Fluoroscopic-guided steering, localization and tracking in vitro**

Developing safe and effective endovascular interventions with EndoBot requires necessitates methods for real-time imaging, localization, and tracking. Since the vasculature is inherently non-transparent for visible light, visualizing small-scale robots depends on imaging techniques routinely employed in clinical practice. Interventional fluoroscopy stands out for its ability to provide deep tissue penetration, high spatial resolution, and near real-time image acquisition. These capabilities make it a powerful tool for guiding small endovascular instruments, such as catheters.

To assess the X-ray visibility of our concept, we developed a real eX-MMPt platform for remote navigation control of EndoBot, as illustrated in **Fig 6Ai, ii**. The perfusion system setup with detail operation condition is shown in **Fig 6Aiii**. Since the magnet has a high contrast under X-ray fluoroscopy, which can obscure the reliable visibility of the EndoBot, the magnet's moving trajectory is set to avoid obscuring the EndoBot's image during operation. This platform is designed for telerobotic drug delivery to the local endothelium and vascular wall. It utilizes a clinical C-arm (OEC 3D C-arm from General Electric, Boston, MA) to capture fluoroscopic videos in both cinefluoroscopic (cine) and digital subtraction (DS) modes, providing real-time imaging of EndoBot as it navigates within the patient's blood vessels under magnetic manipulation. The open workspace of the C-arm enables the successful operation of the magnetic manipulation platform without compromising imaging quality or requiring significant re-design of the two parallel working systems. For reporting purposes, the DICOM image files from the C-arm were converted to .JPEG format and .MP4 using RadiAnt DICOM Viewer (version 2023.1, 64-bit). The C-arm was also utilized in both ex vivo and in vivo experiments.

To ensure the compatibility of EndoBot with clinical C-arm systems, we initially quantified its radiographic contrast relative to iohexol, an iodine-based medical contrast agent, and its endovascular formulation Omnipaque™ 350, as the clinical benchmark. Achieving sufficient contrast is critical for detecting the robot as well as for managing patient radiation exposure and its potential short- and long-term side effects(*44*). For the contrast assessment, we filled 1 mL standard syringes with varying compositions of EndoBot precursor material, i.e., different NdFeB:PDMS ratios, and compared their X-ray contrast to that of iohexol **(Fig. 6B)**. For fluoroscopy-guided interventions, a medical device should exhibit contrast levels at least equivalent to 30% iohexol to ensure adequate visibility. Grayscale analysis of the samples revealed that all tested EndoBot precursor compositions exceed the 30% iohexol threshold, indicating that EndoBot would be readily visible in a clinical setting. This superior X-ray visibility arises from the higher atomic number of $_{60}Nd$ compared to $_{53}I$, which increases X-ray attenuation and thus enhances detectability under C-arm. Notably, the refined EndoBot composition (NdFeB:PDMS = 4:1) surpassed even the contrast level of 100% iohexol, outperforming this clinical standard.

To demonstrate the magnetic manipulation of the EndoBot under fluoroscopic guidance, we applied magnetic actuation to the EndoBot within the C-labeled phantom **(Fig 6C, Supplementary movie 4)**. Under high blood flow rate of up to 75 mL/min, to maintain the controllability with EB $3.6^{10\%}$ the magnet is always rotated clockwise (|B|=~145 mT, $f_m$ = 30 Hz) to make a magnetic propulsion against blood flow direction. We continuously visualized EndoBot and its motion at 15 fps using cine imaging mode and 6 fps using DS mode **(Fig. 6C, S23, and Supplementary movie 4)**. In DS mode, EndoBot was not initially visible while remaining stationary due to the real-time frame subtraction process performed by the C-arm system. Once it started moving, it became clearly distinguishable against the blank background, leaving a white digital trace at its initial position. Although it is slower than the cine mode, DS imaging has unique advantageous for endovascular applications in the chest and head regions, where overlying bone structures can obscure vascular details (*45, 46*). Despite successful visualization of EndoBot and near real-time image acquisition during motion, both cine and DS modes of fluoroscopy presented significant limitations. First, they provided limited structural information within the robot workspace **(Fig. 6C)**. Without clear lumen visibility, localization, tracking and navigation of the robot using external magnetic interactions could be severely compromised, endangering the procedural safety. To address this, we performed fluoroscopic angiography, similar to established clinical procedures for guiding catheters and guidewires(*47*). By injecting iohexol contrast agent into the bloodstream, we achieved lumen visualization that improved proportionally with the agent’s final concentration in the blood **(Supplementary Movie 4)**.

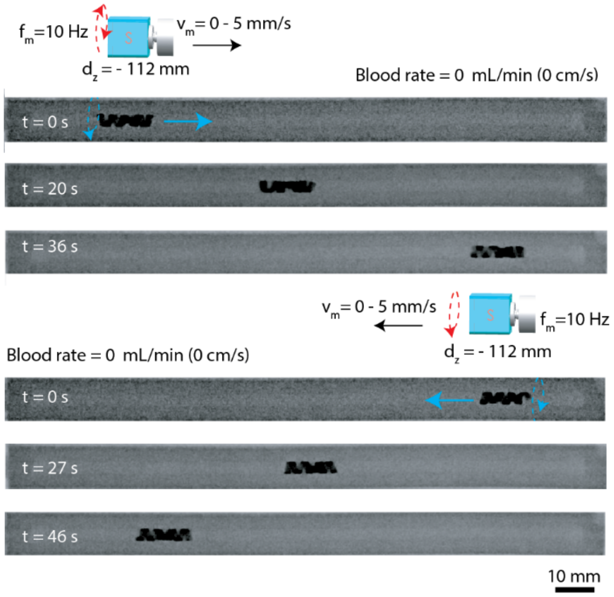

**Figure S23. Fluoroscopic-guided locomotion of $\mathbf{EB3.6_{10\%}^{10\%}}$ within blood-filled phantom vessel (without flow).** To move $EB3.6_{10\%}^{10\%}$ from left to right (0s-36s), the magnet is rotated counterclockwise and moved from left to right ($d_x$ = 0, 25 ≥ $d_y$ ≥0 mm, $d_z$ = -112 mm, $f_m$ = 10 Hz, $v_m$ = 0-5 mm/s) and while to move $EB3.6_{10\%}^{10\%}$ from right to left (0s-46s) the magnet is rotated clockwise and moved from right to left (operation condition: $d_x$ = 0, 0 ≥ $d_y$ ≥ -25 mm , $d_z$ = -112 mm, $f_m$ = 10 Hz, $v_m$ = 0-5 mm/s).

While fluoroscopic angiography is practical for localizing and tracking EndoBots and other untethered milli- and microrobots in vitro and ex vivo settings, adapting this method for in vivo use introduces additional challenges. Unlike catheters, which are tethered and thus carry minimal risk of escaping into the bloodstream, untethered devices such as EndoBot require continuous localization to ensure effective steering and prevent losing it to the bloodstream. Although periodic injections of contrast agents can assist localization in clinical procedures, continuous administration is neither safe nor feasible, particularly given the nephrotoxicity associated with systemic agent use and the potential for hypersensitivity reactions in some patients(*48*).

Therefore, to overcome these challenges, we recently developed an innovative virtual reality (VR)-based approach that integrates a digital twin of the robot's operational environment, a robot avatar and real time robot position data(*49*). In our system, we establish a connection between the real environment setup and the virtual environment setup using our custom interface. Each system performs its unique tasks and shares results with each other via the Robot Operating System (ROS) network, ensuring a flexible design and structure. **Figure S24** illustrates our data handling process among the various systems within this study and outlines potential future setups to improve our system for future applications. The virtual twins of the phantom channel were created by importing the .OBJ file of the phantom design (Fusion 360) into the Unity® 2022.3.0f1 editor version (Unity® Technologies). To achieve spatial alignment of the phantom channels and virtual twins, a calibration method was developed. Prior to the navigation task, an initial image was captured to (i) match the frame size of the input video to the virtual frame size and correct (ii) position and (iii) rotation of the object within the frame to the virtual twin. X-ray-opaque fiducials were used to identify the starting and ending positions of the phantoms defined on the image, followed by pixel-to-mm conversion.

As we did in our previous work (*49*), when the system started, a Python3 code will be called to set up all related data for initialization. These fiducials will be used to mark starting-ending positions of the phantom which we can define our starting point and the angles that our phantom has. Later, these data are used by Unity® to setup virtual environment by placing virtual twin to its correct coordinate and orientation which recently defined by Python3 code and fiducials. Data transfer between Python3 code and Unity® environment realized with a new version of the ROS2, more specifically ROS2 Humble. ROS2 was used to create a communication network for all separate system works within our project. Everything works within our local network and each system can send and retrieve data by publishing and subscribing to the related topics, data packages, on ROS2 network. In our work, we have python codes for object detection, C# codes for Unity® connection and simulation environment control and ROS2 environment that can communicate with all separate systems. For future uses, everything can be connected to the internet network for untethered communication between each system which can lead cross-continental connection. Before the spatial calibration, the connection of the Unity® system with the ROS2 network was established using ROS-TCP-Endpoint libraries from Unity® Technologies GitHub repository. With this connection, initial calibration data and future detected EndoBot data could be received by Unity® virtual environment, and the virtual twin position could be adjusted easily.

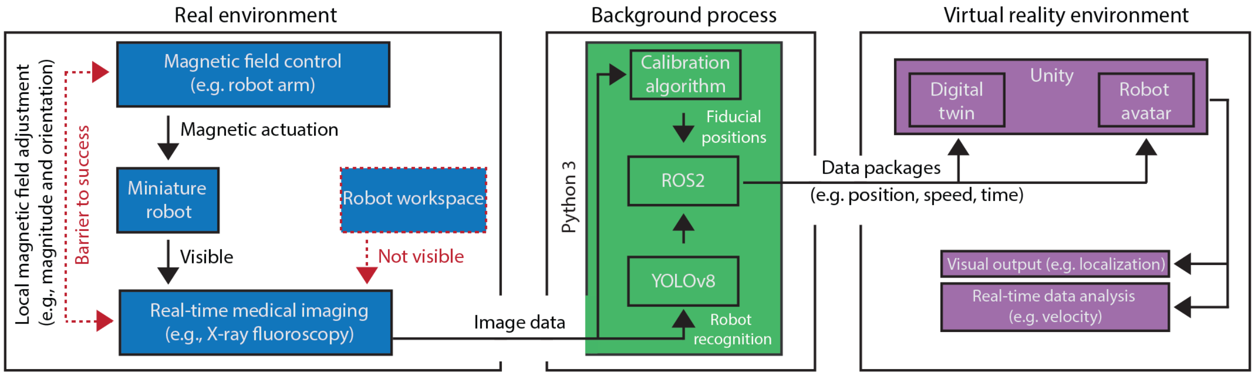

**Figure S24. The schematic diagram illustrating the data process and interface connections in the proposed virtual enhancement strategy.**

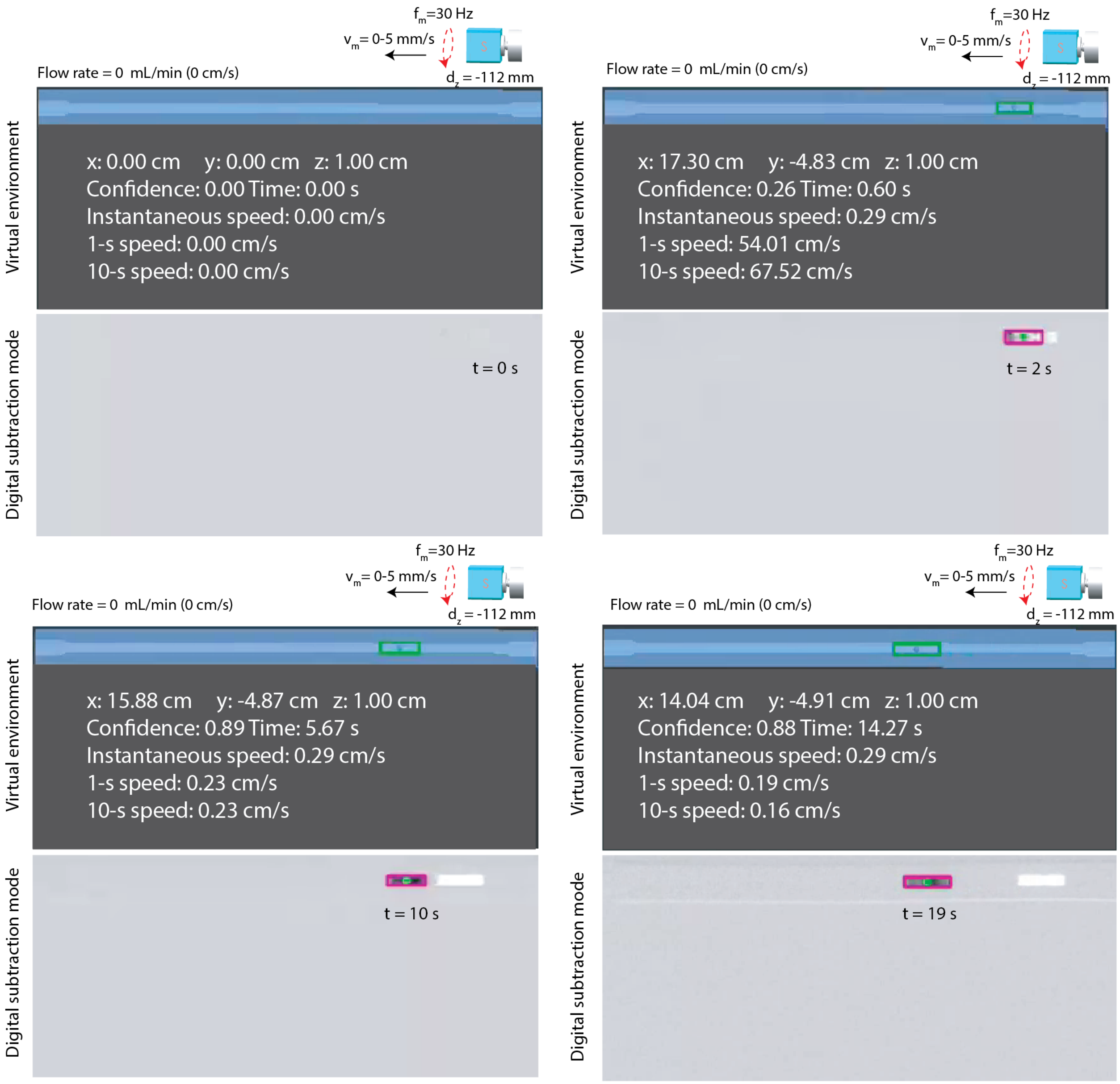

**Figure S25. X-ray fluoroscopy-guided real-time localization and telerobotic navigation control in the virtual twin interface.** During the digital subtraction mode, a series of X-ray images are continuously captured of the same area. The output image is produced by subtracting the previous image from the current one, which enhances robot detection and minimizes background noise. Despite the near-zero environmental information, the digital twin interface ensures safe and efficient navigation of EndoBot (operation condition: the magnet is rotated clockwise and moved from right to left, $d_x = 0$, $0 \geq d_y \geq -25$ mm, $d_z = -112$ mm, $f_m = 30$ Hz, $v_m = 0$-5 mm/s).

For EndoBot detection, same training model was used, which consist of 318 images of EndoBot. Roboflow platform (Roboflow, Inc, Des Moines, Iowa, USA) was used to create and annotate the database(*50*). Images that used to create this data base undergo to same process as our previous work which created 1210 final images. To apply object detection, a new version of the You Only Look Once (YOLOv8, Ultralytics, MD, USA) algorithm was used (*51*).

Subsequently, a Python script was developed to utilize the trained dataset for EndoBot detection on pre-recorded videos or real-time stream. With this EndoBot detection model, the center position of the EndoBot during movement was recorded, as shown in **Fig S19C**. The EndoBot speed in this research was calculated based on the highest slope, as defined in **Fig S19C**.

In the recreation of the twin EndoBot in Unity®, a detected EndoBot bounding box transferred into the virtual environment, along with its center point which marked with a circle in the background. For the visual validation of a detected EndoBot, this bounding box was also displayed in the recorded videos around the EndoBot, as shown in **Fig S25**, whenever a detection event occurred. To demonstrate the feasibility of real-time navigation of EndoBot under X-ray fluoroscopy-mediated detection in the virtual interface, $\mathrm{EB3.6}_{10\%}^{10\%}$ was moved within C-label PDMS-based vessel under X-ray. The virtual twin interface in DS mode results is shown in **Fig S25**, and both cine fluorography and DS modes are demonstrated in **Supplementary Movie 4**.

Fluoroscopic imaging inherently provides 2D projection data, lacking z-axis information essential for 3D tracking. To overcome this major caveat, we implemented an algorithm that infers z-axis positions by correlating 2D positional data with the pre-defined geometry of a virtual model (**Fig S26).** This process begins with high-resolution scanning of the target vessel anatomy using 3D imaging systems, such as X-ray or MRI. The resulting image data are segmented to generate a digital twin of the anatomical structure in a virtual environment. This segmentation data can also be used to fabricate 3D-printed models, enabling interventional or surgical mock-ups for preoperative study. After creating the digital twin, we synchronize the coordinate systems of the virtual and real environments. This synchronization involves the introduction of virtual "milestones," which are spherical markers with known 3D coordinates strategically placed manually along the central axis of the vessel **(Fig 6D** and **Fig S27)**. These milestones span the entire navigable region within the vessel, providing a comprehensive spatial framework for localization. Each milestone is sized at approximately one-quarter of the length of EndoBot (~3 mm), enabling precise segmentation of the robot's trajectory. Using this framework, the algorithm infers z-axis positions by mapping the robot's 2D projected location to the closest milestone in the virtual environment. By combining this inferred z-axis information with the x-y positional data from fluoroscopic imaging, we generated accurate 3D positional datasets **(Fig 6D** and **Fig. S27)**.

To validate the robustness of this approach, we created a digital twin of a human umbilical vein segmented from a fluoroscopic 3D scan (**Figs S26, S27, and Fig 6D**). Fabrication method and production steps for PDMS-based umbilical cord phantom vessel was mentioned detailed in **Fig S28.** The live human umbilical cord was collected from Abrazo Labor and Delivery Arrowhead Campus, Arizona. First, the umbilical vein was slowly perfused with lactated Ringer's injection to remove all residual blood and then perfused with vascupaint yellow material which includes 5 ml of diluent, 4 ml of vascupaint yellow silicone, and 0.45 ml of catalyst (Vascupaint Yellow Kit (Cat# MDL-121), MediLumine Inc, Canada). After the vascupaint yellow material dried and becomes solid. The umbilical cord with the vascupaint yellow material was scanned by OEC 3D C-arm (General Electric, Boston, MA) for X-ray projection image acquisition. The 3D umbilical phantom model was reconstructed from X-ray projection images by using 3D slicer software (*52*). A sacrificial mold for 3D umbilical phantom was 3D printed using a K1 Mx AI fast 3D printer (Creality, Shenzhen, China) using acrylonitrile butadiene styrene (ABS) filament (Creality). This mold is then sprayed with non-adhesive ease release 200 spray and placed in a container printed by the same 3D printer. Then, the container was filled with a mixture of SYLGARD™ 184 Silicone Elastomer Base and Curing Agent, combined at a 10:1 mass ratio. Following an overnight curing at room temperature, the sacrificial ABS mold was dissolved with acetone next day, resulting in the phantom in its final form. Similarly, before the experiment, the PDMS-based umbilical cord vessel was placed in a plasma cleaner for 5 minutes, cleaned with ethanol for 30 minutes, and then dried at 80°C for 30 minutes.

**Figure 6D** shown that the fluoroscopic stream did not show any detail of the vascular trajectory as a model $\mathrm{EB2.1}_{10-30\%}^{10\%}$ moves along the vessel. However, spatially calibrating the virtual and real environments allowed us to localize and track the motion of $\mathrm{EB2.1}_{10-30\%}^{10\%}$ along the phantom vessel.

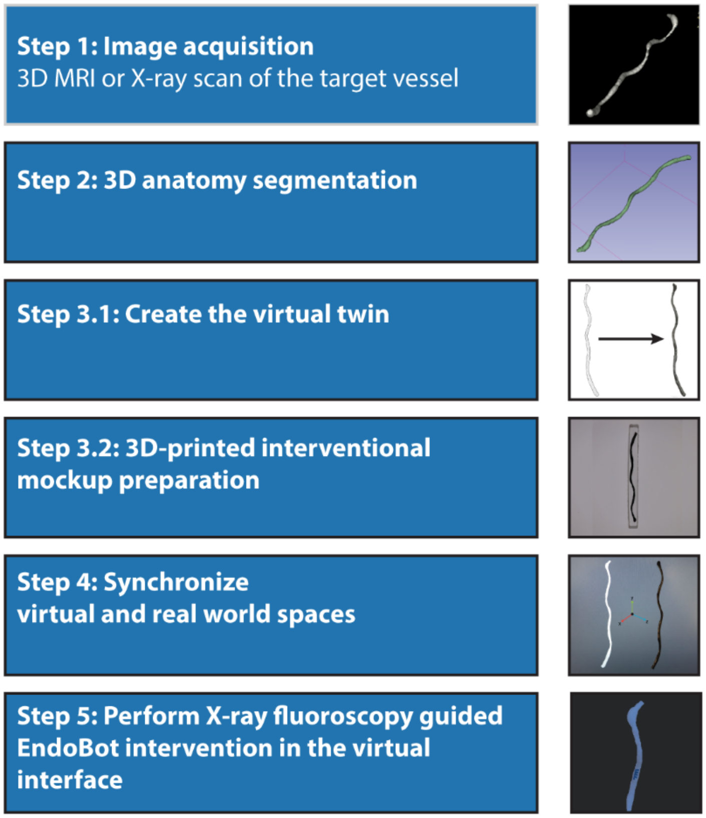

**Figure S26. Envisioned integration process.** Virtual anatomy enhancement is essential for real-time localization and precise remote navigation control of miniature robots within the human body with high precision and acceptable safety. To implement this strategy, a preoperative planning stage would be needed. This stage would begin with a high-resolution scanning of the target tissue or organ anatomy using 3D imaging systems like MRI or X-ray. Following this, the segmentation of the image data would be used to create a digital twin of the anatomical details in the virtual environment. The segmentation data could be further utilized to create 3D-printed models and provide an interventional or surgical mock-up for preoperative study. Once the coordinate systems of the virtual and real environments were synchronized, the EndoBot intervention would be safely performed with the visual tracking and localization feedback integrated with the virtual interface.

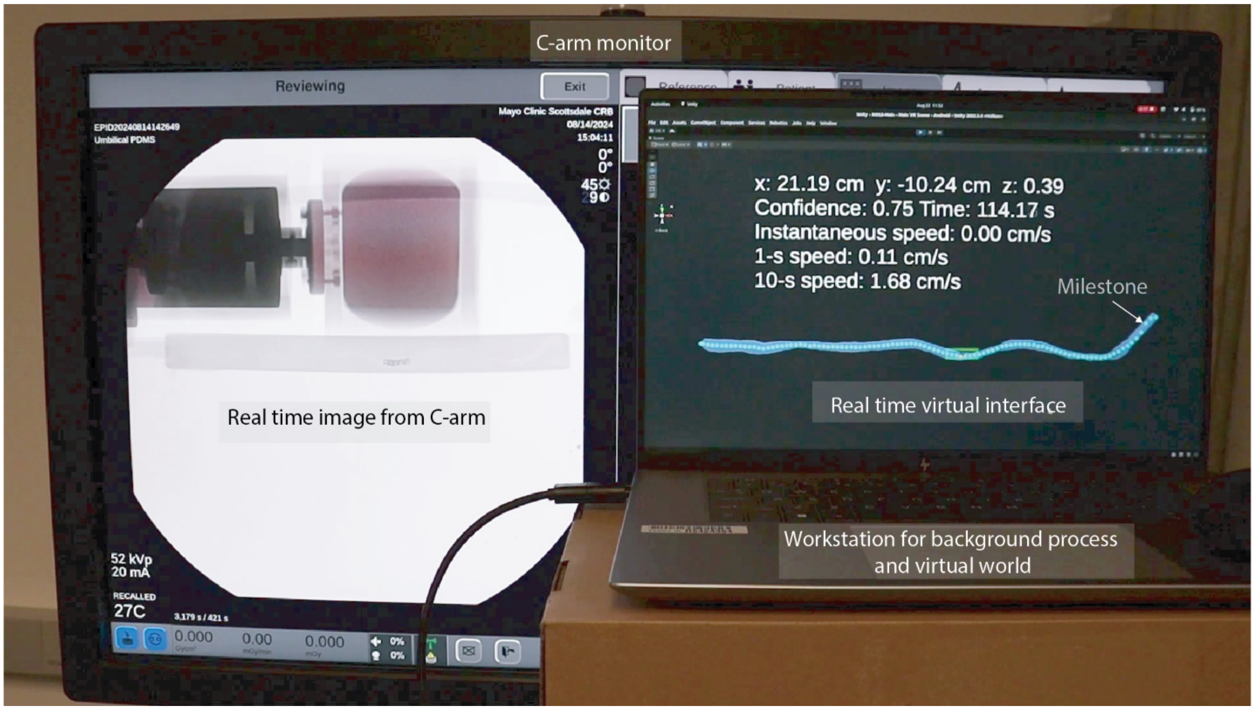

**Figure S27. Close-up view of the real-time system captured from the C-arm monitor and virtual interface, performing X-ray-guided EndoBot intervention for the validation of the approach in a quasi-3D vascular setting.**

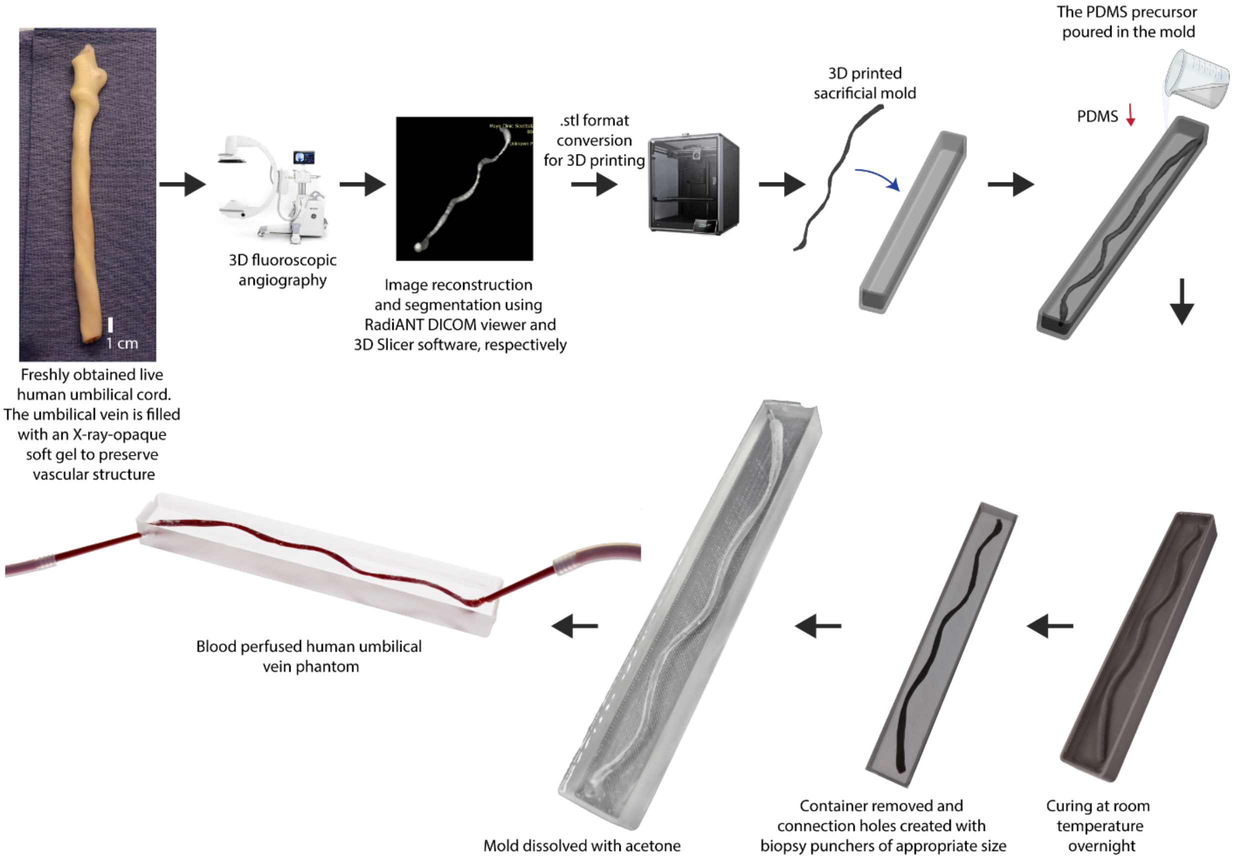

**Figure S28. Fabrication methods and production steps for PDMS-based umbilical cord phantom vessel for virtual interface.**

To clarify the functionality of our real-time matching system, we also define the delay mechanism inherent to the process. As illustrated in **Fig S29**, delays are introduced due to the frame rate of the input stream and the time required for image processing. Upon receiving a new frame ($F$) from the stream input, the detection process is initiated immediately. Under standard conditions, this detection process requires approximately 25 ms ($t_D$) to complete, while the frame rate is set at 15 fps, corresponding to an interval of 66 ms between frames. Consequently, the system is capable of processing the same frame up to 2.5 times within each interval, which introduces several challenges.

Processing the same frame multiple times unnecessarily increases computational workload without offering any additional benefit, and it may also result in erroneous or inconsistent data generation. To address these issues, we implemented a flexible waiting mechanism within the detection algorithm. Once the detection for the current frame ($D$) is completed, the system pauses for a defined waiting period ($t_W$) until the next frame is captured. This approach ensures that each frame is processed exactly once, optimizing system performance.

By enforcing a single processing instance per frame, this method produces the most reliable results and ensures that only one set of data is transmitted to other systems within the ROS2 network. This streamlined communication not only reduces computational overhead but also maintains the integrity and clarity of the data flow within the network.

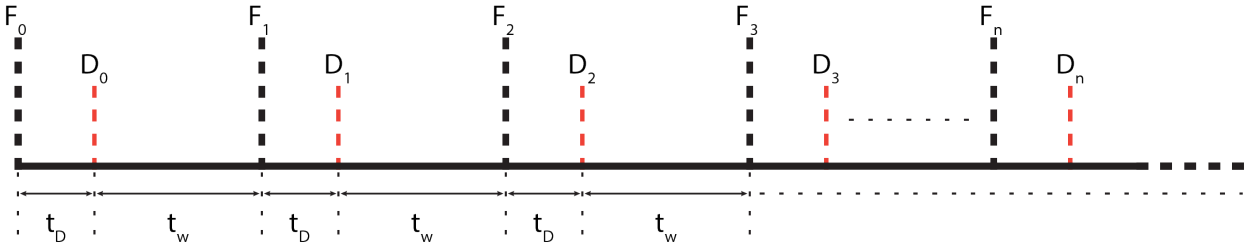

**Figure S29. Delay graph shows how we accomplish clean timing for image processing and data publishing.**

Real-time processing of visual data further allowed us to precisely determine and display average robot velocity. Even when the visibility of structural details diminished due to the passage of the robot arm or magnet through the imaging field, the EndoBot detection algorithm maintained reliable performance without interruptions or loss of localization precision. Similarly, our previous results demonstrated reliable tracking even when the visual signal is nearly indistinguishable from background noise(*49*). This highlights the significance of robot detection algorithms for ensuring the safe and effective navigation of untethered endovascular robots.

While this VR-enhanced method significantly reduces reliance on contrast agents for localizing and tracking EndoBot, future efforts should focus on extending the capability to 3D navigation in animal models. Although biplane fluoroscopy, which employs two X-ray sources could address the challenge of 3D localization limitation, it also significantly increases overall X-ray radiation exposure(*53*). Additionally, for magnetic robot control, biplane fluoroscopy imposes operational constraints on the external magnet and robotic arm, as X-ray beams are obstructed in at least one plane at any given time. Expanding the virtual twin concept offers a promising alternative, enabling precise 3D localization with a single X-ray source while minimizing radiation exposure and preserving the flexibility of robotic actuation.

### Supplementary Text 10: Atraumatic fluoroscopic navigation in perfused human umbilical veins ex vivo

To evaluate the mechanically adaptive, robust locomotion of EndoBot and its impact on the vessel wall integrity and ECs, we conducted navigation experiments using normothermically perfused human umbilical vein models ex vivo (**Fig 7A**). Fresh human umbilical cords were obtained from Abrazo Labor and Delivery Arrowhead Campus, Arizona, based on a material transfer agreement between Mayo Clinic Arizona and Abrazo Health. Fresh human umbilical cords were obtained immediately after cesarean sections, and re-perfusion was established by circulating heparinized whole cow blood (~37 °C) through the umbilical vein, following previously described protocols(*54*) (**Figs 7A, S30**).

Similar to the phantom vessels, perfused veins were completely invisible under the C-arm without contrast agents **(Fig S30)**. Thus, iohexol was infused into the bloodstream to enable uninterrupted visualization of the vessel throughout the experiment to determine the diameter of the umbilical vein (the diameter measurements, obtained via the RadiAnt DICOM Viewer) and used for magnet orientation during navigation. Based on the initial angiographic characterization of the lumen's minimum and maximum diameters across the target segment, we selected EB2.1 to ensure safe navigation by adhering to two fundamental design rules for EndoBot: maintaining continuous surface contact (Rule 1) and avoiding excessive deformation beyond allowable limits **(Fig 2A, B)**. To accommodate variations in vein size, we prepared a set of EndoBots with different diameters in advance, based on both literature data (*54*) and our design principles outlined in Fig. 2A. These design considerations ensured that EndoBot’s locomotion was mechanically adaptive and minimally invasive, safeguarding the structural and cellular integrity of the vessel during navigation.

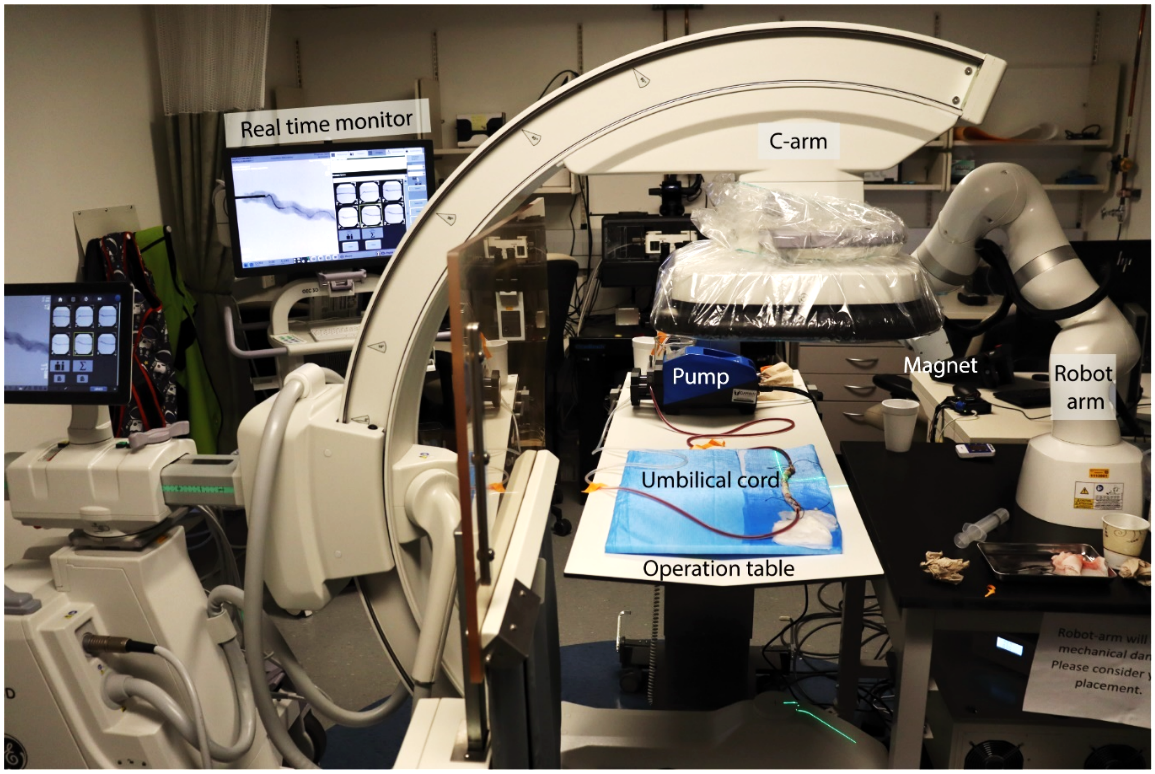

**Figure S30. Our X-ray fluoroscopy-guided magnetic microrobot manipulation setup during an angiography demonstration in a normothermic, blood-perfused, human live umbilical vein.** Initial fluoroscopic angiography is used to determine the vein's lumen size and uniformity. Based on this information, a suitable EndoBot is selected, ensuring that the anticipated robot deformation remains within the range of 10-35% to achieve robust endoluminal navigation. Additionally, based on the structural characteristics of the umbilical vein, the trajectory of the magnet was identified.

**Figure 7B** demonstrates the forward and backward motion of EndoBot against and with a blood flow rate of 10 mL/min (**Supplementary Movie 5**). Along the target vessel segment (Umbilical Vein 1), EndoBot underwent dynamic elastic deformations ranging from ~12% to 32% (the diameter of EB2.1 under locomotion was determined by using the RadiAnt DICOM Viewer **Figs S31**) conforming to the irregular and likely vasoconstricted lumen caused by surgical trauma. Frame-by-frame analysis of deformation patterns confirmed that EndoBot's ability to elastically adapt to these irregularities enhanced the robustness and safety of navigation while maintaining blood flow patency **(Fig 7B)**. This mechanically compliant design prevented excessive force on the vessel walls, reducing the risk of localized damage to the endothelial lining.

In a worst-case scenario experiment, conducted in Umbilical Vein 2, EndoBot was subjected to supraphysiological blood flow conditions (50 mL/min, >200 cm/s flow velocity at regular segments) and extreme vessel dilation **(Fig. 7C)**. Even under this condition, we maintained magnetic stability of EndoBot in the dilated segment without drifting, successfully moving it into the regular segments without losing helical structure and causing disruptions to the blood flow.

After four round trips of EndoBot within a ~60 mm vein segment (<5 min) under optimized actuation parameters (|**B**| = 100 mT, $f_m$ = 30 Hz), the targeted vein was collected for histological analysis. The segment was perfused with 10% neutrally buffered formalin solution for 5 minutes, followed by fixation in excess formalin for 24 hours at 4–5°C. Histological sample preparation followed standard protocols at the Mayo Clinic Histology Core Facility.

Briefly, fixed tissue sections were embedded in paraffin, cut into 2–3 μm slices, and mounted on microscope slides. The first group of slides was stained with Hematoxylin and Eosin (H&E), Masson's trichrome, and periodic acid-Schiff (PAS) to evaluate vessel wall integrity and endothelial lining. The second group underwent CD31/PECAM-1 immunohistochemistry staining (Invitrogen, MA5-18135) to confirm the presence of endothelial cells. H&E, Masson's trichrome, and PAS staining and CD31/PECAM-1 staining images were captured using Olympus EP50 and Olympus IX73 inverted fluorescence microscopes (Olympus, Tokyo, Japan).

The H&E, Masson's trichrome, and PAS staining results, presented in **Figs. 7Di, ii**, and **S32A**, depict the tissue morphology of Umbilical Veins 1 and 3, respectively. CD31/PECAM-1 staining images for the same samples, shown in **Figs. 7Diii** and **S32B**, reveal no evidence of endothelial cell denudation, demonstrating preserved vessel wall structure and single-cell endothelial lining, thereby confirming atraumatic navigation.

Each umbilical vein presented a distinct three-dimensional vascular structure, yet EndoBot demonstrated consistent and reproducible locomotion and safety outcomes across all tested models. For example, in Umbilical Vein 2, EndoBot experienced deformation ranging from 0% to 33% (**Fig. 7C, S34**), while in Umbilical Vein 3, deformations ranged from 3.2% to 27.8% (**Fig S35**). These results highlight the ability of EndoBot to safely adapt to varying vessel geometries, supporting its potential for use in anatomically diverse or pathological vasculature.

The ex vivo results emphasize the translational potential of EndoBot for safe and effective endovascular interventions. The mechanically adaptive locomotion ensures robust navigation in highly variable vessel geometries, such as those encountered in diseased vasculature with stenoses, dilations, or vasospasms. Furthermore, the preservation of vessel wall and EC integrity suggests that EndoBot could perform interventional tasks with reduced vascular trauma compared to traditional devices, such as catheters or stents, which often cause endothelial denudation or mechanical injury.

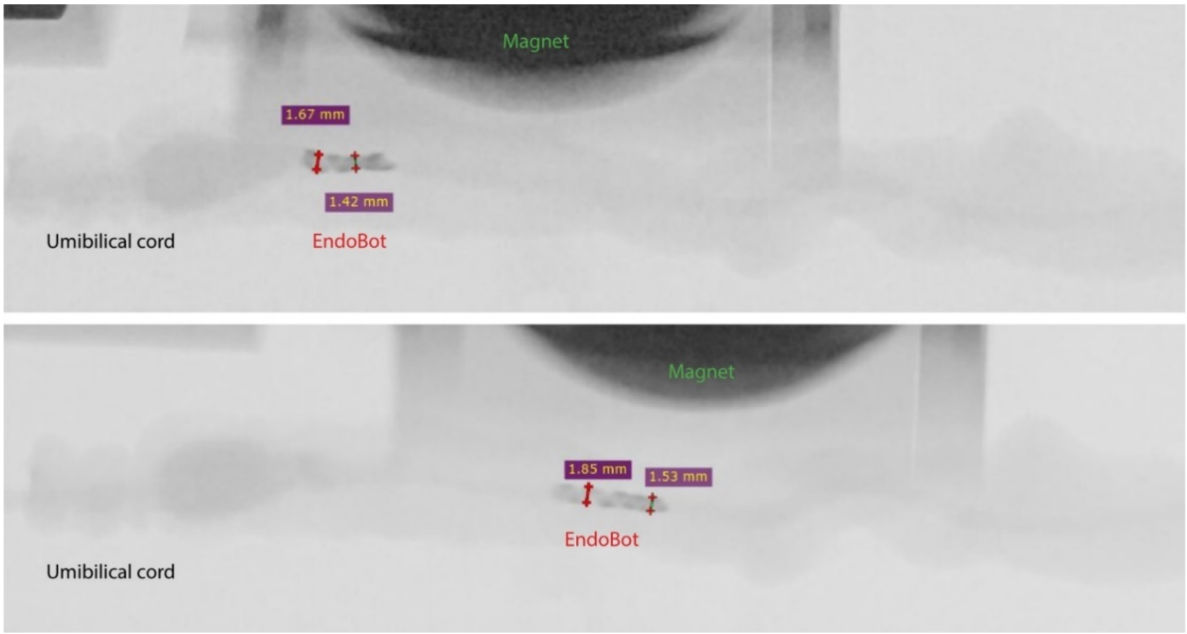

**Figure S31.** $\mathrm{EB2.1}_{12\%-32\%}^{15\%}$ **navigating normothermically perfused human Umbilical Vein 1 under real-time fluoroscopic guidance.**

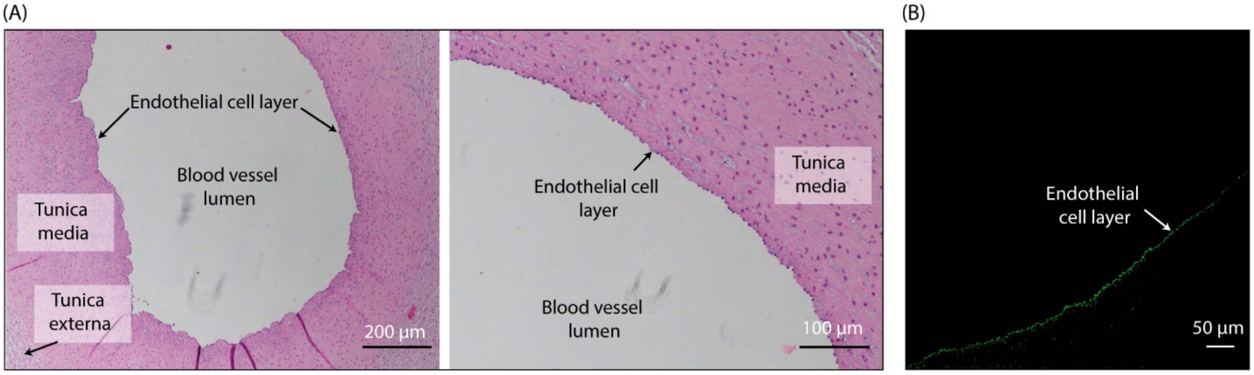

**Figure S32. Illustrative images from histological analysis revealing the physical impact on the endothelial cell layer and inner vessel wall for Umbilical Vein 3.** (A) Masson's trichrome, and periodic acid-Schiff (PAS) staining result. (B) CD31/PECAM-1 immunohistochemistry staining result.

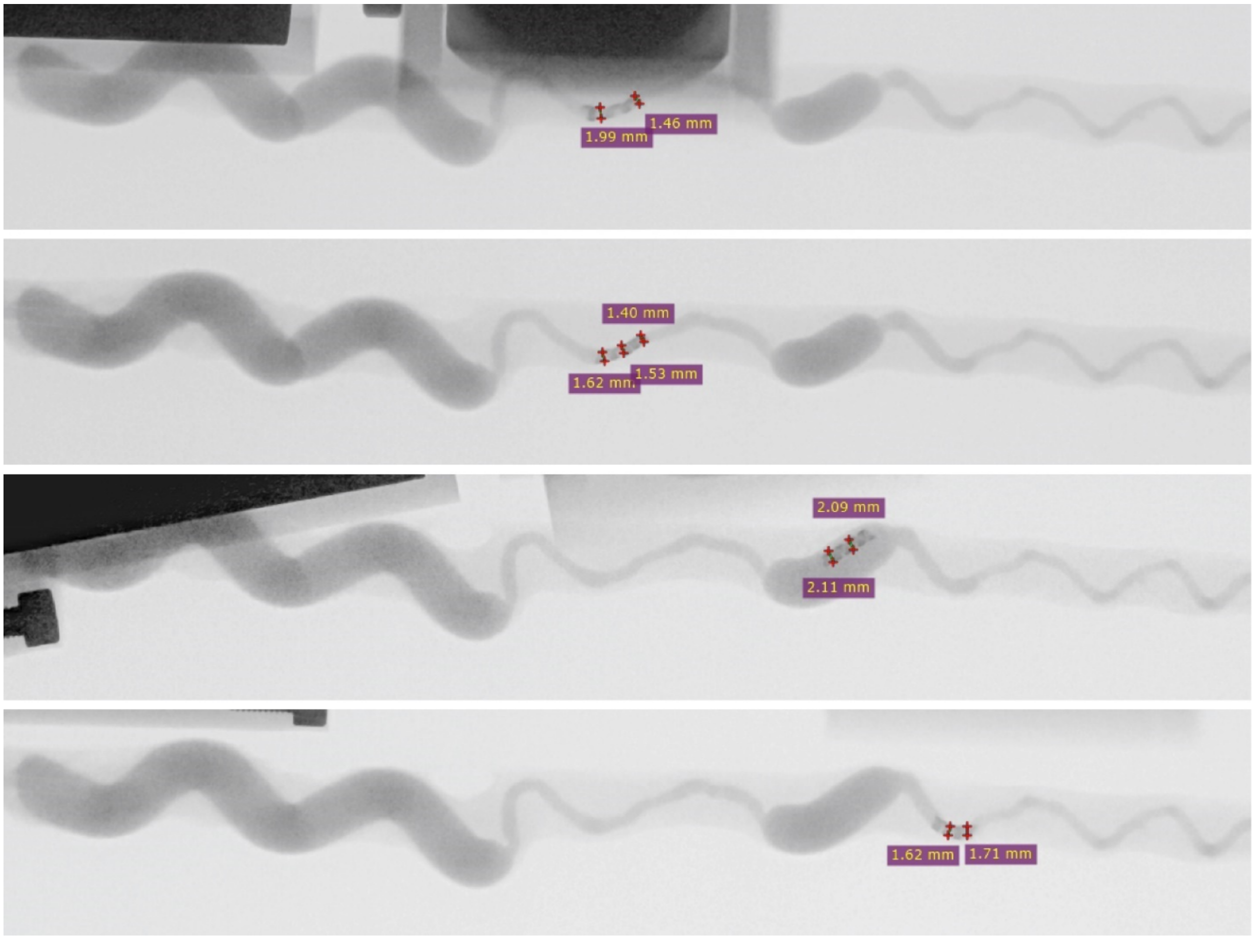

**Figure S33.** $\mathbf{EB2.1^{15\%}_{0-33.3\%}}$ **navigating normothermically perfused human Umbilical Vein 2 under real-time fluoroscopic guidance.**

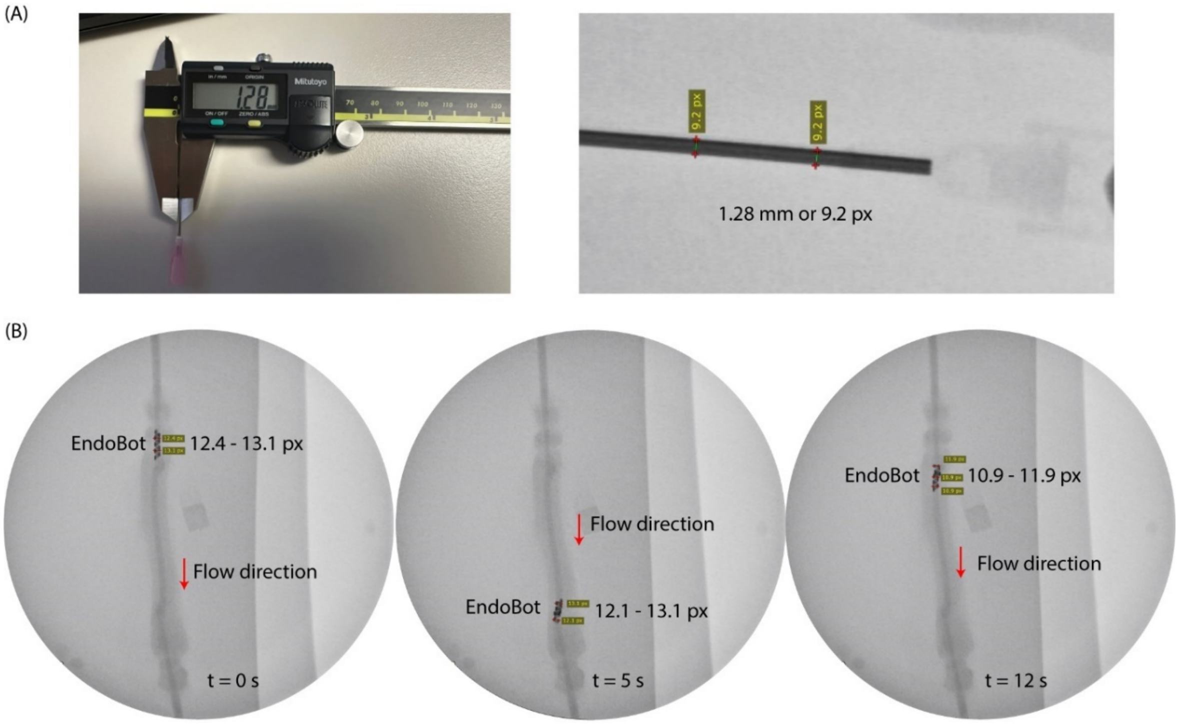

**Figure S34. Umbilical Vein 3 for $\mathbf{EB2.1}^{\mathbf{15\%}}_{\mathbf{3.2-27.8\%}}$ demonstration under real-time fluoroscopic guidance.** (A) Reference for measuring EndoBot diameter. (B) a series of umbilical X-ray images.

**Supplementary Text 11:** ***Endoluminal drug delivery***

The drug delivery mechanism of EndoBot relies on its ability to perform effective surface crawling along the vessel lumen, ensuring constant contact with the vessel surface. This design prevents blood flow from prematurely washing away the coating before it is fully deposited to the vessel wall. As EndoBot emerges from a vascular sheath or catheter and navigates through arteries or veins, its outer hydrophobic coating layer is gently transferred to the lumen via mechanical rubbing against the vessel surface **(Fig 8A)**. For a successful delivery, this process must form a stable, flow-resistant drug depot layer without causing downstream fragmentation.

To enable this capability, we formulated a transfer coating using acetyl tributyl citrate (ATBC), an FDA-approved pharmaceutical excipient(*55*). ATBC is a slightly hydrophobic, oil-like plasticizer that can protect pharmaceutical agents from rapid dissolution into the bloodstream(*55*). A similar coating strategy is employed in current DCB systems. For example, TransPax™ platform (Boston Scientific) utilizes a citrate ester excipient combined with low-dose paclitaxel (2 μg/mm$^2$) to adhere the therapeutic agent to the vessel surface during balloon inflation(*56*).

ATBC exhibits a blood half-life of approximately 0.5 and 5 hours in rats and humans, respectively due to serum esterase activity(*55*). It is metabolically cleared easily by rat and human liver microsomes rapidly with half-life less than 30 min. These properties make ATBC an excellent material foundation for developing a biodegradable, blood-compatible transfer coating tailored for localized drug delivery and sustained release on the endothelium and vessel wall.

To enhance the mechanical properties of the transfer coating for improved surface applicability and increased resistance to aqueous dissolution and flow-induced erosion, we additionally incorporated poly(methyl methacrylate) (PMMA), another hydrophobic, biocompatible polymer, into the formulation.

*11.1 Refinement of coating material*

Five coating formulations were prepared for mechanical refinement including:

**Sample 1:** 20 mg PMMA in 1 mL ATBC or 1.9% PMMA in ATBC (% w/w) and 6 mL acetone solvent.

**Sample 2:** 40 mg PMMA in 1 mL ATBC or 3.7 % PMMA in ATBC (% w/w) and 6 mL acetone solvent.

**Sample 3:** 80 mg PMMA + 1 mL ATBC or 7.1% PMMA in ATBC (% w/w) and 6 mL acetone solvent.

**Sample 4:** 160 mg PMMA + 1 mL ATBC or 13.2% PMMA in ATBC (% w/w) and 6 mL acetone solvent.

**Sample 5:** 160 mg PMMA without ATBC or Pure PMMA and 6 mL acetone solvent.

The Hei-PLATE Magnetic Stirrer (Heidolph Instruments, Germany) was employed to dissolve the components in the solvent. Following this, 200 µL of Rhodamine B, a fluorescent dye molecule, with a concentration of 2 mg/mL in acetone was added to the five above samples. Rhodamine B serves as a fluorescent reporter for imaging with the IVIS system and creates colors that are visible to the naked eye. After adding the Rhodamine B to the samples, the mixture was gradually transferred into a sample container with a radius of 25 mm and a thickness of 1 mm which was made by the FormLabs 3B+ 3D printer. Since the container's volume is much smaller than the total amount of the mixture, it was necessary to wait for the acetone solvent to fully evaporate after each addition before continuing to add more. Once the sample container was completely filled and no solvent remained, mechanical testing commenced. To evaluate the material's properties, viscosity tests were conducted by using a pointing brush to evaluate the material's properties. The drawing pen was dipped into the material to a depth of approximately 1 mm, then slowly withdrawn at a constant speed of 0.1 mm/s using the mechanical testing machine which used for EndoBot. The pointing brush was lifted to a final distance of 4 mm from the sample surface, and an image was captured at this endpoint as shown in (**Fig S35A, Supplementary Movies 6** for optimal formulation containing 3.7 wt% PMMA in ATBC).

From the captured images, it is evident that increasing the PMMA ratio in the coating formulation resulted in a more viscous material. Its high viscosity allowed the material to form cohesive and uniform threads, maintaining strong adhesion and creating a consistent structure. To demonstrate the applicability of brushing with Sample 2, a pointing brush was used to create a Rod of Asclepius medical symbol on a clean plastic surface, which was then imaged using the IVIS system, as shown in **Fig S35B**.

To evaluate the resistance to serum dissolution of the coating material, we prepared five samples, using the same amounts of material as mentioned above. A fluorescent reporter was also prepared to measure the amount of drug eluting from the coating. In this case, we used Fluorescein (FITC) instead of Rhodamine B due to its lower reactivity with ATBC, which helps improve the accuracy of the test. A 1 mg/mL FITC solution in acetone was prepared. We took 1 mL of this FITC solution in acetone and mixed it with the five samples. After thorough mixing, the acetone was allowed to evaporate completely before adding 10 mL of Bovine Serum (BS, LOT 2471439 from Innovative Research, Inc., USA) to each sample. At specific time intervals (e.g., 0 minutes, 15 minutes, 30 minutes, 45 minutes, and every hour until 12 hours), 200 µL of the mixture was taken from each sample and transferred into a 96-well plate for analysis (repeated n = 3). After each sampling, 200 µL of fresh Bovine Serum was added to each well to replenish the plasma and maintain a constant plasma volume. The 96-well plate was then measured using the

IVIS machine, and the FITC concentration was calculated using a calibration curve prepared beforehand from the same source of FITC solution, with concentrations ranging from 20 μg/mL to 0.097 μg/mL. Finally, the percentage of resistance to serum dissolution was calculated, as shown in **Fig S35C**. We observed the same trend in its resistance to serum dissolution with increased PMMA ratio.

To further access the flow-induced erosion stability, we conducted an experiment as shown in **Fig S35D**. A solution was prepared by mixing 160 mg of PMMA, 4 mL of ATBC (3.7% PMMA in ATBC, w/w), and 6 mL of acetone. From this solution, 500 μL was taken and mixed with 500 μL of a Rhodamine B solution at a concentration of 2 mg/mL in acetone to prepare the final solution. Next, 1 mL of the final solution was added to a glass container, and the acetone was allowed to evaporate completely. Once the acetone had evaporated, the EndoBot was ready for use. Another 1 mL solution containing only Rhodamine B at a concentration of 1 mg/mL in acetone was prepared and allowed to evaporate completely. Both samples, after acetone evaporation, were added to 18 mL of BS, and mixing was performed using the Scientific Industries SI-0236 Vortex Genie 2 Mixer (Scientific Industries, Inc., Bohemia, NY, USA) at maximum speed for 5 minutes. The BS was then removed from the samples. As shown in **Fig S35D**, the results confirm that our coating material remains stable under these conditions.

We can conclude that the optimal formulation, containing 3.7 wt% PMMA in ATBC, effectively balances surface applicability, resistance to dissolution, and stability against flow-induced erosion. The refined coating formulation readily wets the surface of EndoBot, forming a stable, lubricant and self-supporting outer layer **(Fig 8B)**.

To evaluate the impact of the coating layer on the mechanical properties of the EndoBot backbone, $EB3.6^{0\%}$ with and without coating were moved in a conical-shaped narrowing vessel phantom (A labeled vessel phantom) filled with whole blood under the standard magnetic actuation conditions **(Fig. S36)**. Both configurations exhibited a maximum magnetic propulsion mediated deformation around 42%, demonstrating that the presence of the thin coating does not impair the safety limits of the robot performance.

#### *11.2 Demonstration of the Endoluminal Drug Delivery Concept and Coating Stability Against Flow-Induced Erosion*

Before demonstrating the endoluminal drug delivery concept, an EndoBot with a coating layer must be prepared. To do this, a solution was first prepared by mixing 160 mg of PMMA, 4 mL of ATBC (3.7 wt% PMMA in ATBC), and 6 mL of acetone. From this solution, 120 μL was taken and mixed with 120 μL of a Rhodamine B solution at a concentration of 2 mg/mL in acetone to prepare a final solution. To coat the surface of the EndoBot, it was first flattened into a bar and glued to the surface of a container. Approximately 240 μL of the final solution was then added to the container, and the acetone was allowed to evaporate. Once the acetone had completely evaporated, the EndoBot was ready for use.

To demonstrate the endoluminal drug delivery concept, we employed the coated $\mathrm{EB3.6}_{10\%}^{10\%}$ for a ~60 mm long phantom vessel segment filled with bovine serum at a locomotion frequency of 10 Hz (**Fig 8C)**. The in vivo imaging system (IVIS) revealed that when $\mathrm{EB3.6}_{10\%}^{10\%}$ traverses the vessel segment a single pass (1x), the spatial distribution of the payload demonstrates a gradual decline in radiant efficiency along the length of the segment. This decline may result from coating depletion, where the device releases its payload as it moves along the segment, leaving less material available for deposition further along the vessel. Additionally, the single pass may provide less physical interaction or contact time with distal regions, leading to lower efficiency in coating those areas. However, increasing the number of passes (1x, 3x, 5x) within the segment enables a more uniform transfer of the coating, with higher retention of radiant efficiency at distal regions. This pattern indicates that the amount of delivery can be spatially modulated by adjusting the number of passes, achieving both spatial control and enhanced distribution uniformity along the vessel.

To assess the stability of the coating against flow-induced erosion, the vessel segment was recirculated with fresh serum at 10 mL/min for 30 minutes **(Fig. 8D)**. The washing step failed to remove the coating from the vessel walls, indicating that the coating is stable and flow-resistant. Surface stability of the coating was also validated against extreme vortex forces in a serum-filled environment **(Fig. 8D)**. These features demonstrate that once EndoBot is delivered to the lumen, it forms a stable, dissolution- and shear-resistant drug reservoir, creating a window for the drug to be transferred across the vessel wall locally.

*11.3 Fragmentation of the coating during and after the delivery*

Fragmentation of the coating during and after the delivery could impact downstream perfusion and lead to ischemic tissue injury. To evaluate coating fragmentation behavior, we placed 10 μm pore-size filters in the circulation line.

Prior to the first experiment, a section of the coated vessel was prepared ensuring the most uniform material distribution. The preparation followed this protocol: a solution was created by mixing 160 mg of PMMA, 4 mL of ATBC (maintaining a ratio of 3.7% PMMA in ATBC w/w), and 6 mL of acetone, resulting in a total volume of approximately 10 mL. From this solution, 240 μL was taken and mixed with 240 μL of a Rhodamine B solution at a concentration of 2 mg/mL in acetone.

Approximately 480 µL of the resulting final solution was applied to the surface of a 6 cm-long platinum-cured silicone tubing-based vessel phantom (with a lumen diameter of 3.2 mm). The acetone was then allowed to evaporate completely, leaving a uniform coating of the material. Finally, sections of the coated vessel with the most uniform material distribution were selected for the experiment.

The section of the coated silicone vessel was then placed in a closed-loop circulation system containing 15 mL of BS. A 10 µm syringe filter was integrated into the system to capture any coating fragments released during flow, and the system was operated at a flow velocity of 6–10 cm/s for 30 minutes. After the closed-loop circulation, the vessel underwent open-loop washing with about 150 mL of clean BS for 15 minutes to remove residual serum and coating fragments from its surface. This experiment was considered a positive coating (or “Coating +” as shown in **Fig 8E**). All BS containing free Rhodamine B and detached coating fragments from the circulation was collected to quantify the total amount of free Rhodamine B in the BS using IVIS imaging and a pre-established calibration curve from the same Rhodamine B solution source. The total amount of Rhodamine B captured on the filter was also measured using IVIS imaging to evaluate the extent of coating fragmentation during circulation.

To perform a comparative analysis, a control solution was prepared by mixing the calculated amount of Rhodamine B, equivalent to the amount released during the previous experimental setup with the second coated silicone vessel, with 15 mL of BS. This control solution was circulated through the similar system, using a new 10 µm filter, for 30 minutes. Following this, the circulation system was washed with 150 mL of clean serum in an open-loop configuration for 15 minutes. IVIS imaging was also then performed on the filter to determine the total amount of Rhodamine B on the filter and assess the level of coating fragmentation during circulation. This experiment was considered a negative coating (or “Coating –” as shown in **Fig 8E**).

The filters from the positive and negative coating experiments, along with an unused filter for baseline fluorescence, were analyzed to compare the amounts of free Rhodamine B and detached coating material captured during circulation. The results show that after 30 minutes of closed-circuit circulation, only 2.1% of the circulating coating material was retained in the filter. Considering that free Rhodamine B exhibits ~0.6% non-specific adsorption to the filter, approximately 98.5% of the coating material passing into circulation was smaller than 10 µm, confirming the safety of this delivery procedure with minimal risk of downstream perfusion obstruction.

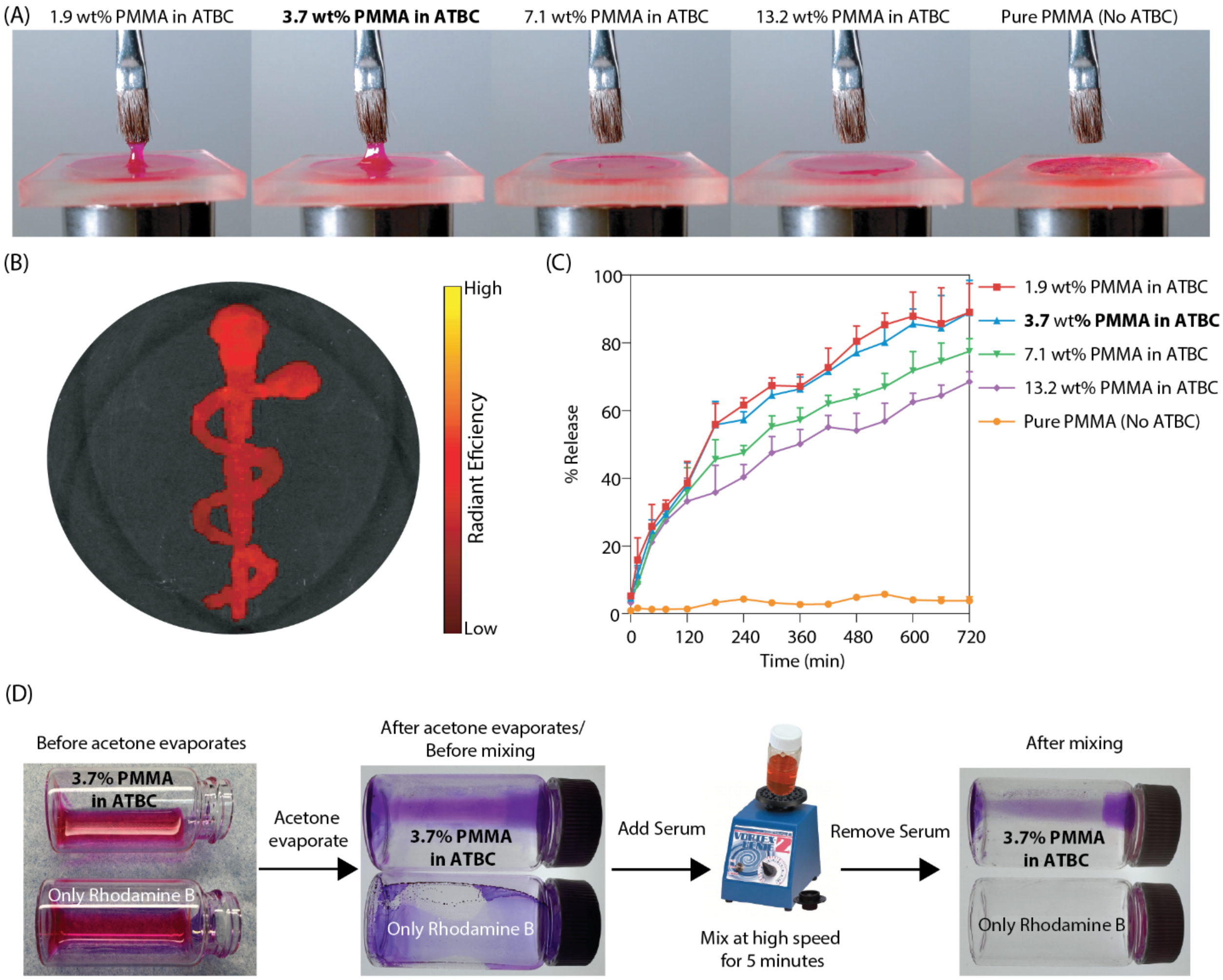

**Figure S35. Refinement and Evaluation of Coating Material for EndoBot-Mediated Drug Transfer Layer.** (A) Mechanical refinement of coating material. (B) Practical demonstration: the Rod of Asclepius symbol created using the optimized coating material, showcasing its precision and stability in forming drug coatings. (C) Resistance to serum dissolution of the coating material. (D) Flow-induced erosion stability at high mixing speeds.

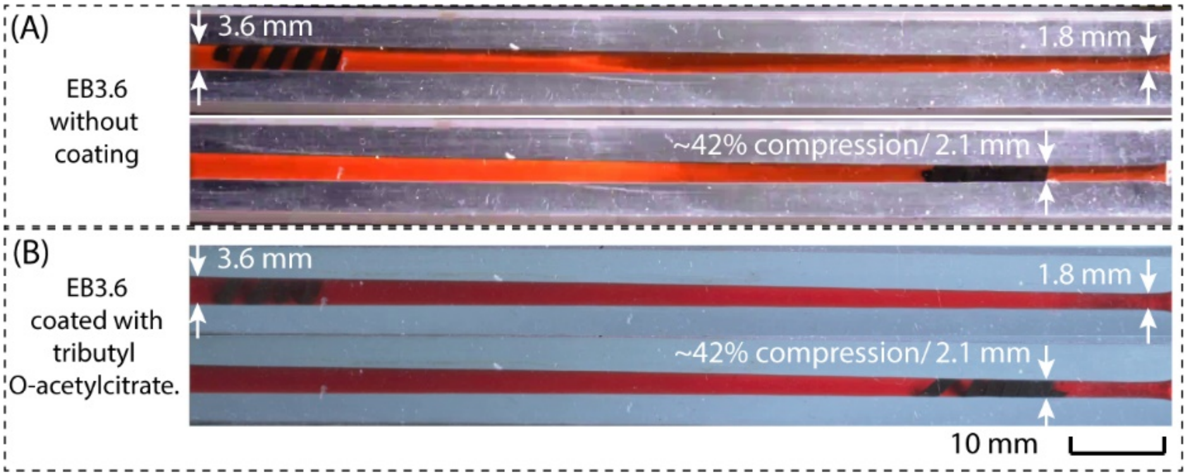

**Figure S36. Surface coating does not impact its mechanical properties, as demonstrated by the maximum deformability under magnetic propulsion.**

### Supplementary Text 12: Fluoroscopic navigation and drug delivery in live rat inferior vena cava

Prior to the experiment, a range of sheath catheters (4F to 7F) compatible with EndoBot sizes ranging from 1.4 mm to 3.8 mm were prepared. The selection of EndoBot size and catheter/sheath was based on the measured IVC diameter to ensure compliance with safety limits during operation within both the catheter/sheath and the IVC. An example of this preparation is shown in **Fig S37**. Prior to deploying the catheter/sheath into the IVC, the shape of the EndoBot was inspected to ensure that its helicity was maintained as shown **Fig S37B**. Note that EndoBot was cleaned with the same protocol as in **Supplementary Text 8.2.**

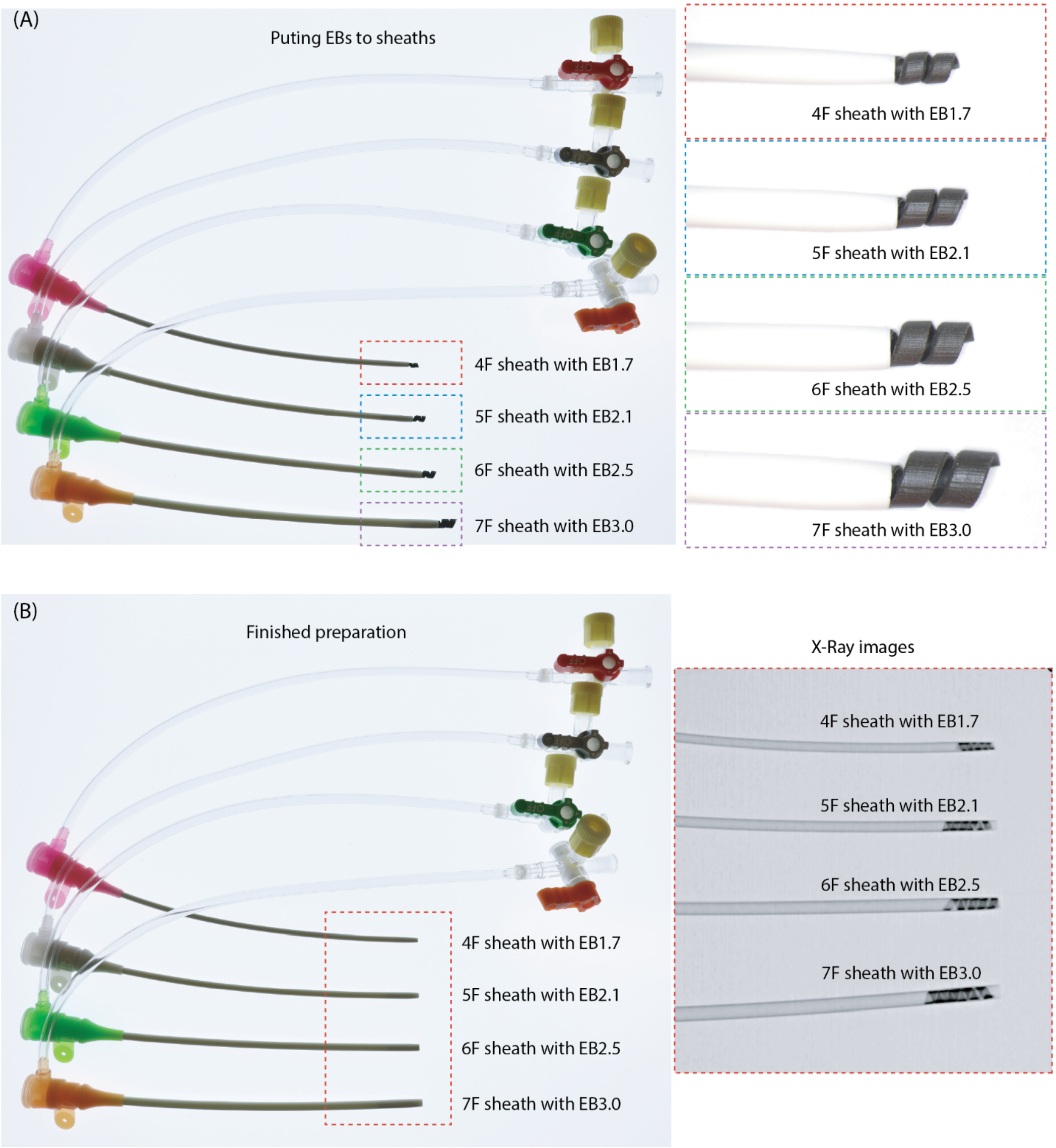

**Figure S37. Preparation catheter and EBs in vivo experiment.** (A) Deploying EBs into catheters. (B) Finished preparation by checking X-ray images to ensure helicity of EBs before deploying the sheath into targeted vessel.

To evaluate the safety and performance of EndoBot for navigation and drug delivery in a live animal model, we selected the inferior vena cava (IVC) of Sprague Dawley (SD) rats as the ultimate testbed. The mean peak flow velocity in the ~300g SD rat IVC ranges from 6–10 cm/s(*57*).

Previous ex vivo results with EB2.1 **(Fig 7)** and in vitro tests with EB3.6 **(Fig 3)** demonstrated successful navigation at flow velocities up to 100 cm/s in comparable sized vessels. Based on these findings, we hypothesized that EndoBot would safely perform in the normal rat IVC.

The rats (Sprague Dawley) used in this study were obtained from Envigo (Indianapolis, IN, USA). All rats were male, weighing between 300 – 375 grams. All experimental procedures involving animals were conducted in accordance with the guidelines outlined in the Application Format for Ethical Approval for Research Involving Animals and were approved by the Institutional Animal Care and Use Committee (IACUC-A00007480-24) of Mayo Clinic Scottsdale, Arizona, USA. Every effort was made to minimize animal suffering. This study used a total of five rats. EndoBots were prepared without coating for rats 1, 2,3 while prepared with coating for rats 4 and 5.

For each experiment, the rat was initially anesthetized with isoflurane (set to 3% with an oxygen flow rate of 0.75 L/min) to facilitate an intraperitoneal (IP) injection. Following this, the rat was anesthetized with pentobarbital at a concentration of 50 mg/mL (20 mL vial, 50 mg/kg, administered intraperitoneally). To alleviate pain before surgery, the rat received buprenorphine (0.3 mg/mL ampoule, 0.1 mg/kg, subcutaneously). Once the rat's abdomen was opened to locate the IVC for sheath/catheter insertion, anesthesia was maintained with pentobarbital at a concentration of 50 mg/mL (20 mL vial, 50 mg/kg, administered intraperitoneally) at a dose of 10–20 mg/kg every 50 minutes or whenever the rat showed signs of waking up. After the IVC was identified, a precise caliper was used to measure its diameter. For all rats in our experiment, a 6F PINNACLE® introducer sheaths (Terumo) and EB2.1 were selected.

To minimize thrombosis in the catheter/sheath used for the EndoBot, the sheath/catheter was prefilled with heparin (1000 IU/mL). During and after successful deployment of the catheter/ sheath into the IVC (the access position for the catheter/ sheath in the IVC is shown in **Figure 9A**(**i**), and a close-up view post-catheterization showing the insertion site on the rat's body is shown in **Fig 9A(ii)**), additional heparin was applied to the abdominal area to reduce thrombosis caused by blood leakage from the IVC into the abdomen. After the catheter/sheath was successfully deployed into the IVC, the rat was transferred to the working area under the C-arm and robot arm for the locomotion step, as shown in **Fig 9A(iii)**.

For all experiments, the initial approach for deploying the EndoBot into the IVC involved using a magnetic field (|**B**| = ~100 mT and $f_m$ = ~ 10-30 Hz) with magnetic propulsion aligned in the target movement direction. Once the EndoBot reached the end of the catheter/ sheath, which was in contact with the IVC, a second approach was employed using a contrast enhancer injection. Flushing the contrast enhancer not only facilitated the successful deployment of the EndoBot into the IVC but also cleared the IVC structure, preparing it for the subsequent locomotion step, as shown in Figure 9B and **Supplementary Movies 7 and 8**. Note that, to maintain the controllability for the EndoBot under high blood flow velocity in the IVC and the high fluid flow rate caused by flushing the contrast enhancer, the magnetic propulsion was reversed to counteract both the blood flow and the fluid flow from the syringe and the magnet was brought to close the IVC as much as possible to increase the magnetic radial force.

As shown in **Movies 7 and 8** (rat #2 and rat #3) and Fig. 9B (rat #3), EB2.1 not only successfully navigated the inferior vena cava (IVC) once but also repeated this process multiple times while the rats were still alive, demonstrating the repeatability of our concept in in vivo conditions. Similar to the ex vivo experiments, we determined the diameter of EB2.1 during locomotion using the RadiAnt DICOM Viewer. The results indicated that EB2.1 experienced compression ranging from 0% to ~31% while moving through the IVC. This observation aligns with our design rules, ensuring that EB2.1 maintained consistent contact with the lumen wall. After successfully navigating the EndoBot, the rats were euthanized using Euthasol at a concentration of 390 mg/mL (100 mL vial) with a dose of 450–550 mg/kg.

During deployment, we performed fluoroscopic angiography to visualize the IVC structure. Although we achieved success in four rats, the angiographic visualization was short-lived (<2 seconds) due to injection limitations in live rats. Consequently, we moved EB2.1 without direct visualization of the vessel line. Additionally, magnetic propulsion was not reversed during the second approach for deployment. As a result, for the first rat, we failed to move back the robot from the diaphragm, and it continued moving past the diaphragm into heart irreversibly (**Supplementary Movie 9**). These experiments highlight both the capabilities and limitations of EndoBot for precise navigation and drug delivery in live vascular environments under realistic physiological conditions.

Several segments of the IVC where EB2.1 motion were isolated for histological analysis. The histological sample preparation was conducted following the similar protocol as shown in ex-vivo experiment (**Supplementary Text 8).** Briefly, the segment of the IVC was first fixed in 10% formalin solution for 24 hours in a fridge (4-5 °C). Then, the fixed tissue sections were embedded in paraffin, cut into 2-3 μm thick sections, and then placed on microscope slides. The first group of slides was stained with Hematoxylin and Eosin (H&E), Masson’s trichrome, and periodic acid-Schiff (PAS) to evaluate vessel wall integrity and endothelial lining. H&E, Masson’s trichrome, and PAS staining images were captured using Olympus EP50 and Olympus IX73 inverted fluorescence microscopes.

The H&E, Masson's trichrome for rats #3, presented in **Figs 9C. The** results demonstrate that our method does not pose any safety concerns for the endothelial cells.

To prove the transferability of our coating material, after successful EndoBot locomotion, segments of the IVC from rat #4 and #5, where EndoBot motion was observed, were isolated for IVIS imaging. Additionally, one segment without EndoBot motion was also isolated from rat #4 as a control sample. The IVIS imaging results are presented in **Fig 9F**. The results demonstrated that EndoBot successfully transferred the coating material onto the IVC surface without causing damage or affecting the endothelial cells.
